# SATB2 and circ3915 RNA chromatin dysregulation drive *KRAS*-like oncogenic transformation

**DOI:** 10.1101/2024.10.04.616681

**Authors:** Rebekah Eleazer, Smitha George, Luke Shoemaker, Wesley N. Saintilnord, Kin Lau, Darrell P. Chandler, Yvonne Fondufe-Mittendorf

## Abstract

Even though epigenetic factors contribute to oncogenesis, most human cancer models still assume that disease originates from driver DNA mutations. Thus, it is still unclear if non-genetic mechanisms are sufficient to trigger malignant transformation. Special AT-rich binding protein 2 (SATB2) is a chromatin organizer that brings distal DNA elements into close proximity, thus remodeling chromatin structures to reprogram cell-specific and/or developmentally-sensitive gene networks. Here, we discover that *SATB2* generates a co-expressed *circ3915* RNA that is translated into a peptide and co-locates with SATB2 in the cell nucleus. Ectopic SATB2 or circ3915 over- expression rearranges global chromatin accessibility, generates *KRAS*- and *NFE2L2*-like oncogenic gene expression patterns, and transforms lung epithelial cells independent of driver mutations. Thus, oncogenic pathways can be activated in mammalian cells without pre-disposing mutations in oncogenes or epigenetic regulators.

**Graphical abstract:** 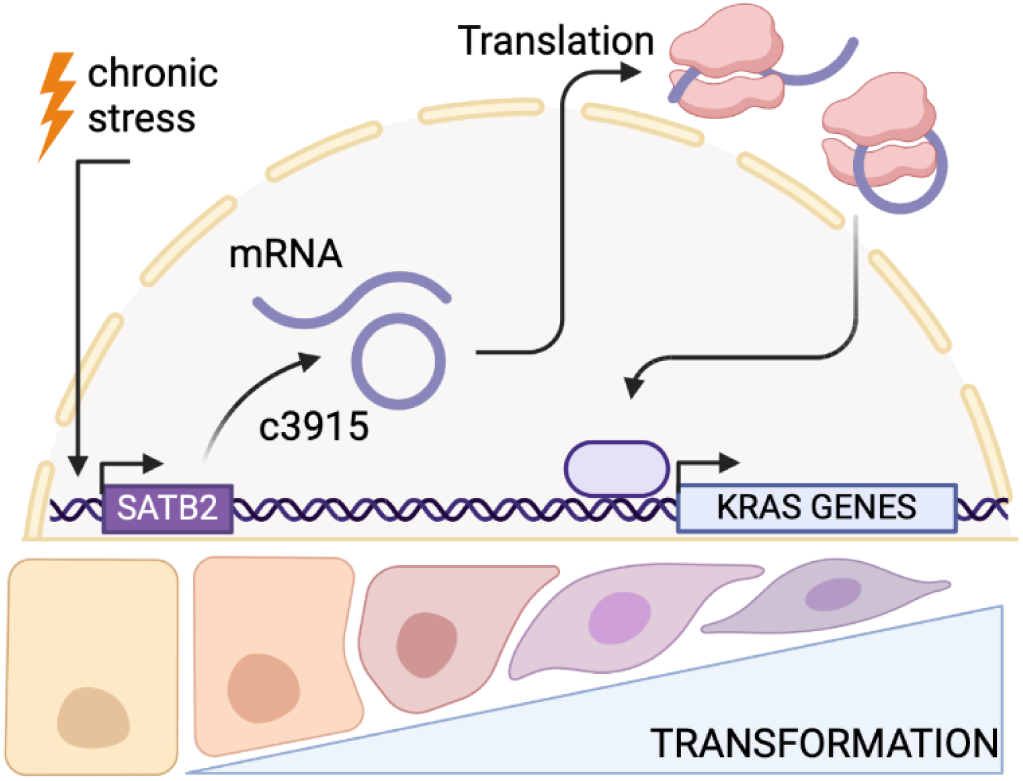

## INTRODUCTION

Cancer is usually viewed as a genetic disease arising from DNA mutations^1,2^, but we now appreciate that cancer is also caused by DNA and histone modifications that alter gene expression patterns^3–5^. Indeed, cancer genomes exhibit substantial changes across multiple layers of epigenetic regulation^6^, and can be heavily influenced by environmental factors^7^. Chromatin and epigenetic alterations implies that cancer can arise independent of DNA mutations, yet most studies into epigenetic dysregulation and oncogenesis still focus on DNA mutations in the factors that regulate histone post-translational modifications, DNA methylation, microRNA expression, or genome folding^8^. And while a recent study showed that transient, Polycomb-mediated silencing can trigger an irreversible switch to a cancer cell fate in *Drosophila*^9^, it is still unclear if non-genetic mechanisms are themselves sufficient to trigger malignant transformation in mammalian systems.

Three-dimensional (3D) chromatin structure is a non-genetic mechanism that can regulate gene expression^10,11^ and oncogenesis^12–14^ and is often related to histone post-translational modifications. 3D chromatin structures are often influenced by environmental factors, as is the case for the CCCTC-binding factor (CTCF)^15^ and its thousands of CTCF-binding sites^16–19^. For example, inorganic arsenic (iAs) is a ubiquitous environmental contaminant^20–23^ that selectively inhibits CTCF binding and generates new 3D chromatin structures that drive (onco)gene expression patterns^24^. iAs exposure also causes other oncogenic epigenetic changes^25–29^ that can occur in genome-wide and gene-specific manners^24,26,30^, thus altering chromatin accessibility at cis-regulatory elements^31–34^. In some respects, then, iAs-exposure and related toxicology models provide novel mechanistic insights into purely epigenetic regulation, chromatin structures, and oncogenesis that might otherwise be overlooked in conventional, mutation-associated cancer models.

Special AT-rich binding protein 2 (SATB2) is a cell-type-specific chromatin organizer that operates independently of and in cooperation with CTCF^35^. One way that SATB2 regulates gene expression^36–38^ is by binding to the nuclear matrix and regulatory DNA elements, thus bringing distal DNA elements into close proximity^36,39^. These SATB2-dependent, 3D chromatin structures are docking sites for chromatin remodeling complexes^40–42^, which then reprogram cell-specific and/or developmentally-sensitive gene networks in response to external cues^43–47^. *SATB2* mRNA is normally silenced in most adult tissues by microRNAs (miRNAs), and dysregulated *SATB2* mRNA expression and/or translation correlates with various cancers^48^ (including iAs- induced transformation^49,50^). In some cancers, *SATB2*-targeted miRNAs are themselves regulated by a circular RNA that is generated from the *SATB2* gene (circSATB2; circBase ID 0008928)^51–54^. Thus, SATB2 expression, regulation, and function involves multiple layers of regulation, and disrupting any one of these layers can alter gene expression patterns to promote oncogenesis.

Here, we identify a *SATB2* circular RNA (*circ3915*) that originates from exons 3-6 in the *SATB2* gene and is co- expressed with *SATB2* mRNA upon iAs exposure. SATB2 and circ3915 over-expression changes chromatin accessibility, elicits *KRAS-* and *NFE2L2*-like gene expression patterns, and promotes the oncogenic transformation of lung epithelial cells independent of any *KRAS* mutations. These data clearly indicate that oncogenic pathways can be activated in mammalian cells without pre-disposing germline or somatic mutations in oncogenes or epigenetic regulators, and suggest that *SATB2* and/or *circ3915* expression levels may be diagnostic biomarkers for active oncogenic pathways in lung (or other) cancers where there is no identifiable driver mutation. These findings also imply that cancer diagnostics targeting DNA mutations (instead of oncogenic gene expression pathways) will miss an important subset of cancer patients who might otherwise benefit from cancer therapies developed for patients with altered *KRAS* pathways.

## RESULTS

### Inorganic arsenic dysregulates oncogenic pathways

It is well known that chronic, low-dose iAs exposure (e.g., via drinking water) causes lung cancer^55^. We, therefore, exposed BEAS-2B human bronchial epithelial cells to 2 µM iAs for 12 weeks, an established low-dose (but rapid) exposure model that faithfully recapitulates the epithelial to mesenchymal transition (EMT)^29^. BEAS-2B cells harboring an activating *KRAS*-G12V mutation were used as a positive control for lung cell transformation and were a generous gift from the Brainson lab^56^. As expected^29,57^ and consistent with other reports^58,59^, chronic 2 µM iAs exposure transformed BEAS-2B cells into a more proliferative and migratory state (**Fig 1A, B**). We identified 8412 differentially expressed genes in iAs- transformed cells (DEGs; **Fig 1C** and **Fig S1A-B**), and validated selected genes using the Nanostring nCounter system (**Fig S1C**). Some of the most dysregulated genes (**Table S1, Fig S1B**) included oncogenes like *MMP1*^60^ and the *GAS1* tumor suppressor^61^, and oxidative stress response genes like *HMOX1*^62^. We were particularly intrigued by *SATB2* up-regulation, since SATB2 is a master regulator of chromatin accessibility^63^. While *SATB2* over-expression is associated with certain cancers^50^, the molecular mechanism for SATB2-mediated carcinogenesis is unknown. Molecular signatures database (MSigDB), KEGG, and Reactome pathway analysis also showed dysregulation of oncogenic pathways, including cell-substrate junction organization, axonogenesis, Wnt signaling, Hippo signaling, TGF-β, NCAM1 interactions, and KRAS signaling (**Fig S1D-G**). *KRAS* dysregulation was also identified in a C6 oncogenic signature analysis (**Fig 1D**). Thus, iAs-exposure dysregulates known oncogenic genes and pathways, perhaps as a consequence of *SATB2* dysregulation.

**Fig 1.**
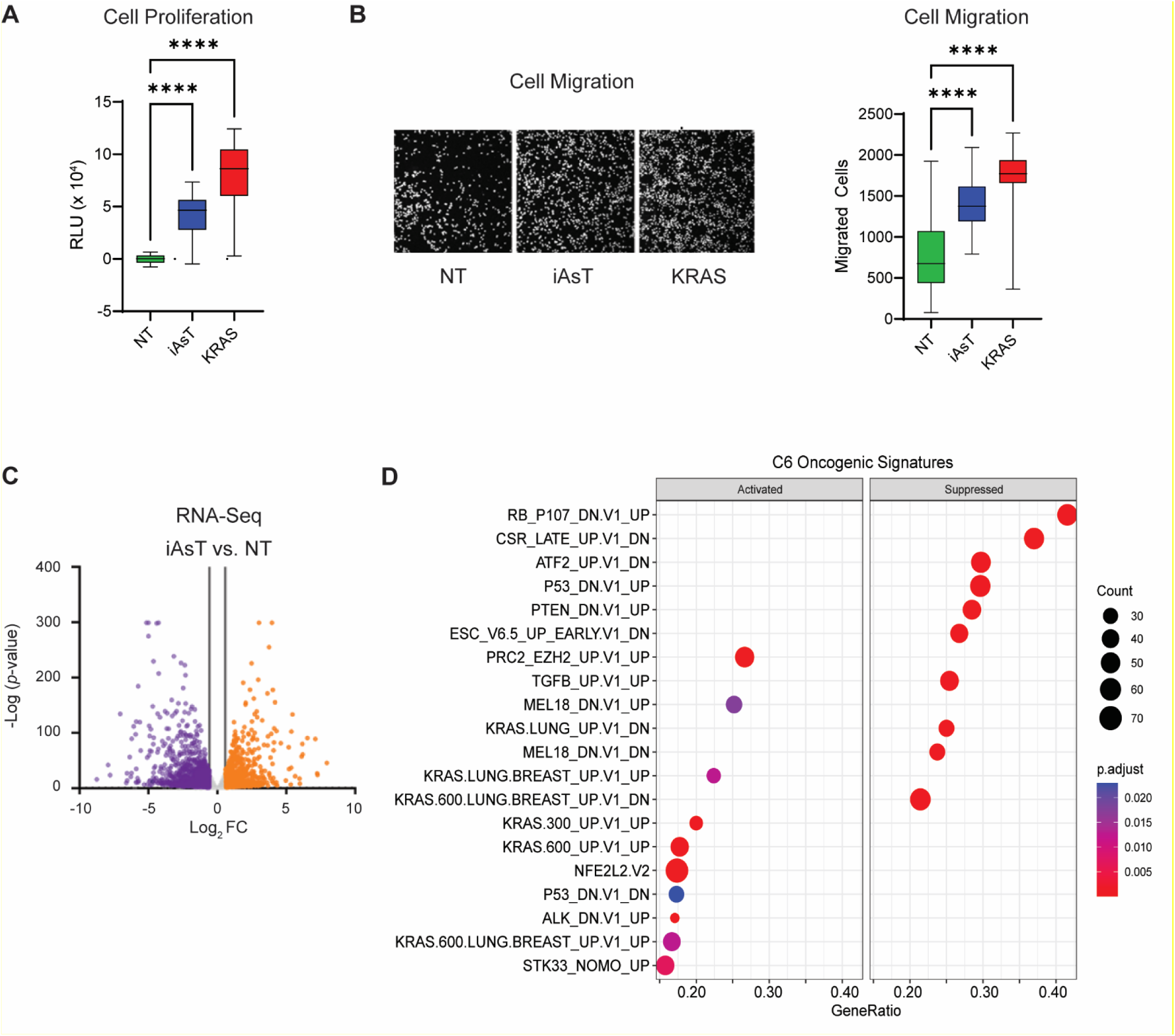
Inorganic arsenic (iAs) promotes cell transformation by driving oncogenic gene expression programs. Data were generated from the rapid (12 week) exposure model. **A**, Proliferation of non-treated (NT), iAs-transformed (iAsT), and KRAS/p53 mutant BEAS-2B cells 72 hr after seeding. Luminescence was measured with a CellTiter-Glo ATP assay. Values were normalized to background, and calculated relative to NT cells. N = 3 independent experiments with 6 technical replicates each; **** *p* <0.0001. **B**, Representative images (top) of migrating NT, iAsT, and *KRAS/p53* mutant BEAS-2B cells after 24 hr in a transwell assay. Cells were fixed and stained with DAPI, and cell counts were obtained using an Olympus Cellsens counting module. Cell numbers (bottom) were normalized to NT control cells. N = 3 independent experiments with 3 technical replicates each; \*\*\*\**p* <0.0001. **C**, Differentially expressed genes (DEGs) in iAs-transformed BEAS-2B cells. Purple dots represent down-regulated genes with a *p_adj_* <0.05, FC <-1.5; orange dots represent up-regulated genes with a *p_adj_* <0.05, FC>1.5. Grey dots represent genes with FC<1.5,>-1.5. **D**, Top enriched MSigDB C6 oncogenic signatures in the iAsT DEGs.

Since *SATB2* (a chromatin organizer) was one of the most dysregulated genes in iAs-transformed cells, we used ATAC-seq to identify differentially accessible regions (DARs), and correlate chromatin state (open/closed) with regulatory loci, nearby genes, and gene expression patterns. We identified ∼39400 DARs (**Fig 2A**; **Table S2**; **Fig S2A,B**), with iAs-transformed cells gaining chromatin access at promoters and losing access at intronic regions (**Fig 2B,C**; **Fig S2C**). iAs-transformed cells gained chromatin access primarily at ENCODE distal enhancer-like signatures while losing chromatin access at promoter-like signatures and proximal enhancer-like signatures (**Fig 2D**). Interestingly, some genes were associated with both open and closed chromatin regions (**Fig S2D**), suggesting differential regulatory mechanisms. When we associated DARs to nearby genes (1 Kb), we again identified known oncogenes and signatures, including those related to cell-substrate junction, ERBB signaling, focal adhesion, Hippo signaling, Wnt signaling, P13K/AKT signaling, the epithelial-to-mesenchymal transition, and KRAS/EGFR pathways (**Fig S2E-I**). Open chromatin was also enriched in KRAS transcription factor binding sites (e.g., JUNB and ERG), and NFE2L2, RUNX, and ZBTB3 binding sites. Closed chromatin associated with FRA1, NFY, SP1, ELK1, and JDP2 transcription factor binding sites and the CCAAT/enhancer binding protein (C/EBP) family of the basic region-leucine zipper (bZIP) transcription factors (**Fig 2E; Table S3**). Finally, there was some overlap between opened and closed DARs and increased or decreased gene expression (**Fig 2F**). In particular, open chromatin was enriched for up-regulated genes involved in the STK33_NOMO_UP, PRC2_EZH2_UP.V1_UP, and KRAS.50_UP.V1_UP, while closed chromatin was enriched for down-regulated genes involved in the ESC_J1_UP_LATE.V1_UP, and RAF_UP.V1_DN pathways (**Fig 2G**). Thus, iAs exposure significantly reorganizes chromatin structure around regulatory loci, transcription factor binding sites, and known oncogenes/pathways.

**Fig 2.**
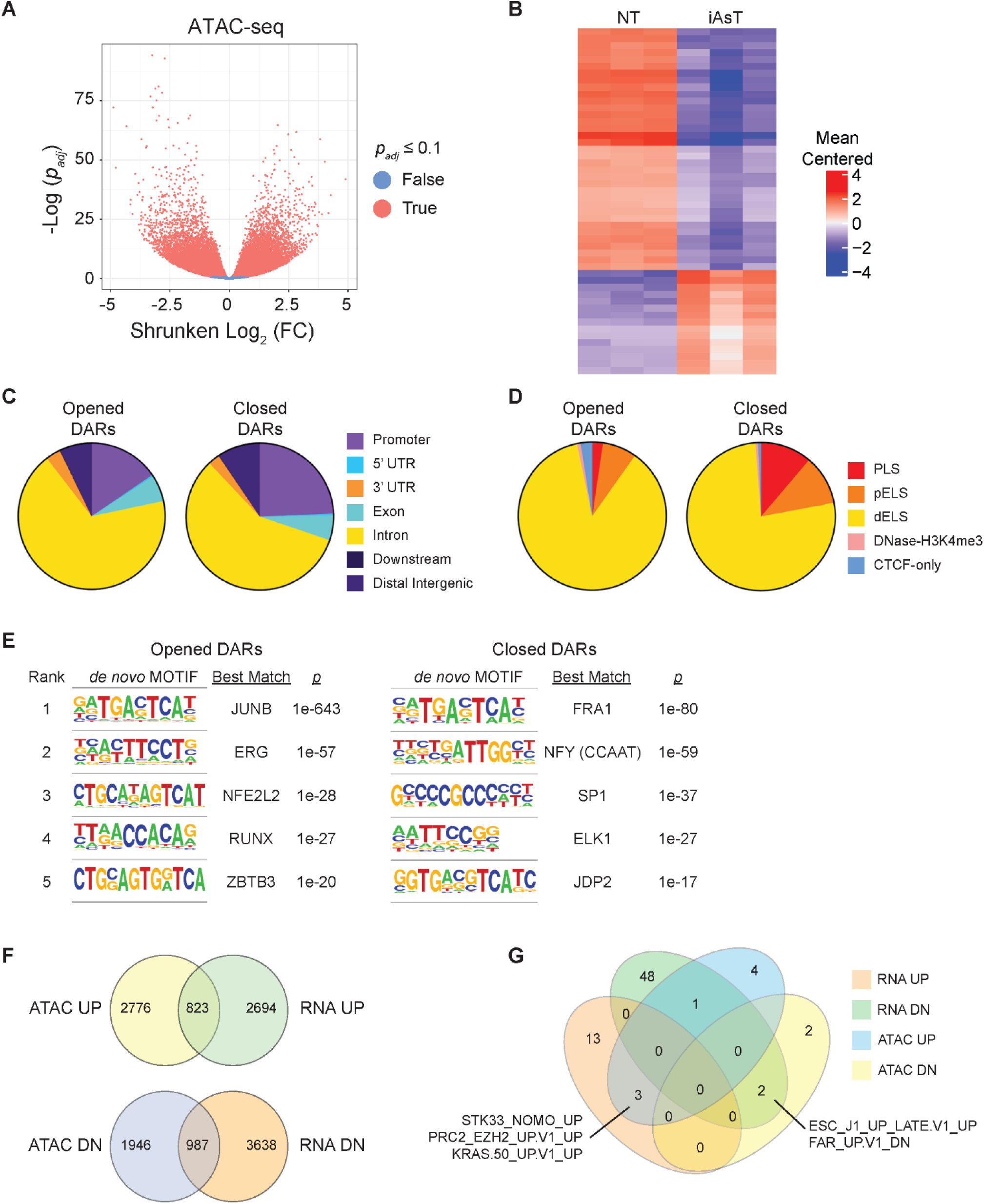
Inorganic arsenic (iAs) reorganizes chromatin structure around known oncogenes/pathways. Arsenic-transformed cells were generated with the rapid (12 week) transformation model. **A**, Volcano plot of differentially accessible regions (DARs) by ATAC seq between non-transformed (NT) and iAs-transformed (iAsT) BEAS-2B cells. DARs were defined by a *p_adj_* ≤0.1. **B**, Heatmap of the top 50 DARs between non- transformed (NT) and iAs-transformed (iAsT) cells. Rows are the top genes; columns are the mean centered peak values for each gene in each sample. **C**, Genomic distribution of all DARs in NT and iAsT cells. DARs regions were defined by *p_adj_* <0.01 and Shrunken Log_2_(FC) >1 for opened DARs, and ShrunkenLog_2_(FC) <-1 for closed DARs. **D**, Proportion of DARs in NT and iAsT cells that are annotated as ENCODE candidate Cis-Regulatory Elements (cCREs). PLS = promoter-like signatures; pELS = proximal enhancer-like signatures; dELS = distal enhancer-like sequences. **E**, Top enriched *de novo* transcription factor motifs and best matching transcription factors in iAsT opened and closed DARs. **F**, Congruence between iAsT DARs and proximal gene expression. ATAC UP = opened chromatin; ATAC DN = closed chromatin; RNA UP = up-regulated transcripts; RNA DN = down-regulated transcripts. Counts are the number of individual genes above the minimum fold-change threshold for mRNA expression (>1 Log_2_(FC)) or open chromatin (>1 ShrunkenLog_2_(FC)). **G**, MSigDB oncogenic signatures in iAs-transformed cells that were commonly enriched in DARs and DEGs. ATAC UP = opened chromatin; ATAC DN = closed chromatin; RNA UP = up-regulated transcripts; RNA DN = down-regulated transcripts.

### *SATB2* generates a circular RNA transcript

iAs exposure can lead to alternative mRNA splicing^29^, and circular RNAs are one possible product of alternate mRNA splicing that can play crucial roles in cell differentiation^64,65^. We therefore used the Arraystar Human Circular RNA Array to screen iAs-transformed cells for differentially expressed circRNAs (**Fig S3A,B**), and discovered several dysregulated, cancer-associated circRNAs (e.g., hsa_circ_0002490, hsa_circ_0002874, hsa_circ_0075320, and hsa_circ_0008928)^51,66–68^. Since circRNA microarray data are subject to false positive signals^69,70^, we used a custom Nanostring analysis to verify the most dysregulated circRNAs identified by microarray. One of the most up-regulated circRNAs from both experiments was *circ3915* (**Fig 3A; Fig S3A**), an uncharacterized circRNA that encompasses *SATB2* exons 3-6 (**Fig 3A**, right panel). *SATB2* mRNA and *circ3915* were co-expressed during iAs-induced transformation (**Fig 3B**), with increased *SATB2* expression correlating with increased SATB2 protein production (**Fig 3C**). Both transcripts are produced in an iAs dose- (**Fig 3D**) and time-dependent manner (**Fig 3E**), beginning with the initial iAs exposure and persisting throughout the transformation process. These findings were also replicated in iAs-exposed 16HBE cells (**Fig S3C,D**).

**Fig 3.**
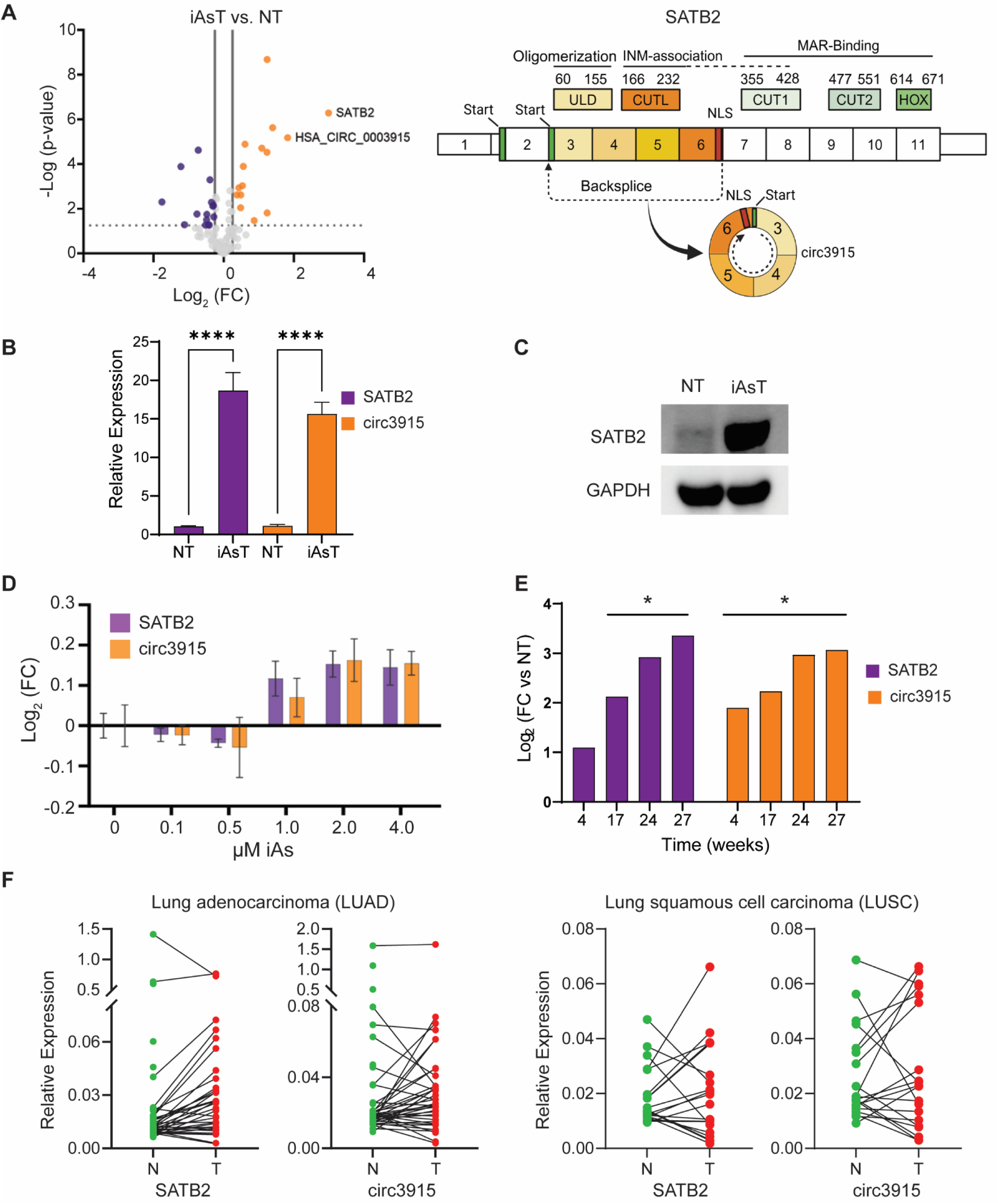
*SATB2* and circ3915 are up-regulated in iAs-transformed cells and lung cancer tumors. Data were generated in the rapid (12 week) iAs transformation model. **A**, Nanostring-based detection of differential RNA and circRNA expression in iAs-transformed (iAsT) BEAS-2B cells relative to non-transformed (NT) cells (left panel). Up-regulated RNAs (orange dots) were defined by *p* <0.05, FC >1.2. Down-regulated circRNAs (purple dots) were defined by *p* <0.05, FC <-1.2. Grey dots represent RNAs with no significant difference in RNA expression or below 1.5 magnitude fold change. Right panel shows *SATB2* and *circ3915* transcripts and corresponding peptide domains. ULD = ubiquitin-like domain ; CUTL = cut-Like domain; CUT1 = cut domain 1; CUT2 = cut domain 2; HOX = homeobox; INM = inner nuclear membrane; MAR = matrix associated region; NLS = nuclear localization signal. **B**, Relative *SATB2* and *circ3915* expression, as measured by RT-qPCR. *SATB2* expression increased 18.7-fold and *circ3915* expression increased by 15.6-fold. N = 3 replicates; **** *p* <0.0001. **C**, Representative Western blot showing SATB2 up-regulation in iAsT BEAS-2B cells. **D**, *SATB2* (purple) and *circ3915* (orange) expression in BEAS-2B cells after acute, 48 hr exposure to iAs. Fold-change was measured by RT-qPCR, and calculated relative to non-treated BEAS-2B cells. **E**, *SATB2* and *circ3915* expression in BEAS- 2B cells at different time-points in the two-hit iAs transformation model, as measured by Nanostring. N= 3 replicates; * *p* <0.05. **F**, *SATB2* and *circ3915* expression in normal (N) and adjacent tumor (T) samples, as measured by RT-qPCR. All expression data were normalized to the expression of the geometric mean of three common husekeeping genes (*GAPDH, TBP, RPII*).

We next investigated if *SATB2* and *circ3915* expression were detectable in human samples. We obtained paired primary tumors and normal adjacent tissues from lung adenocarcinoma (LUAD) and lung squamous cell carcinoma (LUSC) patients. Among the 40 paired LUAD and 20 paired LUSC samples, 24 and 8 showed upregulation of *SATB2* and *circ3915*, respectively (**Fig 3F**). Conversely, when *SATB2* expression decreased, so too did *circ3915* expression. Collectively, these data indicate that iAs exposure triggers *SATB2* over-expression in multiple cell lines and exposure models, resulting in a novel *circ3915* RNA. Both RNA species persist through the transformation process and are found in human normal and lung tumor specimens. These findings suggest a general carcinogenic mechanism that may be related to chromatin remodeling.

#### *circ3915* is translated and contributes to SATB2-chromatin organization in the nucleus

*circ3915* has an ATG start codon, putative nuclear localization signal, oligomerization domain, and inner nuclear membrane domain but no DNA binding domain (**Fig 3A**). We therefore asked if *circ3915* is translated into a peptide (circ3915p), and if that peptide might have a biological function. Since there was no specific antibody available for the putative circ3915 product, we introduced a HiBiT tag into the *circ3915* expression construct and monitored circ3915p translation by bioluminescence. When we transduced BEAS-2B cells with HiBiT-tagged *circ3915* expression constructs, we saw enhanced *in vitro* bioluminescence (**Fig 4A**) and HiBiT protein in Western blots (**Fig 4B**), indicative of circ3915-HiBiT translation. Approximately 60% of all *circ3915* transcripts were associated with actively translated polysomes, with puromycin treatment shifting *circ3915* transcripts towards light and free polysome fractions (**Fig 4C**, **Figs S4A**); similar results were obtained in iAs-transformed BEAS- 2B cells (**Fig 4C**). In contrast, an untranslated SATB2 circRNA hsa_circ_0008928^71^ did not associate with polysomes or show a shift in polysome association (**Fig S4B**).

**Fig 4.**
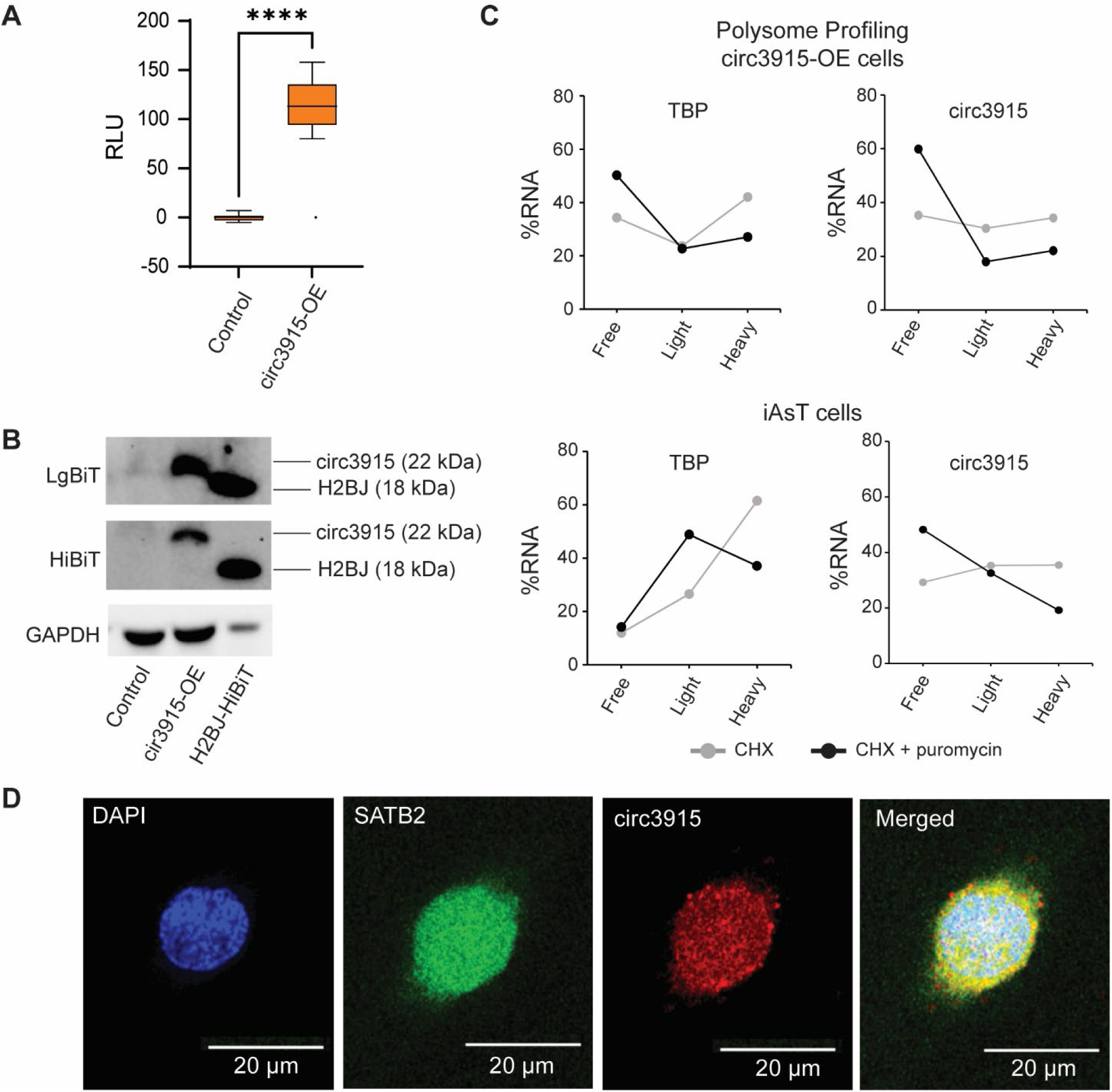
***circ3915* is translated into a peptide, and co-localizes with SATB2 in the nucleus**. **A,** Bioluminescence in BEAS-2B cells containing an empty vector (control), or cells that are over-expressing a HiBiT-tagged *circ3915* construct (circ3915-OE), as measured with a split luciferase translation assay. RLU = relative light units; N= 3; **** *p* <0.0001. **B**, Western blots for 100 µg of total protein or 5 µg of nuclear extract from BEAS-2B cells transfected with an empty vector (control), HiBiT-tagged *circ3915* construct (circ3915-OE), or a HiBiT-tagged histone 2BJ (H2BJ) positive control. HiBiT was detected with either LgBiT (top panel) or an anti-HiBiT antibody (middle panel). **C**, *circ3915* is associated with actively translated polysomes. BEAS-2B cells were either transfected with a circ3915 HiBiT overexpression plasmid (top), or transformed with iAs in the rapid (12 week) transformation model (bottom). Line graphs show the distribution of cytoplasmic RNA across sucrose gradient density fractions. Grey lines show the distribution of RNA under normal culture conditions (CHX = cycloheximide), and black lines show the distribution of RNA after adding puromycin (puromycin disrupts translation elongation). An increase in %RNA within the light fractions after treating cells with puromycin indicates that the RNA was being translated under normal conditions. TBP = TATA-box binding protein mRNA. **D**, Fluorescent microscope images of BEAS2B cells that were transduced with SATB2-GFP and circ3915-HiBiT expression vectors, showing that circ3915-peptide localizes in the nucleus. DAPI-stained DNA (blue) denotes the nucleus.

Initial immunocytochemistry experiments showed that circ3915p co-localizes with SATB2 in the nucleus of stably-transduced BEAS-2B cells (**Fig. 4D**), and alpha-fold predictions suggested that circ3915p might oligomerize with SATB2 (**Fig S4C**). To validate that circ3915p was associated with SATB2, we over-expressed FLAG-tagged SATB2 or circ3915 in non-transformed BEAS-2B cells, immunoprecipitated FLAG-tagged protein complexes and then analyzed the immunoprecipitated proteins by mass spectrometry. After filtering out keratins and immunoprecipitated proteins detected in control (empty vector) cells, we identified 24 immunoprecipitated proteins that were unique to SATB2, 23 that were unique to circ3915p, and 28 that were common to both (**Table S4**). Importantly, SATB2 peptides (69% coverage) were identified in the SATB2 and circ3915 over-expressing cell lines but not in cells transfected with an empty vector. Thus, circ3915p associates with full-length SATB2.

Interestingly, SATB2 and circ3915p also immunoprecipitated with chromatin and chromatin remodelers (H1, H2A.Z, macroH2A, and HDAC2), splicing factors (SNRPB, SNRPD2, SNRPD3, SRSF3, SRSF6, SRSF9, TRA2B), and the nuclear envelop protein BANF1. The chromatin proteins associated with both SATB2 and circ3915p are usually associated with transcriptional repression, while chromatin proteins unique to SATB2 (H2A.X, PARP1, and SSRP1) are often associated with active chromatin. These results suggest SATB2 regulates gene expression by directing changes to chromatin structure. HDAC2 association with circ3915p further suggests that circ3915p (which lacks SATB2 DNA binding domains; **Fig 3A**) might sequester repressive complexes away from cis-regulatory elements. SATB2 and circ3915p both interact with CEBPB, which is also consistent with the enriched transcription factor motifs in differentially accessible genomic regions (below). Taken together, these data indicate that *circ3915* is translated and the resulting peptide (circ3915p) co-locates with SATB2 in the nucleus, where it interacts with SATB2 and/or chromatin to modify chromatin architecture.

### *circ3915* “protects” the linear *SATB2* transcript

Some circRNAs can protect their cognate mRNA from post-transcriptional regulatory pathways^72,73^, and the co-expression of *SATB2* and *circ3915* RNA suggested a similar co-regulatory mechanism. We, therefore, forced *SATB2* or *circ3915* over-expression in non-transformed BEAS- 2B cells (where *SATB2* mRNA is normally absent) and quantified the corresponding RNAs by qRT-PCR. Forced/exogenous *SATB2* expression increased *SATB2* mRNA and protein levels (as expected; **Fig 5A,B**), and also increased *circ3915* expression (**Fig 5B**). In contrast, forced/exogenous *circ3915* expression had no effect on endogenous *SATB2* mRNA expression (**Fig 5C**). We then knocked down *SATB2* or *circ3915* RNAs in iAs- transformed cells. In this case, knocking down linear *SATB2* decreased SATB2 mRNA levels (as expected), but had no effect on *circ3915* RNA levels (**Fig 5D**). However, knocking down *circ3915* depleted both *circ3915* and *SATB2* RNA levels in iAs-transformed cells (**Fig 5E**), with a corresponding decrease in SATB2 protein (**Fig 5F**). Thus, whenever there is *SATB2* over-expression, *circ3915* is generated. These data imply that *SATB2* regulates its own expression and that *circ3915* may “protect” the linear transcript from degradation. The data also suggest that *SATB2* and *circ3915* are required for maintaining an oncogenic or cancerous cell state.

**Fig 5.**
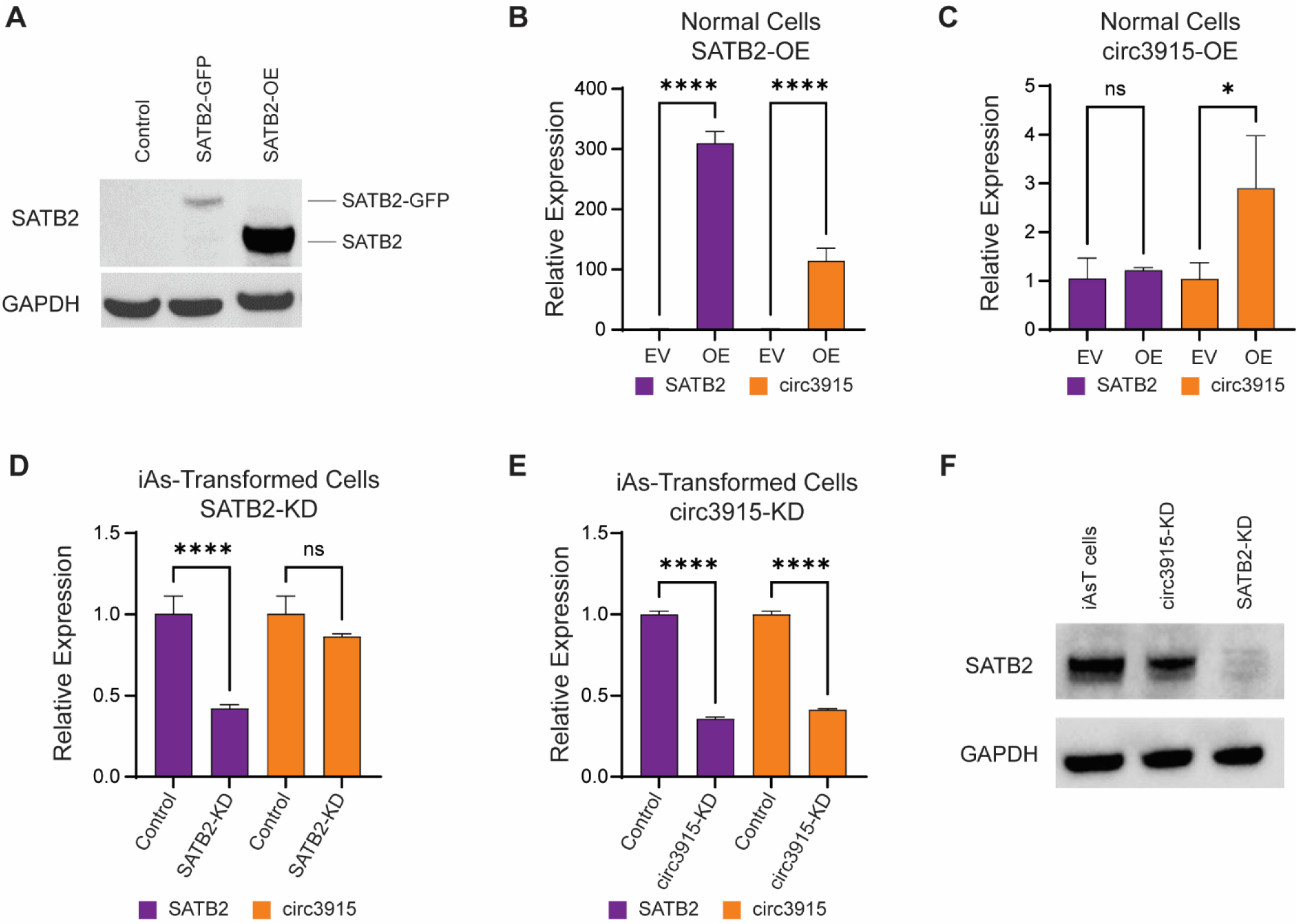
*circ3915* protects the linear SATB2 transcript. **A**, Western blot showing forced SATB2 and SATB2- GFP expression in non-transformed BEAS-2B cells. **B**, Relative *SATB2* and *circ3915* expression in BEAS-2B cells containing an empty vector (EV) or SATB2 over-expression vector (OE), as measured by RT-qPCR. N = 3 replicates; **** *p* <0.0001. **C**, Same as panel **B**, except with forced over-expression of *circ3915*. N = 3 replicates ns = not significantly different; * *p* <0.05. **D,** BEAS-2B cells were transformed with iAs (rapid transformation model), and then treated with siRNAs to knock down *SATB2* (SATB2-KD). *SATB2* and *circ3915* RNAs were then measured by RT-qPCR. N = 3 replicates. ns = not significantly different; **** *p* <0.0001. **E**, same as panel **D**, except cells were treated with siRNAs to knock down *circ3915*. N = 3 replicates; **** *p* <0.0001. **F**, Western blot of SATB2 protein levels in iAs-transformed BEAS-2B cells after knocking down *circ3915* or *SATB2* with target-specific siRNAs.

### SATB2 and circ3915 epigenetically regulate oncogenic gene expression programs

To determine if SATB2 and circ3915 are required for oncogenic transformation, we used lentivirus vectors to consistently over-express *SATB2* mRNA and/or *circ3915* in non-transformed BEAS-2B cells. In all over-expression conditions, we observed enhanced cell proliferation and migratory potential relative to BEAS-2B cells containing an empty lentivirus vector, with the most pronounced effects observed in those cells with forced over-expression of both circular and linear transcripts (**Fig 6A-C**). *SATB2* over-expression dysregulated many genes in non-transformed cells (**Fig 6D**; **Fig S5A**,**B; Table S5**, resulting in a gene expression pattern that was more like iAs-transformed cells than non-treated cells (**Fig 6E**, left panel). In fact, over-expressing *SATB2* in BEAS-2B cells recapitulated ∼35% of the gene expression patterns found in iAs-transformed cells (**Fig S5C**). As in iAs-transformed cells, *SATB2* over-expression caused an increase *HMOX1*, *GOS2*, *CCND1*, and *MMP1* expression, and a decrease in *FAT3*, *KRT17*, *DIO2*, *PBX1*, and *HLA-B* expression (**Fig 6E**, right panel). *SATB2* over-expression also dysregulated key oncogenic signatures, including a striking enrichment in oncogenic KRAS-like signatures (**Fig 6F,G**). Thus, *SATB2* (hence *circ3915*) over-expression epigenetically regulates oncogenic gene expression programs and cell transformation, even without iAs exposure or driver DNA mutations.

**Fig 6.**
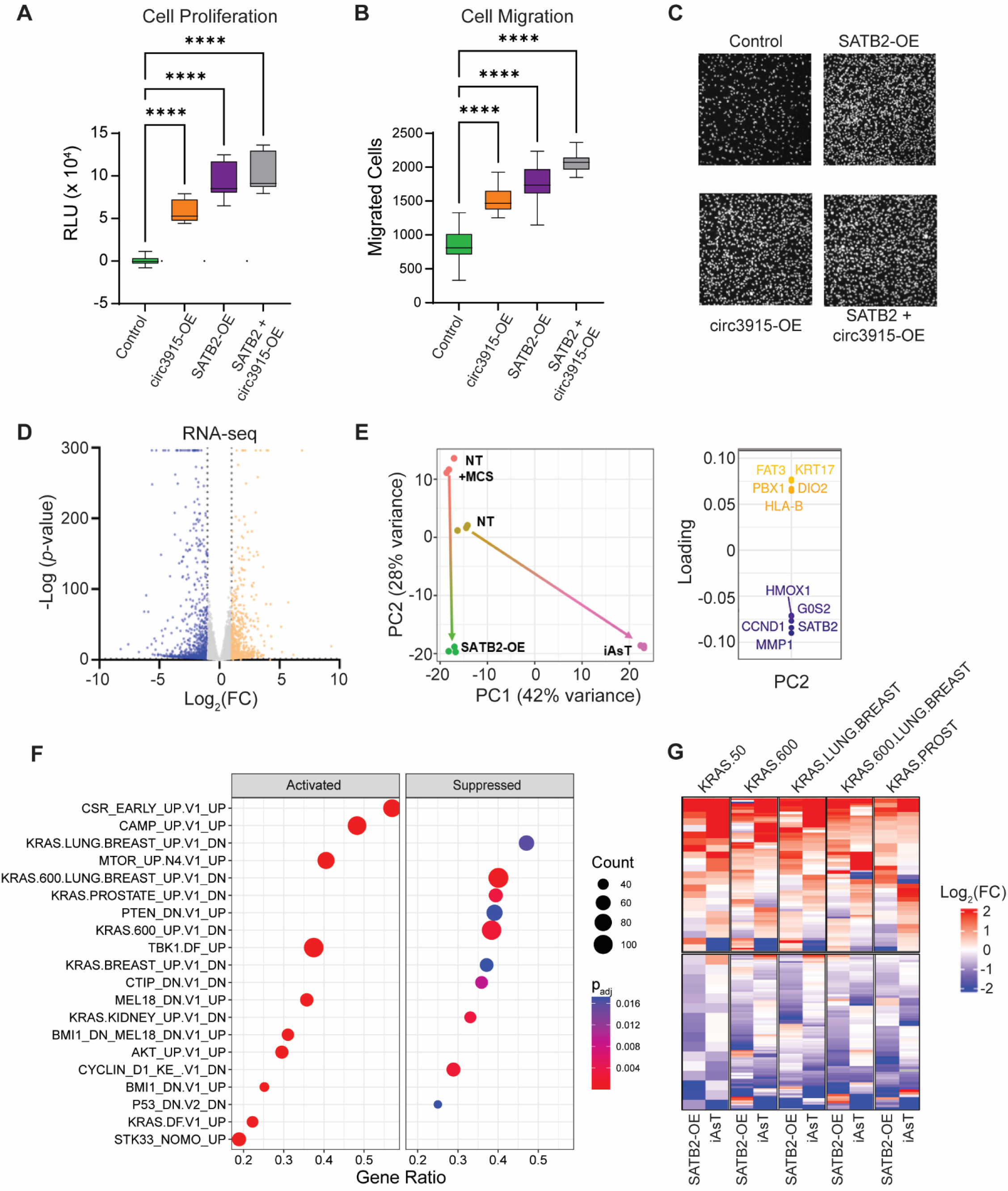
*SATB2* over-expression promotes oncogenic transformation. **A**, BEAS-2B cell proliferation after transfection with lentivirus over-expression vectors, as measured with a CellTiter-Glo ATP assay 72 hours after seeding. Control = non-transformed cells containing an empty lentivirus vector. SATB2 + circ3915-OE = cells that contained SATB2 and circ3915 overexpression vectors. RLU = relative light units. Luminescence was normalized to background, and values calculated relative to the control cells. N = 3 independent experiments with 6 technical replicates each; **** *p* <0.0001. **B**, BEAS-2B cell migration after transfection with lentivirus over- expression vectors, as measured by a transwell migration assay 24 hr after seeding. Cells were fixed and stained with DAPI, and cell counts obtained with an Olympus cellSens counting module. Counts were normalized to the control cells. N = 3 independent experiments with 3 technical replicates each **** *p* <0.0001. **C**, Representative images from the transwell migration assay. **D**, Differentially expressed genes (DEGs) in BEAS-2B cells with forced SATB2 over-expression. Blue dots represent down-regulated genes with a *p_adj_* <0.05, FC <-1.5; orange dots represent up-regulated genes with a *p_adj_* <0.05, FC >1.5. Grey dots represent genes with FC<1.5,>-1.5. **E**, Principal component analysis (left) of RNA-seq expression profiles from non-transformed BEAS-2B cells (NT, brown); non-transformed cells that were transfected with a SATB2 overexpression vector (NT + MCS, red); non- transformed cells containing a SATB2 overexpression vector (SATB2-OE, green); and iAs-transformed BEAS- 2B cells from the rapid (12 week) transformation model (iAsT, pink). The top genes contributing to the variance between PC1 and PC2 are shown in the panel to the right. **F**, Top enriched MSigDB oncogenic signatures reflected in the DEGs from non-transformed BEAS-2B cells with forced SATB2 over-expression. **G**, Relative RNA expression of MSigDB oncogenic KRAS signature genes in non-transformed BEAS-2B cells that over-express SATB2 (SATB2-OE) or in iAs-transformed BEAS-2B cells (iAsT, rapid transformation model).

### SATB2-dependent chromatin modifications are associated with KRAS-like gene expression patterns

SATB2 organizes chromatin by binding to AT-rich cis-regulatory elements and then associating with other SATB2 molecules and the inner nuclear matrix, thus facilitating long-range DNA interactions^36–38,46^. To test for SATB2-dependent chromatin changes, we performed ATAC-seq with the same lentiviral constructs and transformed BEAS-2B cells from above. We found that *SATB2* over-expression opened 19383 genomic regions and closed 14014 others (**Fig 7A, Table S6**), with clear distinctions within the top 50 differentially accessible peaks (**Fig 7B**). Principal component analysis of DARs in non-treated, iAs-transformed, lentivirus-transformed, and SATB2-overexpressing BEAS-2B cells (**Fig 7C**) suggest that a significant fraction of the DARs in iAs- transformed cells could be attributed to SATB2. Like the differentially accessible regions in iAs-transformed cells (**Fig 2** and **Fig S2**), most of the SATB2*-*dependent DARs were found in distal enhancer-like regions, but there were no obvious differences in the genomic distribution of opened/closed DARs or ENCODE cCRE signatures (**Fig S6A**). Motif analysis of the SATB2-dependent DARs identified 221 enriched transcription factor binding sequences (Benjamini q-value <0.01), with 116 enriched in open chromatin and 105 in closed chromatin regions. There was a surprising number of transcription factor binding motifs that were shared between SATB2 over- expressing and iAs-transformed cells, including the NFE2L2 binding site (**Fig 7D**). *De novo* motif discovery identified 12 binding sequences significantly enriched in open chromatin regions (*p* <10^-11^), including binding sites for BATF, TEAD3, ERG, c-JUN, and AP2 (**Fig 7E**), all transcription factors with established roles in oncogenesis^74–78^. Conversely, regions with reduced accessibility were significantly enriched for 13 *de novo* binding sequences, including the NFY/CCAAT promoter, SP2, AP-1 ELF1, and YY2 (**Fig 7E**; **Table S7**), a pattern also observed in closed chromatin from iAs-transformed cells (**Fig 2E**). Thus, SATB2 may regulate the CCAAT/enhancer binding protein (C/EBP) family of bZIP transcription factors that govern cell proliferation, apoptosis, and stress response^79–81^.

**Fig 7.**
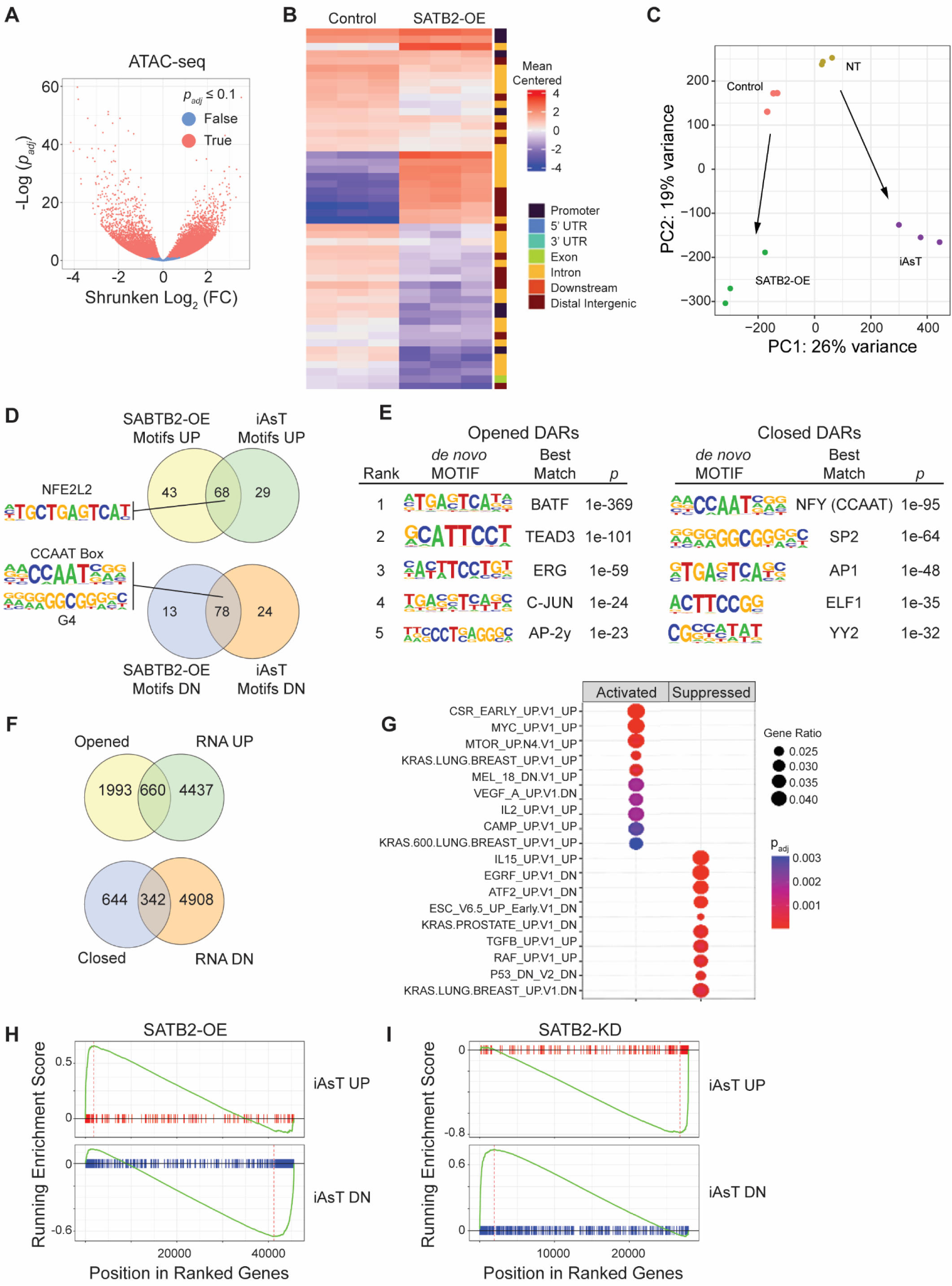
*SATB2* over-expression alters chromatin accessibility at cis-regulatory elements. **A**, Differentially accessible regions (DARs) between non-treated BEAS-2B cells containing an empty expression vector, and BEAS-2B cells containing a SATB2 over-expression construct. DARs were defined by an FDR (*p_adj_*) ≤0.1. **B**, Heatmap of the top 50 DARs between non-transformed BEAS-2B cells containing an empty expression vector (control) or over-expressing *SATB2* (SATB2-OE). Columns are the mean centered peak values for each gene (row) in each sample. The genomic location of each peak is shown to the right of the heatmap, with most peaks occurring in promoters, introns, and distal intergenic regions. **C**, Principal component analysis of DARs from non-transformed BEAS-2B cells (NT, green); non-transformed cells that were transfected with a SATB2 overexpression vector (NT + MCS, red); non-transformed cells over-expressing SATB2 (SATB2-OE, blue); and iAs-transformed BEAS-2B cells (rapid transformation model; iAsT, purple). **D**, Overlapping transcription factor motifs between BEAS-2B cells over-expressing *SATB2* (SATB2-OE) and iAs-transformed BEAS-2B cells (rapid transformation model). **E**, Top enriched *de novo* transcription factor motifs and best matching transcription factors in opened and closed DARs from BEAS-2B cells over-expressing *SATB2*. **F**, Congruence between SATB2-OE DARs and proximal gene expression. RNA UP = up-regulated transcripts; RNA DN = down-regulated transcripts. Counts are the number of individual genes above the minimum fold-change threshold for mRNA expression (>1 Log_2_(FC)) or open chromatin (>1 ShrunkenLog_2_(FC)). **G**, Top enriched C6 oncogenic signatures in non-transformed BEAS-2B cells that over-express *SATB2*. **H,** GSEA plot shows that genes upregulated in the KRAS pathway in iAsT cells were also enriched in SATB2OE cells, while downregulated genes were similarly downregulated. **I**, In SATB2-KD cells, this enrichment is reversed: previously upregulated genes in the KRAS pathway are downregulated, and previously downregulated genes are upregulated.

We next asked how SATB2-dependent changes in chromatin accessibility related to changes in gene expression. We again annotated DARs to their nearest genes, and compared proximal gene expression to (open vs. closed) chromatin state. Of the 2643 opened DARs, 660 proximal genes (25%) were up-regulated; of the 986 closed DARs, 342 proximal genes (35%) were down-regulated (**Fig 7F**). The seemingly low correlation between DAR proximity and gene expression may be due to gene regulation by distal enhancer elements that are spatiotemporally arranged across chromosomes^82^. This hypothesis is supported by altered accessibility at ENCODE enhancer signatures and CCAAT enhancer binding sites in SATB2-overexpressing cells and iAs- transformed cells (e.g., **Fig 2E** and **7E**).

Next, we performed GSEA on the SATB2-dependent DARs and DEGs. Among 79 enriched oncogenic signatures in DEGs (**Table S8**), eight up-regulated signatures occurred within newly opened chromatin, including KRAS signatures (**Fig 7G; Fig. S6B,C**) originally identified in iAs-transformed cells (**Fig 1D**). Gene set enrichment analysis (GSEA) of the *SATB2* over-expressing cells also showed enrichment for the NFE2L2-like oncogenic signature (**Fig S6D**), and other pathways (**Fig. S6E**) that were first observed in iAs-transformed cells. When we performed GSEA on the SATB2 overexpression ranked gene set vs. the iAs-transformed ranked gene set, *SATB2* over-expression (in non-treated) cells showed positive enrichment for upregulated and downregulated genes in iAsT cells (**Fig 7H**). Further, knocking down *SATB2* resulted in negative enrichment of those genes that were upregulated (**Fig 7I**, top panel) and downregulated (**Fig 7I**, bottom panel) gene sets in the iAs-transformed cells. These data show that SATB2 regulates these pathways in iAsT cells and that knocking down SATB2 reverses the gene expression profiles. Thus, SATB2-dependent changes in chromatin accessibility are responsible for a significant portion of the oncogenic KRAS signatures found in iAs-transformed cells.

### *SATB2* and *circ3915* knockdown in transformed cells reverse oncogenic gene expression programs

Having demonstrated that *SATB2* (and *circ3915*) over-expression is sufficient to activate oncogenic KRAS gene expression patterns in non-transformed BEAS-2B cells, we asked if knocking down *SATB2* and/or *circ3915* in iAs-transformed cells could reverse these oncogenic signatures and gene expression patterns. We transfected iAs- transformed cells with lentiviral vectors containing *SATB2*- and *circ3915*-targeted shRNAs and then performed RNA-seq. Successful knockdown was confirmed by Western blots and qRT-PCR (**Fig 8A-C**). Knocking down *SATB2* mRNA affected 4421 genes, and knocking down *circ3915* affected 4085 genes (*p_adj_*. ≤ 0.05; **Fig S7A**). Principal component analysis (**Fig 8D**) showed that SATB2 and circ3915 knockdown cells clustered together, indicating that the two transcripts control similar genes (as expected). Knockdown also shifted gene expression patterns away from *SATB2* over-expressing cells (PC1) and iAs-transformed cells (PC2), suggesting a subset of gene expression changes are dependent on, but not directly driven by, *SATB2* over-expression. GSEA confirmed that *SATB2* and *circ3915* knockdown silenced the oncogenic signatures and pathways identified in iAs- transformed and *SATB2* over-expressing cells, including the oncogenic KRAS and NFE2L2 signatures (**Fig 8E, Fig S8B-D**). Knocking down either transcript also decreased the proliferative and migratory capacity of iAs- transformed cells (**Fig 8F-H**).

**Fig 8.**
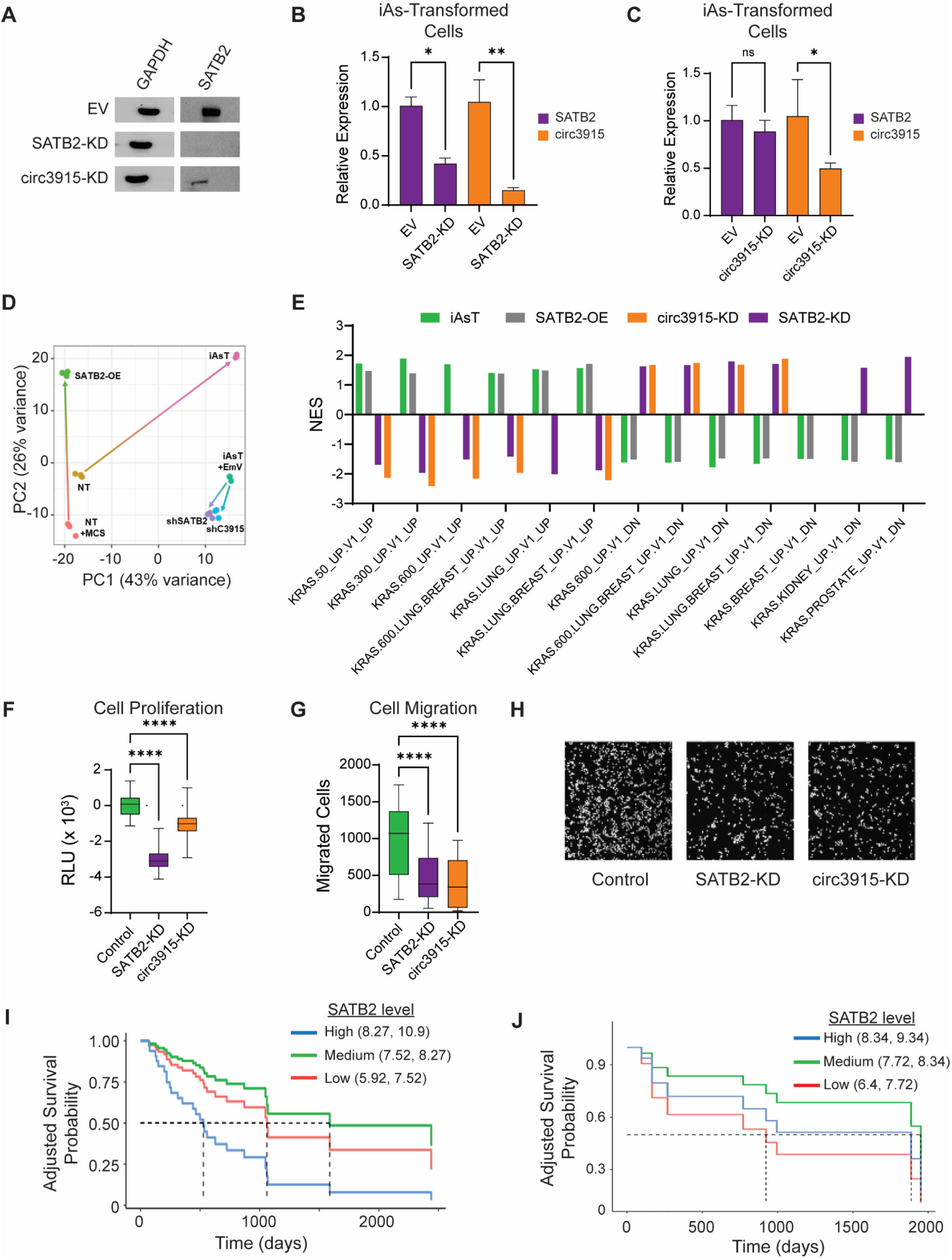
Knocking down *circ3915* or *SATB2* in iAs-transformed BEAS-2B cells reverses oncogenic KRAS- like gene expression. **A**, Western blot showing lentiviral SATB2 knockdown in iAs-transformed BEAS-2B cells. EV = empty vector control. **B**, Relative *SATB2* and *circ3915* expression in iAs-transformed BEAS-2B cells after *SATB2* knockdown with shSATB2, as measured by RT-qPCR. EV = empty vector control. N = 3 replicates; * *p* <0.05. **C**, Same as panel **B**, except after *circ3915* knockdown with shcirc3915. N = 3 replicates; * *p* <0.05, ns = not statistically significant. **D**, Principal component analysis (left) of RNA-seq expression profiles from non- transformed BEAS-2B cells (NT; brown); non-transformed cells containing an empty lentiviral expression vector (NT + MCS; red); non-transformed cells over-expressing SATB2 (SATB2-OE, green); iAs-transformed cells (iAsT; pink); iAs-transformed cells containing an empty lentiviral vector (iAsT + EmV; turqoise); iAsT cells with stable SATB2 knockdown (shSATB2; purple); and iAsT cells with stable circ3915 knockdown (shC3915; light blue). iAs-transformed cells were generated with the rapid (12 week) transformation model. Arrows indicate how RNA expression profiles shifted from control samples to the matching experimental group. **E**, Bar plot of normalized enrichment scores for significantly enriched (*p_adj_* <0.05) MSigDB oncogenic signatures in iAs- transformed BEAS-2B cells (iAsT, rapid transformation model), non-transformed cells overexpressing SATB2 (SATB2-OE), BEAS-2B cells with stable *SATB2* knockdown (SATB2-KD), and BEAS-2B cells with stable *circ3915* knockdown (circ3915-KD). **F**, iAs-transformed BEAS-2B cell proliferation after shRNA knockdown, as measured with a CellTiter-Glo ATP assay 72 hours after seeding. Control = iAs-transformed cells containing an empty lentivirus vector. RLU = relative light units. Luminescence was normalized to background, and values calculated relative to the control cells. N = 3 independent experiments with 6 technical replicates each; **** *p* <0.0001. **G**, iAs-transformed BEAS-2B cell migration after after shRNA knockdown, as measured by a transwell migration assay 24 hr after seeding. Cells were fixed and stained with DAPI, and cell counts obtained with an Olympus cellSens counting module. Cell counts were normalized to the control cells. N = 3 independent experiments with 3 technical replicates each **** *p* <0.0001. **H**, Representative images from the transwell migration assay of panel **I, J**. Kaplan-Meier plots of overall survival relative to SATB2 expression in patients that either did not have a KRAS mutation (**I**, n=342) or harbored a KRAS mutation (**J**, n=150). Patient data were controlled for age at diagnosis, sex, and pack years.

Because *SATB2* up-regulation is associated with lung cancers in the TCGA database^83^ (**Fig 3F**), we next asked if *SATB2* expression is associated with survival. Reasoning that *SATB2* expression might be driving a *KRAS*-like phenotype, we separated tumor samples (from the TCGA database^83^) into those with (n = 159) or without (n = 342) a *KRAS* mutation. We then used differential expression analysis (|Log_2_(FC)| > 1, p < 0.01) and Cox Proportional Hazard methods to control for SATB2 expression, age at diagnosis, gender, and race (filtered for White and Black/African American individuals due to limited data availability for other groups), and smoking status (calculated in pack years). We then censored for long-term survival by filtering individuals who survived at least 2 years and tracking them until the last follow-up date. When controlled for these factors, *SATB2* expression was associated with poor long-term survival in patients who did not harbor a *KRAS* mutation (HR = 1.85, p < 0.005; **Fig 8I**), but not for patients who did have a *KRAS* mutation (HR = 0.979, p = 0.968; **Fig 8J)**. These data support the hypothesis that *SATB2* over-expression mimics a *KRAS*-like signature and that *SATB2* expression is not necessary if a *KRAS* mutation is present. Thus, SATB2 epigenetically regulates *KRAS* oncogenic gene expression patterns.

## DISCUSSION

### SATB2 regulates its own biogenesis and produces a novel circRNA

In this study, we show that SATB2 regulates its own biogenesis and identify a novel *SATB2* circular RNA (*circ3915*) originating from the *SATB2* gene that helps maintain the linear *SATB2* transcript. These data are consistent with other studies showing that circRNAs can protect their cognate mRNA from post-transcriptional regulatory pathways^67,72^and suggest that SATB2 directly or indirectly regulates its own splicing. The novel *circ3915* RNA is co-generated upon *SATB2* mRNA expression in two lung cell lines and is detected in human lung tumors. Interestingly, *circ3915* is also translated in iAs-transformed BEAS-2B cells, a feature not yet seen with *circSATB2* transcripts. The resulting peptide (circ3915p) is translocated to the nucleus, where it co-locates with full-length SATB2 and associates with a range of nuclear factors, including chromatin and chromatin remodelers, splicing factors, and transcription-related proteins. Thus, circ3915 promotes oncogenic transformation and cancer cell proliferation by “protecting” *SATB2* mRNA and producing a circ3915 peptide that may interact with chromatin remodeling machinery.

### SATB2 alters chromatin adjacent to known oncogenes and *KRAS*-related transcription factors

iAs exposure caused significant changes in chromatin accessibility around promoters, promoter-like structures, and transcription factor binding sites. Genes within 1 Kb of these differentially accessible regions (DARs) included those associated with the epithelial-to-mesenchymal transition, C6 oncogenic signatures (i.e., *KRAS* and *EGFR*), and oxidative stress response (*NFE2L2*). Motif analysis of “opened” DARs likewise pointed to transcription factors related to *KRAS* activation, while “closed” motifs related to the CCAAT/enhancer binding protein (C/EBP) family of basic region-leucine zipper (bZIP) transcription factors that regulate cell proliferation, apoptosis, and stress response^79–81^. We over-expressed *SATB2* mRNA in non-treated BEAS-2B cells to identify chromatin regions and genes that were specifically regulated by SATB2 and circ3915 rather than iAs and the oncogenic transformation process and found that SATB2/circ3915-dependent DARs were enriched in distal intergenic regions and intergenic sites. It is difficult to correlate distal enhancer accessibility (ATAC-seq) with non-proximal gene expression (RNA-seq), but future Hi-C interaction assays^84^ might help us identify which distal enhancer regions target promoters and up-regulated genes identified here. Nevertheless, open chromatin was again associated with the KRAS oncogenic signature and NFE2L2 transcription binding sites, and closed chromatin pointed to reduced accessibility of CCAAT enhancer binding sites. In total, ∼20% of iAs-induced changes in chromatin accessibility could be attributed to SATB2 up-regulation. Thus, SATB2 over-expression causes chromatin changes that are associated with known oncogenes and gene expression profiles.

### SATB2/circ3915 regulate two oncogenic gene expression programs

iAs-induced and SATB2-dependent changes to chromatin accessibility were likewise reflected in the transcriptome. Of the ∼8400 genes that were differentially expressed during iAs-induced transformation, ∼28% of them could be attributed to SATB2 over- expression. We were surprised to find that SATB2-regulated genes were enriched for oncogenic signatures, including a striking enrichment in KRAS- and NFE2L2-associated signatures. *KRAS* mutations are found in ∼30% of all lung cancer cases^85,86^, but our data show that SATB2 overexpression can stimulate KRAS-related oncogenic pathways independent of *KRAS* mutations. NFE2L2 is also interesting because it is a master regulator of the antioxidant response^87^, and sustained upregulation of antioxidative response genes supports cell proliferation, resistance to apoptosis, and cancer cell survival after chemotherapy^88^. In renal cancer, SATB2 coordinates enhancer-promoter looping to boost NFE2L2-mediated gene expression^40^, and we suspect a similar mechanism is operating here. Importantly, knocking down *SATB2* and/or *circ3915* expression in transformed BEAS-2B cells reversed the KRAS and NFE2L2 gene expression signatures. Thus, *SATB2* and *circ3915* overexpression epigenetically transforms lung epithelial cells by reprogramming gene networks commonly activated in KRAS- and NFE2L2-driven lung cancers.

### SATB2 and circ3915 are potential therapeutic targets

Chronic iAs exposure causes cancer^23,89–94^, as does *SATB2* over-expression^48,51–54,95^ and the production of SATB2 circular RNAs^51,96,97^. The work presented here is, therefore, consistent with a model where low-dose iAs exposure induces a mild state of oxidative stress that elicits *SATB2* and *circ3915* expression. *SATB2* and *circ3915* are translated and translocated to the nucleus, where they engage with chromatin to sustain pro-cancerous gene expression patterns and promote oncogenic transformation. While it is not yet clear if SATB2 is a generalized therapeutic target for different cancers, our knockdown studies suggest that *circ3915* might be a viable therapeutic target for iAs-induced cancers or (by extension) smoking-related lung cancers that are not driven by a *KRAS* mutation. Because *SATB2* and *circ3915* dysregulation generate *KRAS*-like gene expression programs, we might also speculate that therapies targeting downstream, *KRAS*- dependent pathways might benefit patients with epigenetically driven, *SATB2*/*circ3915*-driven cancers.

While this study used iAs to transform bronchial epithelial cells, our system is just as much an epigenetic model for how 3D chromatin structures regulate (oncogenic) gene expression programs^10–13^ as it is a model of iAs-induced carcinogenesis^29,57^. Future studies might, therefore, focus on *SATB2* and *circ3915* expression levels as potential diagnostic biomarkers for those (lung) cancers where there are no detectable DNA mutations and possibly other environmentally induced cancers with an epithelial cell-of-origin. Since *SATB2* over-expression triggered an NFE2L2 antioxidant response, it is tempting to speculate that chemo-resistant lung cancers might also be characterized by *SATB2* or *circ3915* over-expression. Understanding how SATB2 and its multiple circRNAs drive various cancers (or drug resistance) will require knowledge about alternative *SATB2* splicing mechanisms in different cell types and cancers, the specific genomic loci bound by SATB2, and how SATB2 might alter nucleosome positioning^98,99^ to modify transcription. It will also be important to understand how SATB2 and circ3915p interact with chromatin remodelers to bring distal enhancers into close proximity to transcription factors and gene promoters, thus regulating oncogenic gene expression programs.

In summary, this study details how an environmental signal can trigger *SATB2* and its *circ3915* to transform lung epithelial cells without any DNA mutations. In dissecting the role of *circ3915*, we reveal that it protects *SATB2* mRNA from post-transcriptional degradation, is translated into a peptide, and supports SATB2 function in the nucleus. *Circ3915* is a new biomarker for iAs-induced lung epithelial transformation that might also be a biomarker for other environmental insults in different cells of origin. In the same way that SATB2 reprograms cells during development, our integrated analysis demonstrates how SATB2 transcriptionally reprograms cells towards an oncogenic state, and shows that epigenetic mechanisms alone can trigger malignant transformation in mammalian systems. Taken together, these findings should generate new avenues for developing lung cancer diagnostics and interventions for those patients who do not harbor cancer-driving mutations.

## ACKNOWLEDGEMENTS

We thank the Fondufe-Mittendorf lab for scientific discussions contributing to this manuscript and members of the VAI Core Facilities (Genomics, Bioinformatics, and Biostatistics) for technical assistance. We also thank the VAI Pathology and Biorepository Core (RRID:SCR 022912) for collecting the de-identified human lung tumor samples. Y.F.M. is supported by the National Science Foundation grant NSF/MCB 016515, National Institutes of Health grants R01ES024478, R01ES034253, and 1R01ES036051-01, and the Van Andel Institute (VAI). R.E. is supported by an NSF Graduate Fellowship (GRF-1839289).

## AUTHOR CONTRIBUTIONS

Conceptualization: RE. and YFM; investigation: RE, SG, LS, WS, KL, and YFM; data analysis: RE, SG, LS, WS, KL, and YFM.; data curation: RE, SG, LS, WS, KL, and YFM; writing—original draft preparation: RE, SG, DPC, and YFM; writing-editing: RE, SG, DPC, and YFM; visualization: RE, SG, LS, KL, DPC, and YFM; resources, YFM; supervision: YFM; funding acquisition: YFM. All authors have read and agreed to the published version of the manuscript.

## Data Availability Statement

Data analyzed have been deposited in GEO with accession numbers GSEXXXXX.

## SUPPLEMENTAL INFORMATION

- Document S1. Supplementary Figures S1-S8.
- Table S1. Excel file of differentially expressed genes in iAs-transformed BEAS-2B cells.
- Table S2. Excel file of differentially accessible regions in iAs-transformed BEAS-2B cells.
- Table S3. Excel file(s) of transcription factor binding sites/motifs in iAs-transformed BEAS-2B cells.
- Table S5. Excel file of dysregulated genes in non-transformed BEAS-2B cells that over-express *SATB2*.
- Table S6. Excel file of differentially accessible regions in non-transformed BEAS-2B cells that over- express *SATB2*.
- Table S7. Excel file(s) of transcription factor binding sites/motifs in non-transformed BEAS-2B cells that over-express *SATB2*.
- Document S2. Supplementary Tables S4, S8-S12.

## STAR METHODS RESOURCE AVAILABILITY

### Lead contact

Further information and requests for resources and reagents should be directed to and will be fulfilled by the lead contact, Yvonne L. Fondufe-Mittendorf).

### Materials availability

Plasmids generated for this study are available upon request from the lead contact, without restriction.

### Data and code availability

This paper does not report the original code. The lead contact can provide any additional information required to reanalyze the data upon request.

## METHOD DETAILS

### Patient samples

All de-identified patient samples used in this study were obtained from the Van Andel Institute Pathology and Biorepository Core (RRID:SCR_022912) and histologically confirmed as either adenocarcinoma or sarcoma. Lung tissues were snap-frozen in liquid nitrogen, pulverized, and RNA purified using a Qiagen RNeasy Plus Universal Mini Kit.

### Cell growth conditions and iAs-induced transformation

BEAS-2B cells were cultured in Dulbecco’s Modified Eagle’s Medium (DMEM) supplemented with 10% fetal bovine serum (v/v) and 100 U/mL penicillin- streptomycin. Cells were grown to ∼80% confluency in a humidified chamber at 37°C, 5% CO_2_. The Human Bronchial Epithelial cell line (16HBE) was a gift from Dr. Shaun McCullough^100^ and grown on tissue culture flasks coated with type I collagen. Cells were cultured in Modified Eagle’s Medium (Advanced MEM (Gibco) supplemented with 2% fetal bovine serum (v/v), 1% glutamax, and 0.2% penicillin/streptomycin. All cells were routinely tested for mycoplasma every month and were confirmed negative.

Thereafter, cells were treated with iAs (as NaAsO_2_) in two ways. For “rapid” cell transformation, we sub-cultured and attached cells overnight and then treated cells with 2.0 µM until they reached ∼80% confluence. This process was repeated every 3 or 4 days for 12 weeks. We also transformed cells with a “two-hit” and time-dependent *in vitro* iAs exposure system recently described by Rea et al.^57^. In this case, cells were treated with freshly diluted 0.5 μM iAs at each cell passage for 17 weeks. At the end of 17 weeks, cells were then exposed to 2.0 μM iAs for another 10 weeks. The time-matched controls for the “second hit” were cells that continued to be exposed to 0.5 μM iAs. Every experiment with both exposure models included time-matched vehicular controls, and we monitored all cells for altered morphology throughout the exposure period. The specific exposure model used for each experiment is indicated in the corresponding figure legend.

### SATB2 and circ3915 siRNA knockdown

siRNAs used to knock down *SATB2* and *circ3915* RNA are itemized in **Table S9**. To knock down *SATB2* mRNA, we used pre-validated ON-TARGETplus siRNAs from Horizon. Three of these siRNAs did not overlap with any circ3915 sequence (probes SATB2-1, -2, and -3) and were therefore pooled for linear *SATB2* mRNA knockdown. The remaining ON-TARGET siRNA (SATB2- Circular+Linear) was used to simultaneously knock down *SATB2* and circ3915 RNAs (**Fig S8A**). We also used the circular RNA interactome resource (https://circinteractome.nia.nih.gov/index.html) to design three siRNAs specifically targeting the circ3915 back-splice junction (circ3915-1, -2, and -3). The ON-TARGET non-targeting siRNA was used as a control for all experiments.

### *SATB2* and *circ3915* shRNA knockdown

Pre-designed and validated shRNAs from Millipore Sigma were used for stable *SATB2* knock-down (**Table S10; Fig S8B**). The reported knockdown efficiency for the *SATB2* circular and linear probe (SATB2-2) is 84%, and our experimental knockdown efficiency was 72-82% (not shown). The circ3915 shRNA was based on the circ3915 siRNAs, above. Arsenic-transformed cells were stably transduced with each shRNA (lentivirus U5 promoter-shRNA; CMV-GFP vector) by the University of Michigan Vector Core Laboratory.

### *SATB2* and *circ3915* overexpression

Full-length *SATB2* mRNA was amplified by PCR from plasmids that were kindly provided by the Costa Lab^50^. PCR products were purified with a Monarch PCR and DNA Cleanup Kit, and assembled in the pcDNA3.1(+) vector using the NEBuilder® HiFi DNA Assembly Cloning Kit to generate SATB2 3xFLAG-tagged constructs with the FLAG tag at the SATB2 C-terminus. For circ3915, we cloned the spliced exons of circ3915 (*SATB2* exons 3-6) into the pcDNA3.1(+) CircRNA Mini Vector. This vector surrounds the insert with splice sites and flanking *Alu*-containing intronic sequences for efficient back-splicing^101^ (**Fig S8C**). Gibson assembly was then used to generate circ3915 constructs with the 3xFLAG-tag at the C-terminus of the putative circ3915 peptide. All tags on circular constructs were inserted downstream of the back-splice junction but upstream of putative start codons to ensure that tags were only translated from back-spliced transcripts. The sequence of all over-expression constructs were confirmed by Sanger sequencing.

### Transient transfection of human cell lines

For transient *SATB2* and *circ3915* overexpression, plasmids (or empty vectors) were transfected into non-treated BEAS-2B cells; for siRNA knockdown experiments, siRNAs were transfected into iAs-transformed BEAS-2B cells. In both cases, we transfected cells with Lipofectamine 3000 reagent in Opti-MEM™ I Reduced Serum Medium according to the manufacturer’s instructions. Cells were incubated for 24-48 hours after transfection and then collected for downstream analyses. RNA over-expression was confirmed by RT-qPCR and Nanostring, and protein expression was confirmed by immunoblots.

### Lentiviral production, transduction, and selection

For stable *SATB2* and *circ3915* overexpression from lentiviruses, we first replaced the SATB2 3xFLAG tag with GFP and replaced the circ3915 3xFLAG tag with HiBiT (**Fig S8D**). These plasmids were then subcloned into lentiviruses by the University of Michigan Vector Core. Briefly, lentivirus packaging vectors psPAX2 and pC1-VSVG were co-transfected with proviral plasmid at approximately a 1:1:2 ratio, respectively, using standard polyethylenimine (PEI) precipitation methods (MW 2500). Three µg PEI per 1 µg DNA were incubated in Opti-MEM at room temperature for 20 min, and the mixture was then added to fresh DMEM supplemented with 10% FBS (v/v). The DNA/PEI-containing media was then added to HEK-293T cells to propagate the lentivirus. Supernatants were collected after 72 hours and spun at 800 x g for 10 min to remove cellular debris. The viral supernatant was then concentrated on a 0.45 µm HV-Durapore Stericup 13,000 rpm on a Beckman Avanti J-E centrifuge at 4°C for 4 hrs and resuspended at 10X the original concentration in DMEM (∼1×10^7^ TU/mL). The concentrated lentivirus was stored in aliquots at -80°C until use.

Lentivirus transduction and FACS sorting were performed at the University of Michigan Vector Core and Flow Cytometry Core, respectively. Briefly, BEAS-2B cells were transduced with several lentivirus titers (between 0.5-1X; MOI ≤ 1) in multiple wells of a 6-well plate. After 48 hours, cells were either treated with 1 µg/mL puromycin for 5 days (or until non-transduced cells were dead), or subjected to GFP or RFP sorting on a Sony SH800 Cell Sorter.

### Split luciferase translation assay

BEAS-2B cells containing circ3915-HiBiT or empty vector were detached by incubating in 0.5% trypsin for 5 min, washed in PBS, and then re-suspended in PBS. Cells were diluted 1:10 in PBS, and an equal volume of trypan blue was added to the diluted cells. Cells were then loaded onto a slide and counted on a Countess II FL automated cell counter (Invitrogen). The concentration of the stock cell suspension was then calculated, and an appropriate amount of PBS was added to the stock cell suspension to achieve ∼10,000 cells per 100 µL. 100 µL of cell suspension (∼10,000 cells) was then added to replicate wells (n=6 per cell group) in a white opaque 96-well plate. An additional 6 wells were loaded with 100 µL of room temperature PBS to measure background luminescence. Next, an equal volume of Nano-Glo® Luciferase Assay Reagent was added to each well. To ensure complete lysis and sufficient mixing, the plate was shaken on an orbital shaker for 30 sec and then incubated for 10 min at room temperature. Relative luminosity (RLU) was measured on a Biotek Synergy HT Multi-Mode Microplate Reader with an integration time of 1.25 sec. The RLU from PBS-only wells was subtracted from the RLU values from cell-containing wells in the same row, and relative luminescence was calculated as the fold-change of circ3915-HiBiT expressing cells relative to empty vector cells. Each experiment was performed in biological triplicate.

### Luciferase cell proliferation assay

We used the CellTiter-Glo ATP Assay to measure cell proliferation 72 hours after seeding. Non-treated cells were the negative control for iAs-transformed and KRAS-G12V/TP53 BEAS-2B cells; iAs-transformed cells treated with non-targeting siRNAs were the negative control for iAs-transformed cells treated with siRNAs; and cells transfected with empty vector were the negative controls for cells stably over- expressing *SATB2*, circ3915, and circular + linear *SATB2* (see Results). Luminescence was quantified on a BioTek Synergy plate reader. Background luminescence was subtracted from the luminescence of the samples and background-corrected using the luminescence readings from cells that did not receive the knockdown treatment (above). Each experiment was performed three times with six technical replicates each.

### Trans-well migration assay

We used an *in vitro* trans-well assay to measure cell migratory potential. BEAS- 2B cells were grown in complete media (DMEM + 10% FBS + penicillin-streptomycin), and then starved in blocking media (DMEM + 0.1% BSA) for 24 hours. Starved cells were collected with trypsin, and ∼2 x 10^4^ cells (in 100 µL) were seeded on the upper compartment of each trans-well insert (Greiner Bio-One ThinCert, 8 µm pore) that had been treated for 30-60 min with a coating solution (1X PBS, 15 µg/mL collagen, 0.1N NaOH). The lower chamber contained 600 µL of a migration-inducing media (DMEM, 0.1% BSA, 20 ng/µl EGF). Cells were incubated for 24 hr, and those cells that had migrated through the insert were washed in 1X PBS, fixed in methanol for 15 min, stained for 15 min in (1X PBS, 1% TritonX-100, 10 μg/mL DAPI), and washed in 1XPBS. Stained cells were then imaged on an Olympus microscope at 40X magnification and counted using the cellSens counting module. To compare the effect of a specific treatment on cell migration, we normalized all cell counts to non-treated control cells.

### Total RNA extraction

Unless otherwise specified, total RNA was extracted from washed and pelleted cells using the Zymo Quick-RNA Miniprep Kit per the manufacturer’s protocol and eluted in RNase-free water. RNA concentrations were measured by spectrophotometry, and RNA integrity was assessed on an Agilent Bioanalyzer before downstream RT-qPCR, Nanostring, and RNA-seq analysis. Purified RNA was stored at -80C.

### Polysome fractionation and RNA extraction

Samples were prepared using previously published methods^102,103^, with some modifications to accommodate experimental needs. iAs-transformed BEAS-2B cells or BEAS-2B cells overexpressing the circ3915-HiBiT RNA were seeded in 15 cm^2^ plates (three replicate plates per condition) and grown to 70-80% confluency in DMEM + 10% FBS + penicillin-streptomycin (complete media). Cells were then treated with 1 mM puromycin for 3 hours, after which the media was replaced with complete media + 100 µg/mL cycloheximide for 5 min at 37°C. Cells were then washed twice with 10 mL of ice-cold PBS + 100 µg/mL cycloheximide and collected by centrifugation at 250 x g for 5 min at 4°C. Cell pellets were resuspended in 450 µL hypotonic buffer (140 mM NaCl, 1.5 mM MgCl_2_, 10 mM Tris pH 7.5, 0.5% NP40, 100 µg/mL cycloheximide, 1X HALT Protease Inhibitor Cocktail, 2.5 mM DTT, and 1 U/µL RNAseOut, and vortexed for 5 seconds. We then added 25 µL of 10% Triton X-100 and 25 µL of 10% sodium deoxycholate to the re-suspended cells. Nuclei were recovered by centrifugation at 16,000 x g for 7 min at 4°C. The supernatant was collected and kept on ice, and 10% of the total volume (50 µL) was immediately added to a 1.5 mL microfuge tube containing 750 µL TRIzol reagent and flash-frozen in liquid nitrogen. The OD was measured, and all cytoplasmic lysates were equilibrated by OD. Lysates were loaded dropwise onto 10 mL 5%-50% sucrose gradients, and the loaded sucrose gradients were centrifuged at a high speed (36,000 rpm) for 2 hours at 4°C using a SW41Ti rotor. Fractions were collected (at room temperature) from each gradient using a BioComp automated piston gradient fractionator. The resulting fractions were collected in 750 µL TRIzol, flash frozen in liquid nitrogen, and stored at -80°C until RNA extraction.

RNA was extracted from each fraction by first mixing 0.2 mL chloroform per 1 mL of TRIzol and separating the phases by centrifugation at 12000 x g for 15 min at 4°C. The aqueous phase (∼0.5 mL) was transferred to a new tube, mixed with 0.5 mL isopropanol, and incubated for 10 minutes at 4°C. Precipitated RNA was collected by centrifugation at 12000 x g at 4°C, washed twice with 75% ethanol, air-dried, and re-suspended 10-20 µL nuclease-free water. Purified RNA was then treated with DNAse I according to the manufacturer’s instructions. The distribution and abundance of *SATB2* and circ3915 RNA molecules across the polysome gradient were then analyzed by RT-qPCR (below).

### SATB2 and circ3915 RT-qPCR

T-qPCR primers are listed in **Table S11**. The circ3915 consists of four *SATB2* exons and is generated from *SATB2* mRNA by back-splicing exons 3 and 6 (**Fig S8E**). We, therefore, used this backsplice junction to specifically detect circ3915. RT-qPCR was performed on a BioRad CFX96 Real-Time system using a thermal cycling program consisting of [95°C for 5 min; and 40 cycles of 30 sec at 95°C, 30 sec at 60°C, and 10 sec at 72°C). We also instituted a melting curve step based on the instrument’s temperature gradient settings. C_t_ values were normalized against the total RNA input or the geometric mean of 3 housekeeping genes (GAPDH, RPII, and TBP), and normalized values were used to calculate fold-change relative to control samples.

### Microarray hybridization and analysis

Total RNA from each sample was quantified using a NanoDrop ND- 1000 (Thermo Fisher Scientific), and then sent to Arraystar (Rockville, MD) for circRNA analysis. Briefly, total RNA was first digested with RNase R to enrich for circRNAs. The enriched circRNAs were amplified and labeled using the Arraystar Super RNA labeling kit, purified with a Qiagen RNeasy Mini Kit, and quantified on a NanoDrop ND-100. Then, 1 µg of the amplified circRNAs were fragmented by adding 5 µL of 10X blocking agent and 1 µL of 25X fragmentation buffer and incubating at 60°C for 30 min. Fragmented RNAs were then diluted with 25 µL of 2X hybridization buffer, and the entire solution (50 µL) was added to an Arraystar Human Circular RNA Array (#AS-S-CR-H-V2.0). Microarrays were incubated for 17 hours at 65°C in an Agilent hybridization oven, washed, fixed, and then scanned on an Agilent G2505C scanner. The resulting images were processed with the Agilent feature extraction software (v11.0.1.1; Agilent Technology, Inc.). Expression analysis was conducted by applying quantile normalization and processing the normalized data with the R limma package (https://www.r-project.org/). A low-intensity filtering step was implemented, retaining only circRNAs surpassing a specific threshold. Differentially expressed circRNAs were defined as those exhibiting ≥1.2 fold-change and a significance level of *p* < 0.05. Volcano plots were generated by R Software R-3.3.1 gplots, and we used the heatmap2 function to generate hierarchical clusters and visualize variable circRNA-expression patterns among samples. Data conversion was conducted based on Z-score normalization.

### RNA-seq

Total RNA-seq libraries were prepared and sequenced by the Van Andel Genomics Core. Briefly, ribosomal RNAs were depleted from 500 ng of total RNA using a QIAseq FastSelect –rRNA HMR Kit. Libraries were then prepared with a KAPA RNA HyperPrep Kit, making sure to shear RNAs to the recommended 300-400 nt length before converting RNA to cDNA. cDNA fragments were ligated to IDT for Illumina TruSeq UD Indexed adapters, and indexed libraries were amplified via PCR. Library quality and quantity were then evaluated using Agilent DNA High Sensitivity chips, the Promega QuantiFluor® dsDNA System, and Kapa Illumina Library Quantification qPCR assays. The individually indexed libraries were then pooled, and 50 bp paired-end sequencing was performed on an Illumina NovaSeq6000 sequencer to an average depth of 150M raw paired reads per transcriptome. Base calling was done with Illumina RTA3, then demultiplexed and converted to FastQ format with Illumina Bcl2fastq v2.20.0.

Adapter sequences and low-quality bases were removed with TrimGalore v0.6.10 (https://github.com/FelixKrueger/TrimGalore) using the parameters ‘--paired -q 20’. Trimmed reads were aligned to the hg38 reference genome (GENCODE release 33)^104^ and counted using STAR v2.7.10a^105^, with the parameters ‘--twopassMode Basic --quantMode GeneCounts’. Raw counts were then converted to variance stabilizing transformation (VST) counts. Principal component analysis (PCA) was performed with the top 10,000 genes with the highest variance, using the vst() function in DESeq2 v1.38.3 with ‘blind=FALSE’^106^. We used DESeq2 and the design ‘∼ group’ to identify differentially expressed genes (DEGs). Pairwise contrasts were tested for significance at *p_adj_* ≤ 0.01. Log-fold changes were shrunk using the lfcShrink function with the parameters ‘type="ashr"’^107^. Gene set enrichment analysis (GSEA) was conducted using clusterProfiler v4.6.2^108^ with the parameter ‘eps = 0.0’. Genes were ranked based on the shrunken log fold changes and a significance cutoff of *p_adj_* ≤ 0.05. MSigDB annotations were retrieved using msigdbr v7.5.1. Summary plots were created with enrichPlot v1.18.4.

### Nanostring analyses

We used direct digital counting with custom Nanostring codesets (**Fig S8F, Table S12**) to verify differentially expressed circular (microarray) and linear (RNA-seq) transcripts. For circRNA expression, we designed capture and reporter probes for the backsplice junction of the top candidate circRNAs. In cases where circRNAs shared an exon at the backsplice junction, protector probes were designed to prevent cross-hybridization. For linear RNAs, capture and reporter probes were designed to target regions overlapping linear (consecutive) splice junctions. To detect linear transcripts arising from those genes that also produce circRNAs, probes were designed against regions that did not correspond with candidate circRNA sequences. We also included probes for 12 reference genes that are known to have high, medium, or low intrinsic expression levels. Total RNA isolated from experimental and control cells was subjected to nCounter™ SPRINT (Nanostring Technologies) analysis according to the manufacturer’s instructions. Background subtraction, normalization, quality control, and differential expression analysis were performed using the ROSALIND® cloud-based pipeline (https://rosalind.onramp.bio/). The ROSALIND pipeline generates read distribution percentages, violin plots, identity heatmaps, and sample MDS plots. Normalization, fold change, and significance were calculated using criteria provided by NanoString®. ROSALIND follows the nCounter protocol by dividing counts with the geometric mean of the internal normalizer probes within the same lanes. The reference probes selected for normalization are chosen based on the geNorm algorithm^109^. Fold changes and p-values were calculated using the fast method described in the nCounter® Advanced Analysis 2.0 User Manual.

### ATAC-seq

BEAS-2B cells (non-transformed; iAs-transformed; empty plasmid vector; and/or SATB2 over- expressing cells) were grown to 70-90% confluency, cryopreserved, and sent to ActiveMotif (Carlsbad, CA) for ATAC-seq. Briefly, cell pellets were resuspended in a lysis buffer, and nuclei were pelleted by centrifugation. Genomic DNA was recovered from ∼1x10^5^ nuclei, tagmented using an Illumina Nextera Library Prep Kit, and the resulting DNA fragments purified with a Qiagen MinElute PCR purification kit. Tagged DNA fragments were amplified through 10 PCR cycles, and amplified products were purified with Beckman Coulter Agencourt AMPure SPRI beads. Libraries were quantified using the KAPA Library Quantification Kit and then subjected to paired-end 42 (PE42) sequencing on an Illumina NextSeq 500 sequencer. Illumina base-call data was processed and demultiplexed with Bcl2fastq2 (v2.20). Adapter sequences were eliminated using skewer^110^ v.0.2.2, duplicates were removed with Samtools^111^ (v0.1.19), and then the remaining sequences aligned to the hg38 reference genome using the BWA algorithm^112^ (mem mode; default settings). Sequencing reads were processed as matched pairs, and only uniquely mapped reads with a mapping quality ≥ 1 were retained for subsequent analysis. To generate genome-wide coverage data, the alignments were extended *in silico* to a fixed length of 200 base pairs, and assigned to 32-base pair bins along the genome. Bigwig files were created using deeptools^113^ (v3.5.1).

For comparative analysis of chromatin accessibility, we normalized data by down-sampling the usable number of tags for each sample in a group to the level of the sample in the group with the fewest usable number of tags. Peaks were called using the MACS 2.1.0 algorithm^114^ without input control and with the ’--nomodel’ option, applying a cutoff of *p* ≤ 10^-7^. Any peaks that appeared in ENCODE false peak blacklists^115,116^ were removed, and a fraction of reads in peaks (FRIP) value ≥ 10% was considered good data quality. Signal maps and peak locations were used as input data for further analysis in the Active Motif (Carlsbad, CA) proprietary algorithm, which generates sample comparisons for peak metrics, peak locations, and gene annotations. Peak locations were used to annotate ENCODE-predicted cis-regulatory elements (cCRE). Peaks were annotated to the nearest gene using the annotatePeak function in ChIPseeker v1.34.1^117^ with the parameters ‘tssRegion = c(-3000, 500)’. We also used GenomicRanges v1.50.2^118^ to annotate peaks with at least a 1 bp overlap with ENCODE3^115^–predicted cis-regulatory elements. Differentially accessible peaks were selected based on specific peak metrics, including an absolute shrunken log2 fold change > 0 and *p_adj_* < 0.05.

For pathway enrichment analysis, a given ATAC peak was removed if the distance to the nearest transcription start site (TSS) was >100 kb or if it was annotated as ‘distal intergenic’ and did not overlap with the ENCODE cCRE promoter-like (PLS), proximal enhancer-like (pELS), or distal enhancer-like (dELS) regions. Peaks were grouped together if they were annotated to the same nearest gene. A given gene was considered up-regulated if there were more significantly up-regulated peaks than down, and *vice versa* for down-regulated genes. The resulting gene lists were input into the compareCluster function in clusterProfiler v4.6.2 with a default significance cutoff of *p_adj_* ≤ 0.05. We then used msigdbr v7.5.1 to retrieve MSigDB annotations. Dot plots of the enrichment results were created using enrichPlot v1.18.4.

### Protein extraction

Cells were grown in 75 cm^2^ cell culture flasks (Nest Scientific #708001) and detached in 0.25% Trypsin-EDTA (Thermo Fisher Scientific #25200072). Approximately 1 x 10^7^ cells were washed with Dulbecco’s Phosphate Buffered Saline (DPBS, Sigma-Aldrich # D8537) and resuspended in 0.5 mL of radioimmunoprecipitation assay (RIPA) lysis and extraction buffer (Thermo Fisher Scientific #J62885.AE) that was supplemented with 1 mM phenylmethylsulfonyl fluoride (PMSF, Sigma-Aldrich #10837091001), 1X protease inhibitor cocktail (EpiGentek #R-1101), 10 mM 3-methoxybenzylamine (3-MBZ, Sigma-Aldrich #159891), and 0.5 mM benzylamine (BZA, Sigma-Aldrich #185701). Cell suspensions were incubated on ice for 30 min, gently vortexed, sonicated for 6 cycles (10 sec on/10 sec off) in a Diagenode Bioruptor300 (Denville, NJ), and then centrifuged at 16,000 x g for 10 min at 4°C. The resulting supernatants were collected in a fresh tube and on ice, and quantified using a Pierce BCA Protein Assay Kit (#23225).

### Western blots

Proteins were separated on precast polyacrylamide gels (NuPAGE™ 4 to 12%, Bis-Tris, 1.0–1.5 mm, Mini Protein Gels-NP0321BOX), and transferred to iBlot™ 2 Transfer Stacks (Invitrogen #IB23001) using an iBlot 2 Gel Transfer Device (Invitrogen #IB21001). Membranes were then washed in 25 mL 1X TBS (50mM Tris-Cl, 150mM NaCl, pH 7.5) for 5 min at room temperature, and incubated for 1 hr at room temperature in 10 mL blocking buffer (1X TBST (Fisher BioReagents #BP337-500), 5% w/v nonfat dry milk). Membranes were then washed three times for 5 min each with 15 mL of TBST at room temperature, and incubated overnight at 4°C with gentle agitation in primary antibody mixed to desired concentration with primary antibody dilution buffer (1X TBST with 5% nonfat dry milk). Membranes were again washed three times in 15 mL TBST, and then incubated in 10 mL of blocking buffer containing the appropriate HRP-conjugated secondary antibody for 1 hr at room temperature. Primary antibodies targeted SATB2 (Invitrogen #PA5-83092; 0.1 mg/mL; diluted 1:2000); GAPDH (as a loading control; Abcam #ab9485; 0.94 mg/mL; diluted 1:3000); and anti-HiBiT (Promega #N72000; 1 mg/mL; diluted 1:1000). An HRP-conjugated IgG anti-mouse or anti-rabbit IgG (Proteintech SA00001-1 or SA00001-2; 0.2 mg/mL; diluted 1:100000) was used as a secondary antibody for chemiluminescent imaging. For chemiluminescence, we used a SuperSignal West Dura Extended Duration Substrate (ThermoFisher Scientific #34075) and the manufacturer’s recommendations to develop the signal. To detect HiBiT-tagged circ3915, we used the Nano-Glo HiBit Blotting system as per the manufacturer’s instructions (Promega, #N2410). In both cases, developed membranes were placed between transparent plastic sheets or plastic wrap, and images were acquired with a ChemiDoc MP imaging system (BioRad).

### Immunoprecipitation

Total protein lysates were generated from transfected BEAS-2B cells expressing FLAG-tagged SATB2 and/or circ3915, as described above. FLAG-tagged SATB2 and the circ3915peptide were then immunoprecipitated from 1 mg crude protein lysates using the FLAG Immunoprecipitation Kit according to the manufacturer’s protocols (Millipore Sigma #FLAGIPT1). Briefly, lysates were incubated at 4°C overnight with 40 µL of anti-FLAG M2 monoclonal antibody agarose affinity gel (anti-FLAG M2 affinity gel). The next day, the gel was washed four times with 1X wash buffer and five times with 1 mL of 25 mM ammonium bicarbonate. Agarose beads were then resuspended in 20 µL of sodium bicarbonate for mass spectrometry.

### Proteomics

Agarose beads (from above) were provided to the Michigan State University Integrated Mass Spectrometry Unit, where the ammonium bicarbonate buffer was removed and replaced with acetonitrile. Beads were briefly vortexed and incubated for 1 hr at room temperature, condensed with a magnetic bead stand, and the solvent was combined with the buffer. Samples were dried to completion at 30°C, resuspended in 50 µL of 50% acetonitrile + 25 mM ammonium bicarbonate, and digested overnight at 37°C with 1 μg of trypsin and 500 ng of LysC. Samples were again dried to completion at 30°C and resuspended in 50 µL 2% acetonitrile, 0.1% formic acid. Each analysis used 5 µL of sample and a 60-minute gradient, followed by 1 blank cycle. Samples were separated on a C18 EasySpray column (A = 0.1% formic acid, B = 0.1% formic acid in acetonitrile). The mass spectrometry analysis consists of one Full-scan MS followed by top 20 ddMS2. Database searching and protein identification were performed by Proteome Discoverer 2.2.0 via SEQUEST HT against the Uniprot Homo sapiens (UP000005460) database.

### Immunocytochemistry

BEAS-2B cells transfected with the SATB2-GFP and circ3915-HiBiT lentivirus were cultured as described above. Cells (∼1x10^5^) were seeded onto poly-L-lysine, collagen, or gelatin-coated coverslips in 12-well plates (Corning #3513), and incubated overnight at 37°C. The culture medium was removed, cells gently washed in 1X PBS at room temperature, and fixed in 4% paraformaldehyde for 15 min. Fixed cells were then incubated for 10 min at room temperature in 1X PBS + 0.5% Triton-X 100, washed in 1X PBS, blocked for 1 hr in 1X PBS containing 5% goat serum, and then incubated overnight at 4°C with mouse anti-HiBiT antibody (Promega #N7200; 1 µg/mL; diluted 1:1000) in 1X PBS containing 5% BSA. The next day, cells were washed three times in 1X PBS and then incubated in the dark for 1 hr at room temperature with anti-mouse Alexa Fluor 594 secondary antibody (Invitrogen #A-11005; 2 mg/mL; diluted 1:1500). Cells were washed 3 times in 1X PBS, stained with DAPI (Sigma-Aldrich #D9542; diluted 1:1000 in PBS) for 10 min at room temperature, and washed three times in 1X PBS. Coverslips were mounted onto glass slides with ProLong Diamond Antifade Mountant (Invitrogen #P36965), and images were captured with a Zeiss 880 confocal microscope (Zeiss Canada, Toronto, ON) and Olympus cellSens software. Immunofluorescence experiments were performed on two independent cell preparations.

## QUANTIFICATION AND STATISTICAL ANALYSIS

### Categorical data

Most categorical data were compiled in GraphPad Prism (v10.0), and pairwise comparisons were performed with a Student’s t-test or one-way ANOVA within GraphPad. Statistical details are found in the associated figure legends. Software and methods for analyzing RNAseq or ATAC-seq data are described in the Methods section above.

### Survival curves

De-identified patient data were retrieved from the TCGA database and included 150 entries from lung cancer patients who had a *KRAS* mutation and 342 entries from lung cancer patients who did not have a *KRAS* mutation. We used Kaplan–Meier curves^119^ to examine differences in categorical (quantile) *SATB2* expression levels and patient survival and to detect violations of the proportional hazards assumption. We used differential expression analysis (|Log_2_(FC)| > 1, p < 0.01) and the Cox Proportional Hazard method to control for continuous *SATB2* expression levels and age at diagnosis, gender, race (filtered for White and Black/African American individuals due to limited data availability for other groups), and smoking status (calculated in pack years). Models were limited to patients without KRAS mutations and were censored for either 2-year (N = 192 without a *KRAS* mutation, 100 with a *KRAS* mutation) or long-term survival (filtering for individuals who survived at least 2 years, and tracking them until the last follow-up date, N = 53 without a *KRAS* mutation, 16 with a *KRAS* mutation). We then used Cox proportional hazard models^120^ to assess the univariate prognostic significance of tumor variables on overall survival. Hazard ratios and P values were obtained from these models, where P values <0.05 were considered to indicate statistical significance.

## KEY RESOURCES TABLE

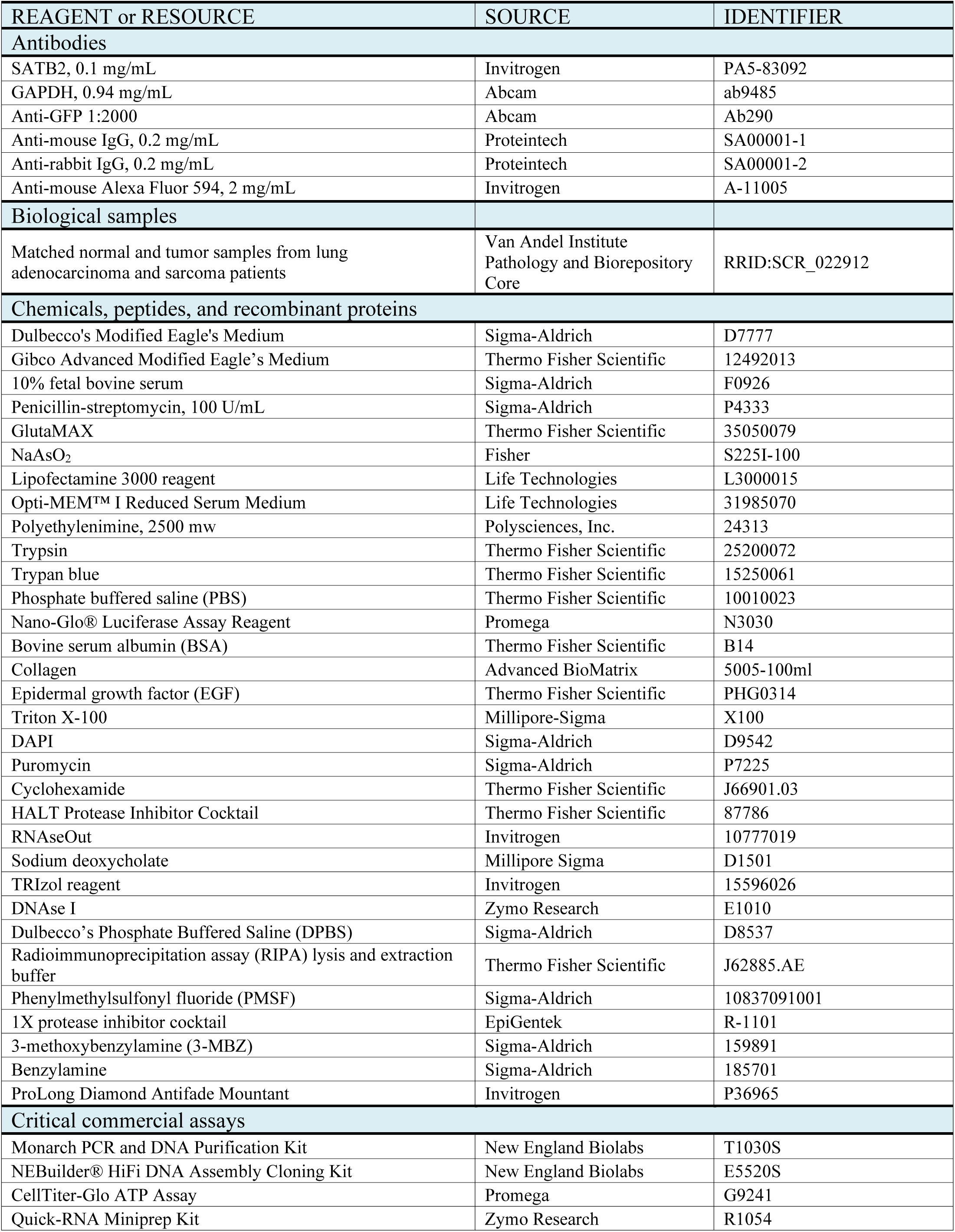
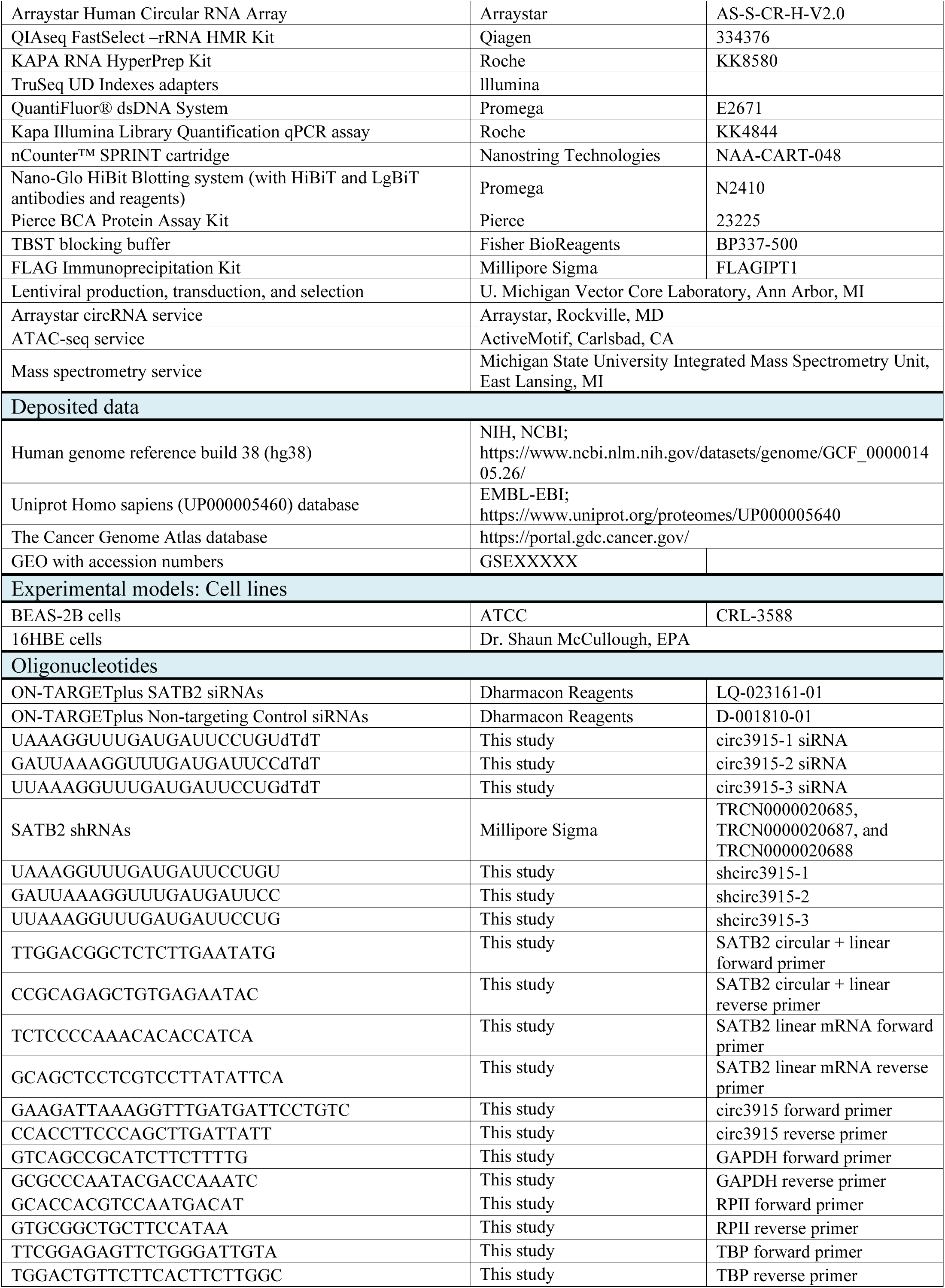
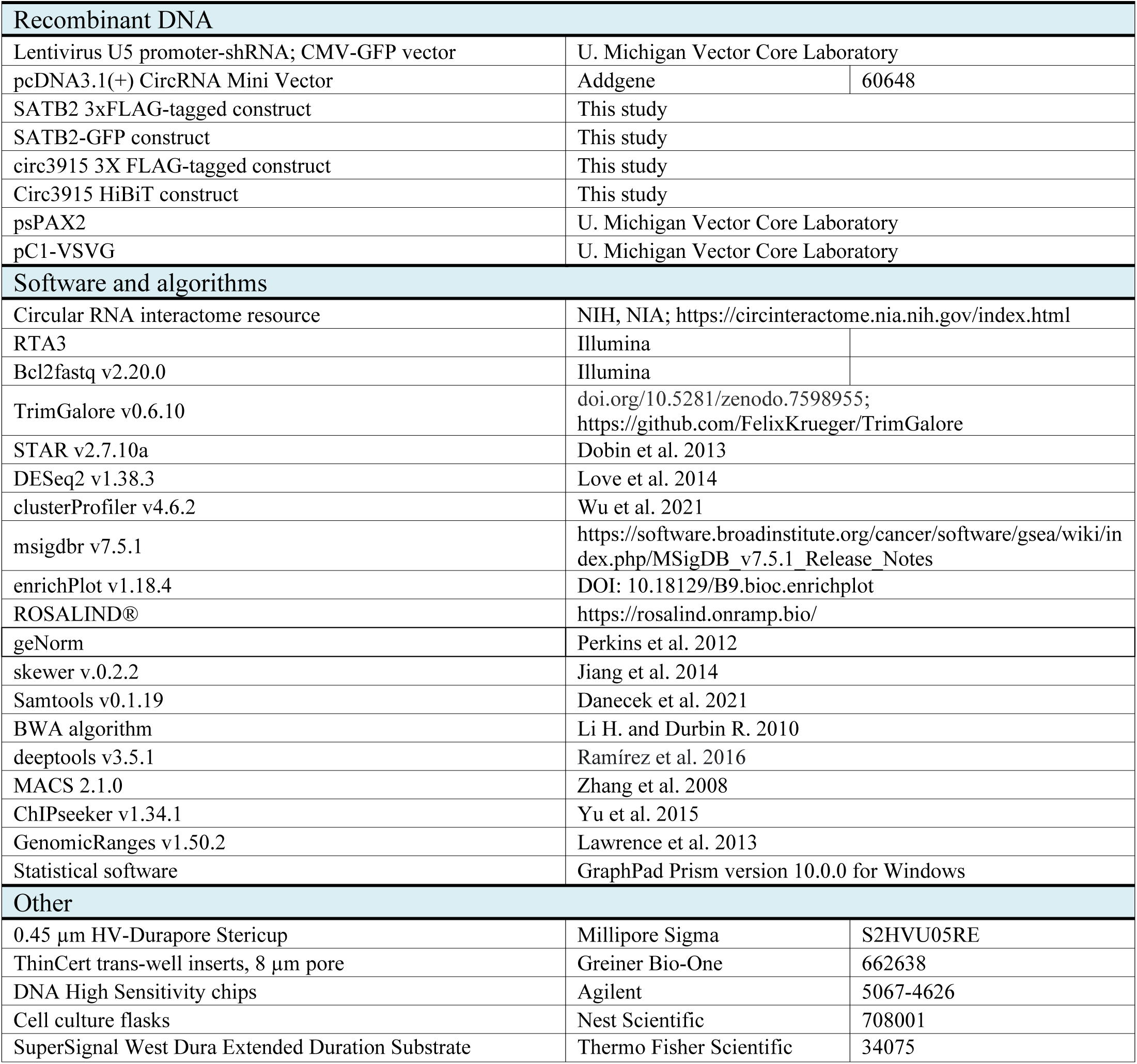

