## Supplemental Figures for "SATB2 and circ3915 RNA chromatin dysregulation drive *KRAS*-like oncogenic transformation"

Eleazer et al. 2024

**Fig S1**. Inorganic arsenic (iAs) exposure promotes transformation by driving oncogenic gene expression programs.

**Fig S2**. Inorganic arsenic induces global changes in chromatin accessibility.

**Fig S3**. circRNA detection in iAs-transformed cells.

**Fig S4**. Predicted interaction between SATB2 and circ3915p.

**Fig S5.** Differentially expressed genes (DEGs) in non-transformed BEAS-2B cells that over-express SATB2.

**Fig S6**. Differentially accessible regions (DARs) in non-transformed BEAS-2B cells that over-express SATB2.

**Fig S7.** Knocking down circ3915 or SATB2 in iAs-transformed BEAS-2B cells reverses oncogenic KRAS-like gene expression.

**Fig S8**. General experimental methods.

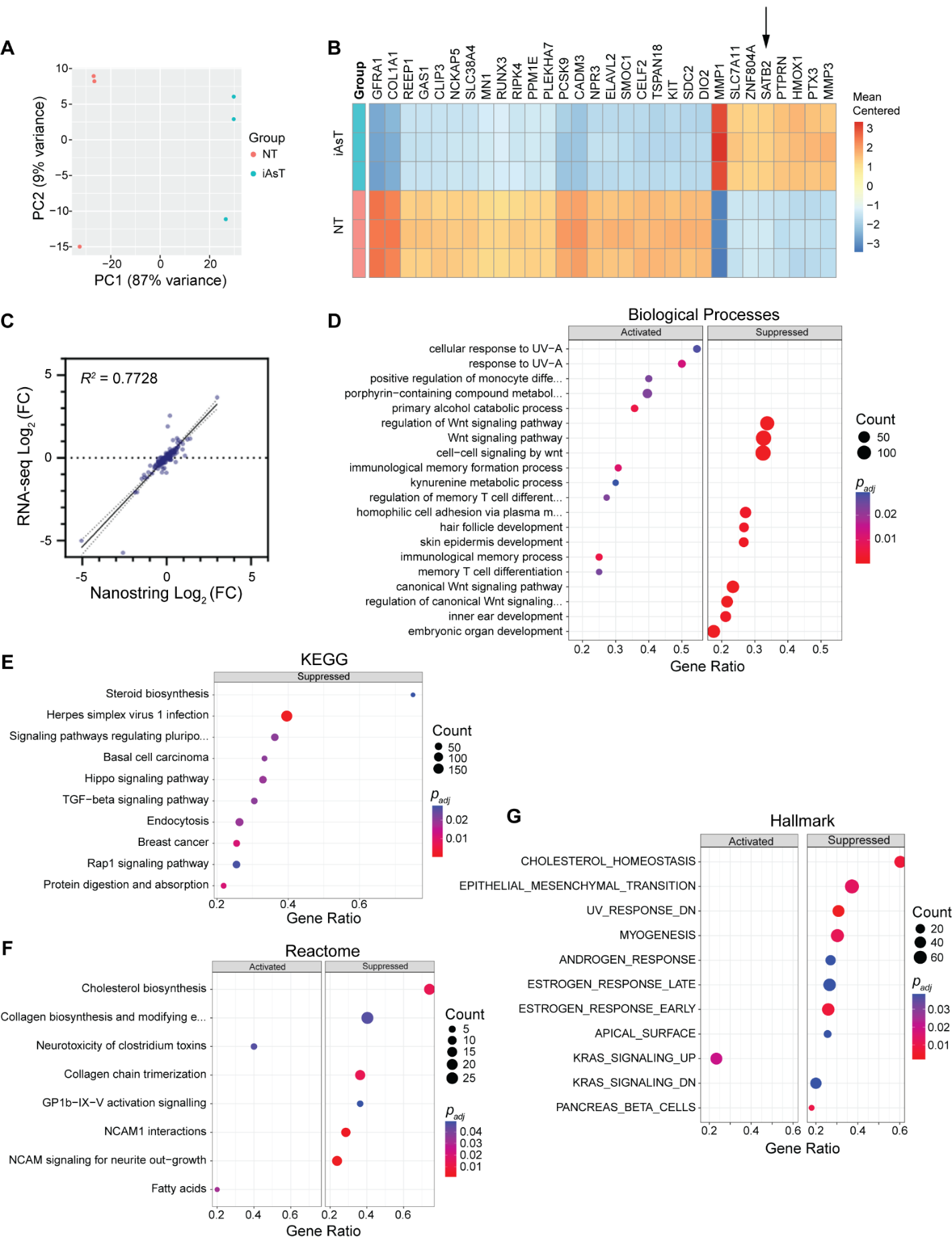

**Fig S1. Inorganic arsenic (iAs) exposure promotes transformation by driving oncogenic gene expression programs**. Data were generated from the rapid (12W) exposure model. **A**, Principal component analysis of RNA-seq expression profiles from non-transformed (NT, pink) and iAs-transformed BEAS-2B cells (iAsT, blue). iAs-transformed BEAS-2B cells were generated with the rapid (12 week) transformation model. **B**, The top 30 differentially expressed genes (DEGs) in iAs-transformed (iAsT) cells relative to non-transformed (NT) controls. **C**, Correlation between RNA-seq and Nanostring gene expression levels for 191 DEGs. Dashed lines = 95% confidence interval. **D**, The top enriched biological processes reflected in the iAsT DEGs. **E**, The top enriched KEGG pathways in the iAsT DEGs. **F**, The top enriched Reactome pathways in the iAsT DEGs. **G**, The top enriched MSigDB hallmarks in the iAsT DEGs.

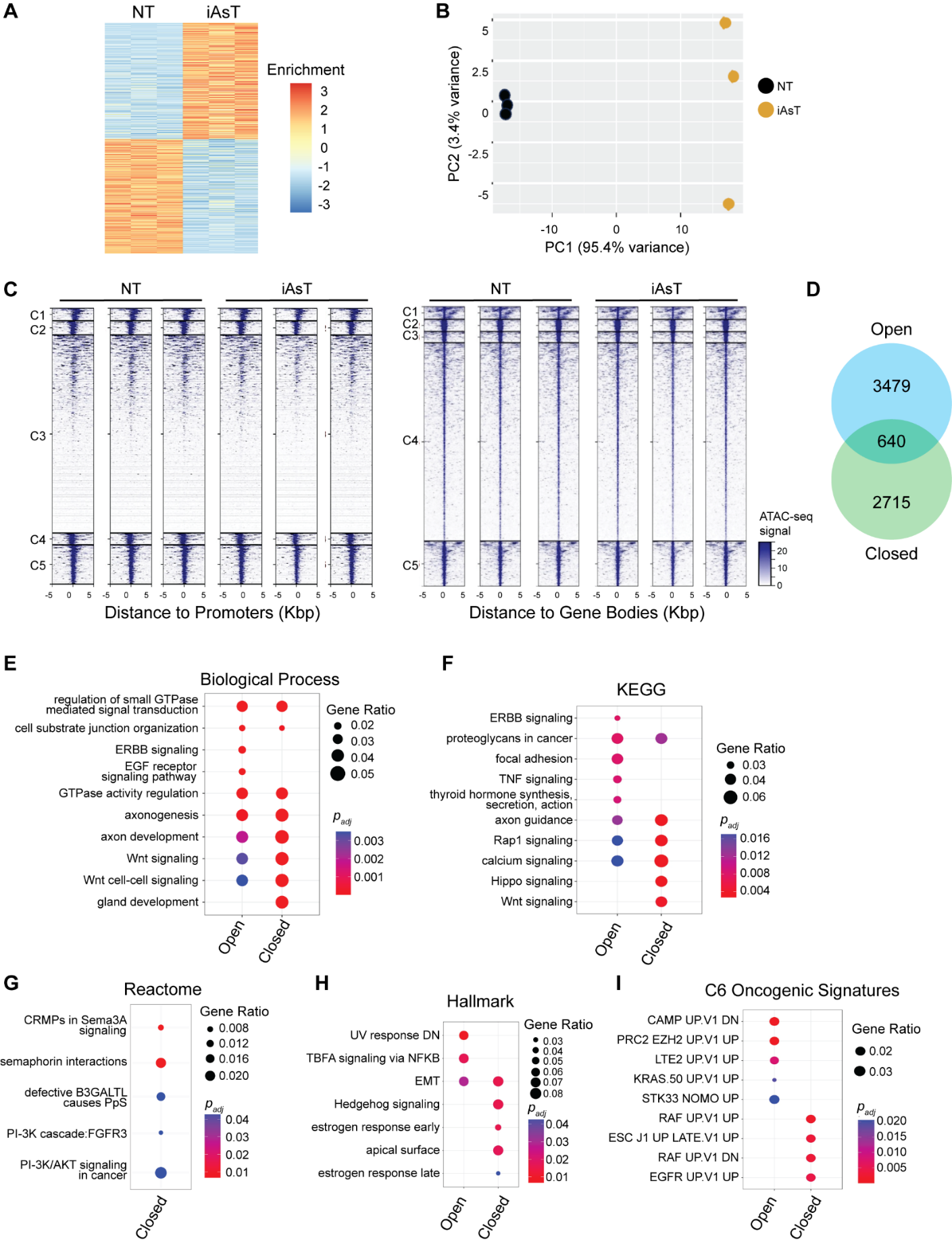

**Fig S2. Inorganic arsenic induces global changes in chromatin accessibility.** iAs-transformed cells were generated with the rapid (12 week) exposure model. **A,** DARs between non-transformed (NT) and iAs-transformed (iAsT) cells. **B**, Principal component analysis of differentially accessible regions in non-transformed (NT, black) and iAs-transformed (brown, iAsT) BEAS-2B cells. Transformed cells were generated with the rapid (12 week) transformation model. **C,** Genomic distance between ATAC-seq peaks and promoters (left) or gene bodies (right) in iAsT DARs. **D**, Congruence between genes associated with opened chromatin and closed chromatin in iAsT cells. **E**, The top enriched biological processes reflected in the iAsT DARs. **F**, The top enriched KEGG pathways in the iAsT DARs. **G**, The top enriched Reactome pathways in the iAsT DARs. **H**, The top enriched MSigDB hallmarks in the iAsT DARs. **I**, The top encriched C6 oncogenic signatures in the iAsT DARs.

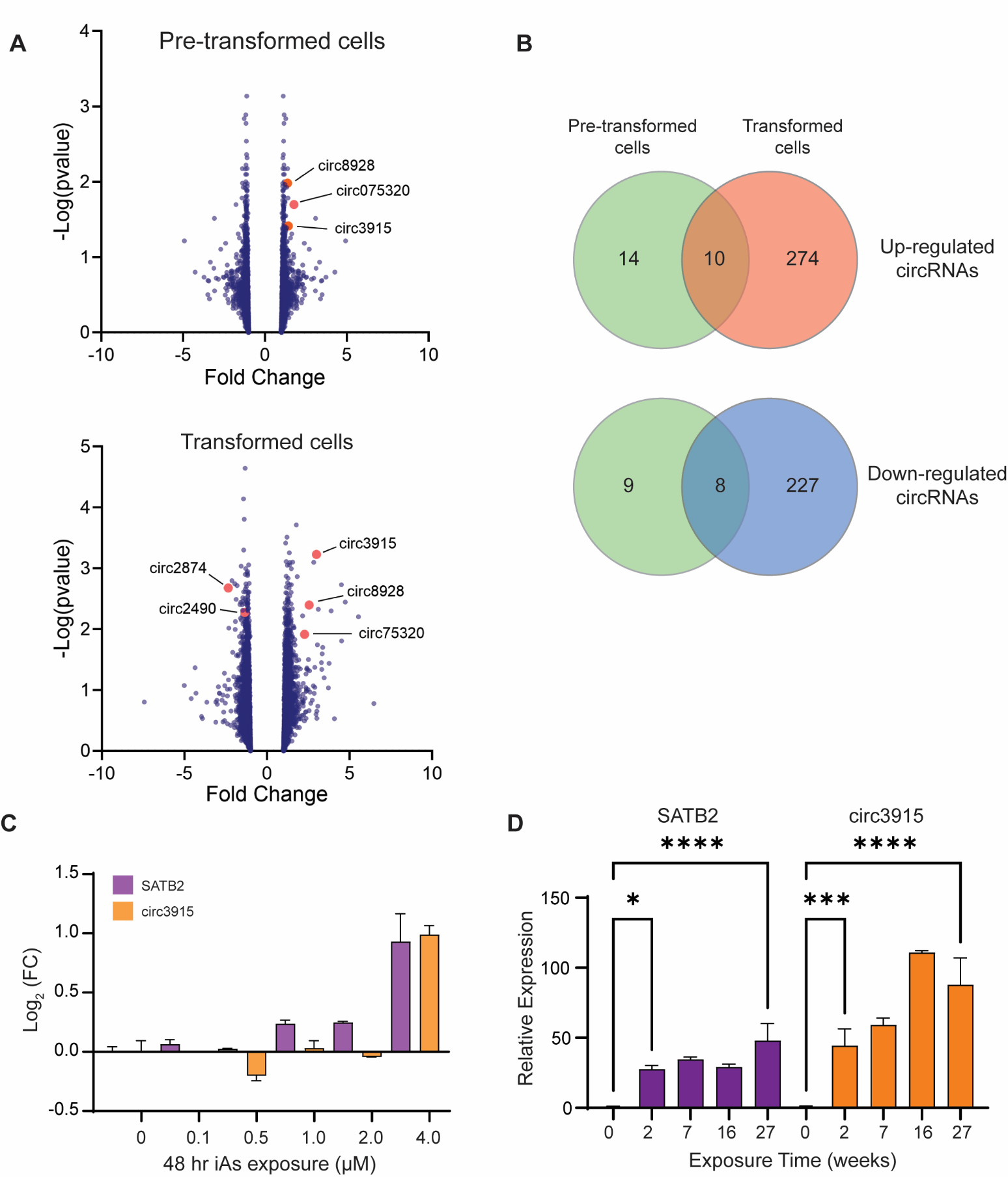

**Fig S3. circRNA detection in iAs-transformed cells.** **A**, Microarray-based detection of differential circRNA expression in pre-transformed (0.5 µM iAs for 17 weeks) and fully transformed (+ 2.0 µM iAs for another 10 weeks) BEAS-2B cells relative to non-treated cells. Up-regulated circRNAs were defined by *p* <0.05, FC >1.2. Down-regulated circRNAs were defined by *p* <0.05, FC <-1.2. **B**, Overlapping circRNA expression in pre-transformed and fully transformed BEAS-2B cells, as measured by Arraystar microarray. **C**, SATB2 mRNA and circ3915 expression in 16HBE cells after 48 hr exposure to increasing concentrations of iAs. **D**, SATB2 mRNA and circ3915 expression in 16HBE cells in the two-hit exposure model.

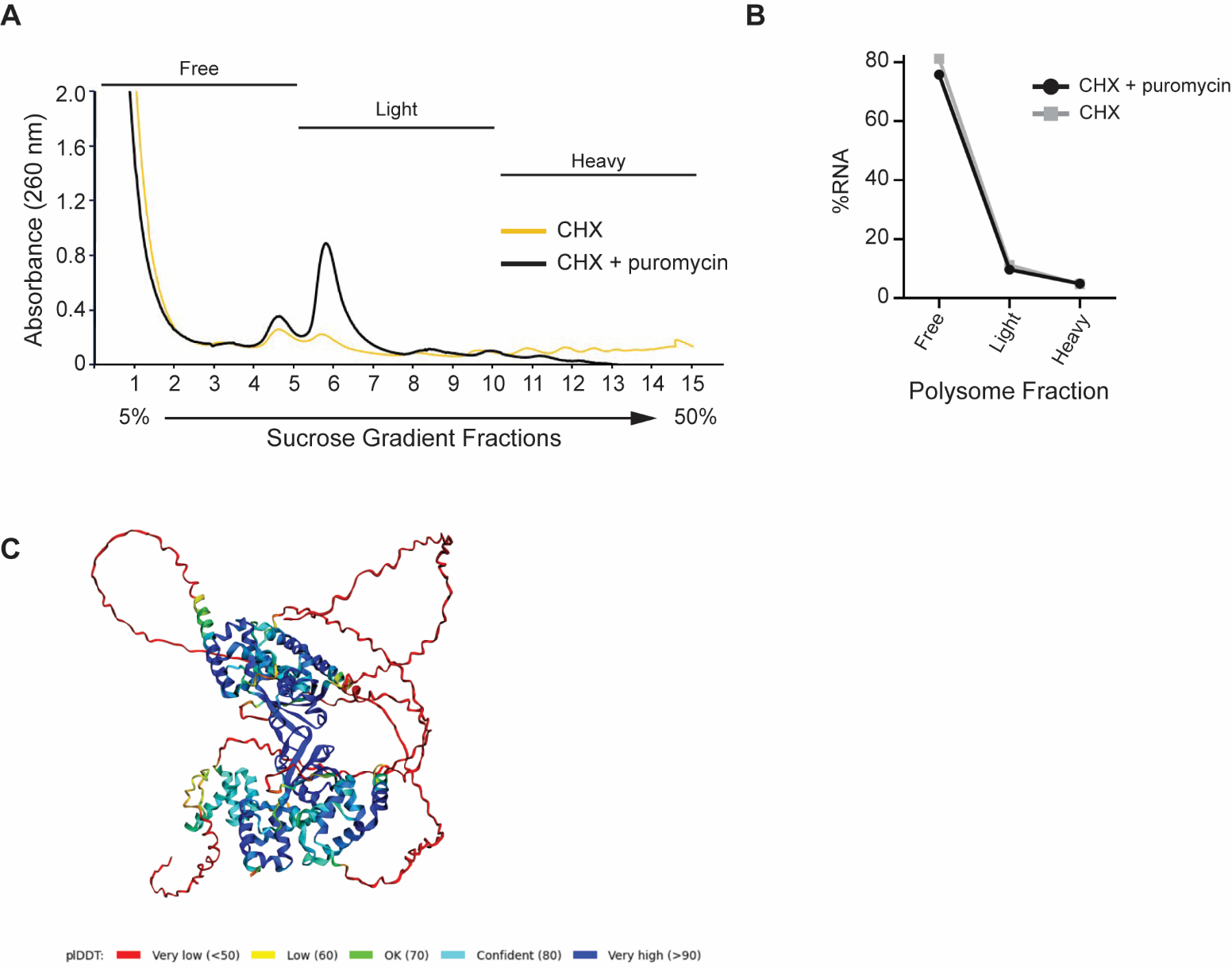

**Fig S4**. **Predicted interactions between SATB2 and circ3915p. A**, Puromycin treatment shifts circ3915 translation towards light and free polysomes. CHX = cycloheximide. **B**, In contrast, a known, untranslated circRNA (hsa_circ_0008928) does not associate with polysomes and does not show a shift in polysome association when treated with puromycin. **C**, Predicted structure for the SATB2-circ3915p complex. The SATB2 (UniProt accession Q9UPW6-1) and circ3915p amino acid sequences were submitted to the Alphafold2 Multimer tool using a cloud-based Colab notebook. The resulting complex was visualized with PyMOL, where color coding indicates the predicted Local Distance Difference Test (plDDT), or the quality of the predicted interactions. Disordered regions in both proteins are reflected in red (low-quality interactions), while structured regions in each protein are reflected in blue. The structured region closest to the viewer is circ3915p, and the other structured region is SATB2. The highest-confidence interactions between SATB2 and circ3915p (blue shading) occur in the SATB2 N-terminal ULD.

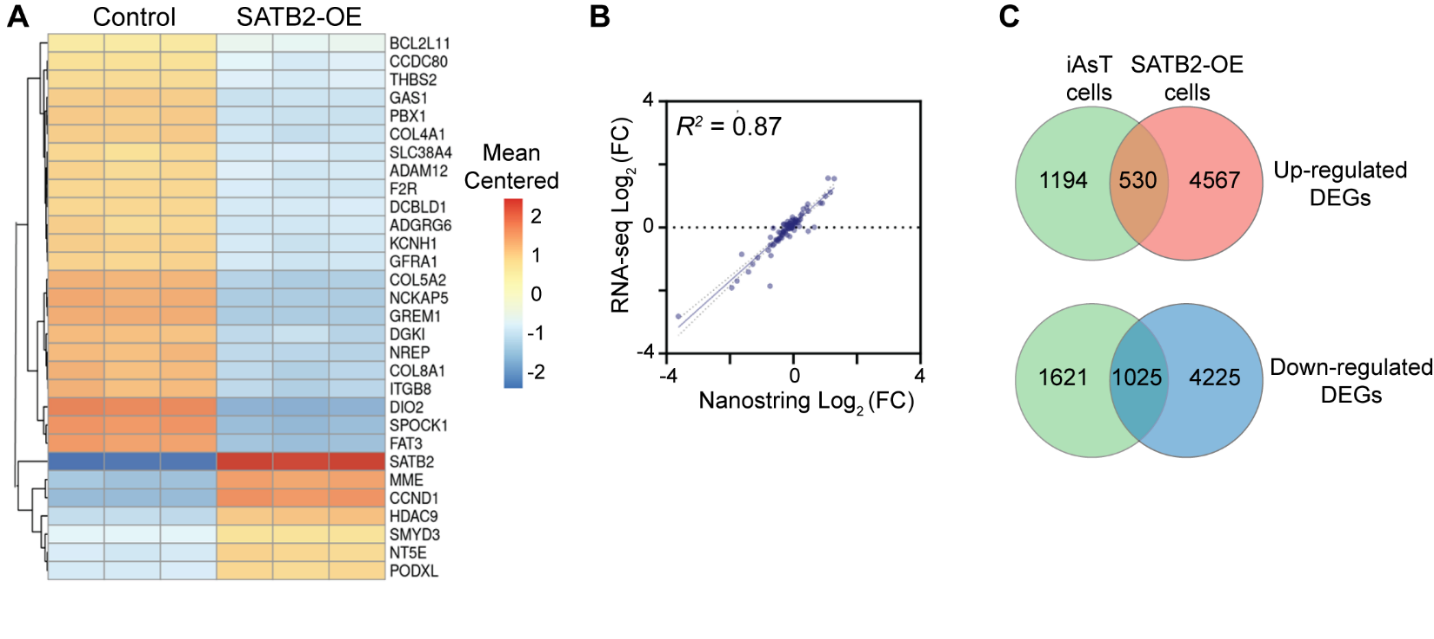

**Fig S5. Differentially expressed genes (DEGs) in non-transformed BEAS-2B cells that over-express *SATB2*.** **A**, Top 30 differentially expressed genes (DEGs) in non-transformed BEAS-2B cells that over-express *SATB2* (SATB2-OE). Control BEAS-2B cells contained an empty expression vector. **B**, Correlation between RNA-seq and Nanostring gene expression levels for 191 DEGs in non-transformed BEAS-2B cells that over-express *SATB2*. Dashed lines = 95% confidence interval. **C,** Overlapping up- and down-regulated genes in iAs-transformed (iAsT) BEAS-2B cells and non-transformed BEAS-2B cells with forced over-expression of SATB2. iAsT cells were generated with the rapid (12 week) transformation model.

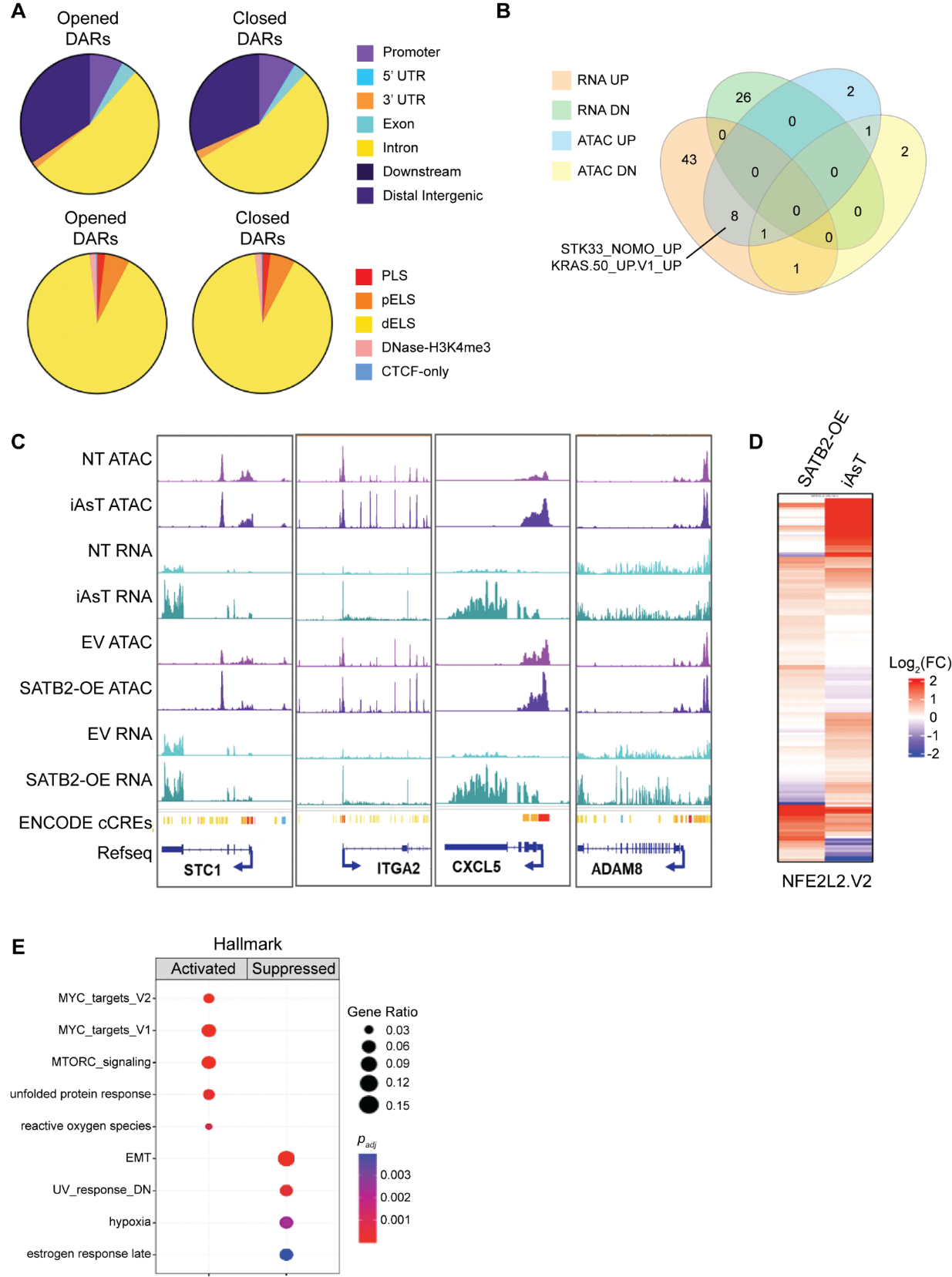

**Fig S6. Differentially accessible regions (DARs) in non-transformed BEAS-2B cells that over-express *SATB2*.** **A**, Genomic distribution of DARs (top) and ENCODE candidate cis-regulatory elements (cCREs; bottom) in non-transformed BEAS-2B cells with SATB2 over-expression. DARs were defined by FDR <0.01 and Shrunken Log_2_(FC) >1 for opened DARs, and Shrunken Log_2_(FC) <-1 for closed DARs. PLS = promoter-like signatures; pELS = proximal enhancer-like signatures; dELS = distal enhancer-like sequences. **B**, MSigDB oncogenic signatures in non-transformed BEAS-2B cells over-expressing SATB2 that were commonly enriched in DARs and DEGs. ATAC UP = opened chromatin; ATAC DN = closed chromatin; RNA UP = up-regulated transcripts; RNA DN = down-regulated transcripts. **C**, SATB2 opens chromatin at KRAS signature genes, increasing their gene expression. Shown is an integrative genomics viewer (IGV) snapshot of normalized average peak distributions for KRAS_UP_UP signature genes. NT = non-transformed BEAS-2B cells. iAsT = iAs-transformed BEAS-2B cells, generated with the rapid (12 week) transformation model; EV = non-transformed BEAS-2B cells containingg and empty expression vector; SATB2-OE = non-transformed BEAS-2B cells over-expressing SATB2. Encode cCREs are shown in yellow (dELSs), orange (pRLSs), and red (PLS). **D**, RNA expression of 161 genes from the MSigDB oncogenic NFE2L2 signature. **E**, Top enriched MSigDB Hallmark pathways in non-transformed BEAS-2B cells that over-express SATB2.

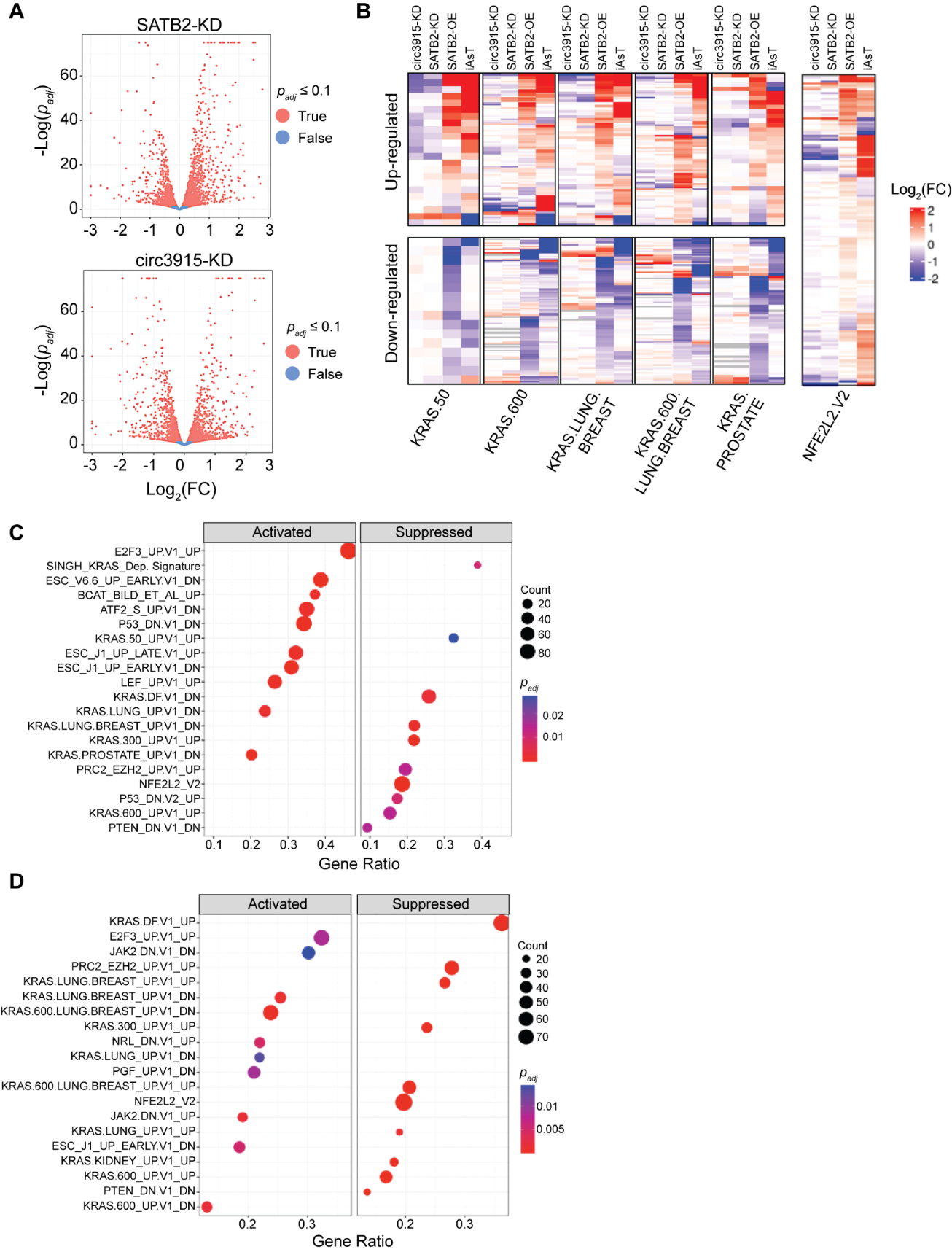

**Fig S7. Knocking down circ3915 or *SATB2* in iAs-transformed BEAS-2B cells reverses oncogenic KRAS-like gene expression. A**, Differentially expressed genes in iAs-transformed cells with stable SATB2 (SATB2-KD) or circ3915 (circ3915-KD) knockdown. **B**, *SATB2* (shSATB2) and circ3915 (shcirc3915) knockdown reverses MSigDB KRAS-like oncogenic signatures seen in iAs-transformed (iAsT) BEAS-2B cells (rapid transformation model) and non-transformed BEAS-2B cells that over-express SATB2 (SATB2-OE). Each row represents a KRAS signature gene from the indicated MSigDB oncogenic signature. **C** and **D**, Dot plots of GSEA-enriched MsigDB oncogenic signature gene sets in differentially expressed genes (*p_adj_* < 0.05) from stable *SATB2* (**C**) or circ3915 (**D**) knockdown in iAs-transformed cells show that KRAS-like and NFE2L2 MSigDB oncogenic gene expression signatures are reversed (suppressed).

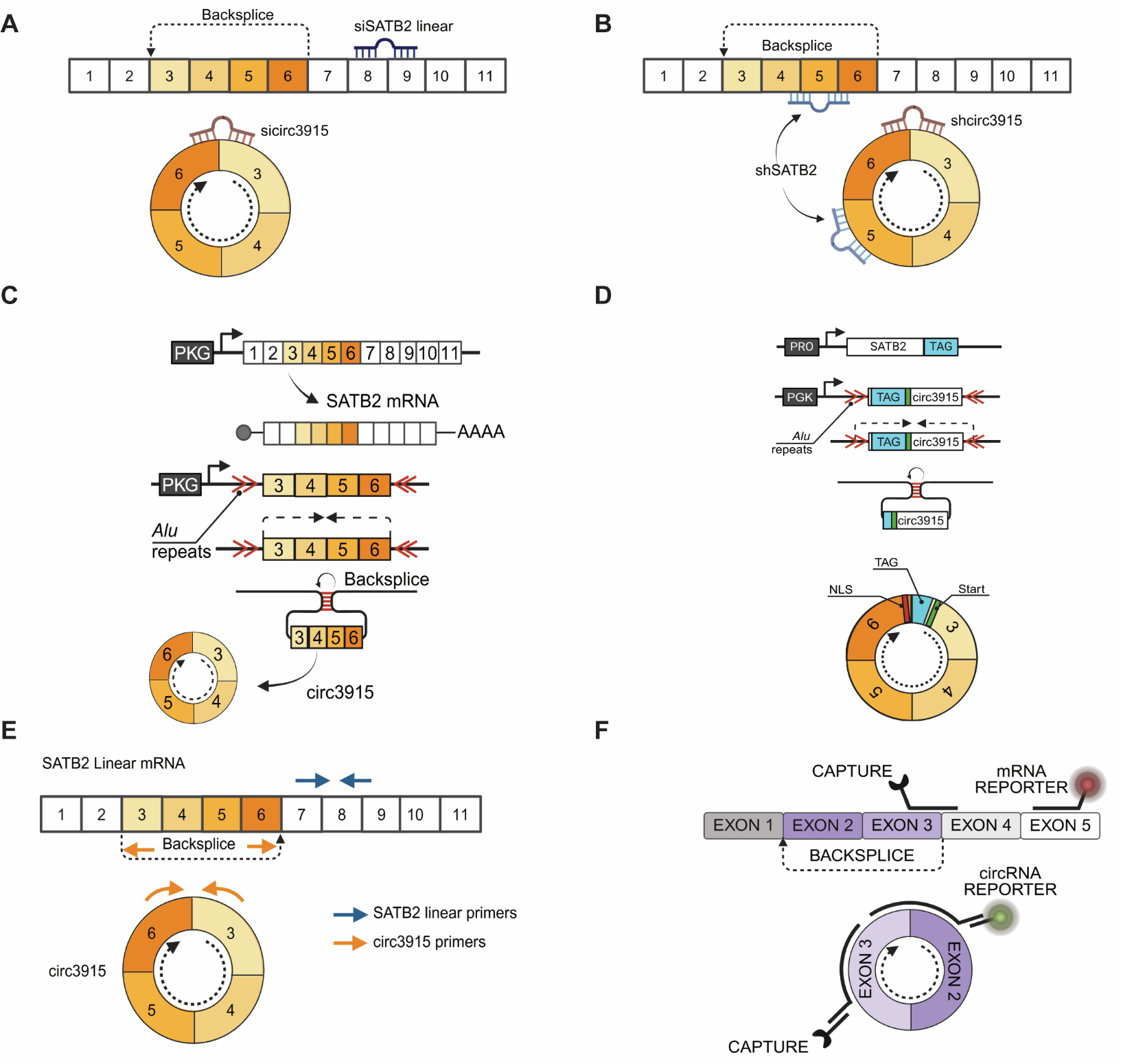

**Fig S8. General experimental methods. A**, General location of ON-TARGETplus small interfering (si) RNAs for transiently knocking down *SATB2 mRNA* or circ3915. Numbers correspond to *SATB2* exons. C+L = probe for circular and linear RNA. **B**, General location of Millipore Sigma short hairpin (sh) RNAs for stably knocking down *SATB2* mRNA or circ3915. **C**, Cloning strategy for over-expressing the linear *SATB2* mRNA (top) or circ3915 (bottom) from the pcDNA3.1(+) vector in non-transformed BEAS-2B cells. Once transcribed, the inverted *Alu* repeats in the pre-mRNA base pair, forming a loop that promotes backsplicing. **D**, Position of the fluorescent tag (HiBiT, FLAG, or GFP) in the lentiviral circ3915 overexpression vector. The fluorescent tag is placed upstream of the putative start codon to ensure that any peptide is derived from the backspliced transcript. **E**, RT-qPCR primer design strategy for specifically detecting linear *SATB2* mRNA, circ3915, or both transcripts. **F**, Nanostring probe design for detecting linear (top, red fluor) and circular (bottom, green fluor) RNAs. RNA isoforms are counted with complementary capture and reporter probes designed to hybridize the linear splice and circular backsplice junctions. RNA isoforms were counted via a unique barcode assigned to each reporter probe.
