## Supplemental Tables S4 and S8-S12 for "SATB2 and circ3915 RNA chromatin dysregulation drive *KRAS*-like oncogenic transformation"

**Supplemental Tables S4, S8-S12**

Eleazer et al. 2024.

**Table S4.** Proteins that interact with SATB2 and/or circ3915p, as determined by immunoprecipitation (IP)-mass spectrometry.

| Protein | GENE ID | SATB2  IP | EV IP | circ3915p  IP | Coverage  [%] | # Peptides |
| --- | --- | --- | --- | --- | --- | --- |
| DNA-binding protein SATB2 | *SATB2* | + | - | + | 69 | 53 |
| Unconventional myosin-Ic | *MYO1C* | + | - | + | 57 | 75 |
| Barrier-to-autointegration factor | *BANF1* | + | - | + | 43 | 2 |
| Histone H2A.Z | *H2AZ1* | + | - | + | 31 | 4 |
| Histone H1.1 | *H1-1* | + | - | + | 27 | 7 |
| Peroxiredoxin-2 | *PRDX2* | + | - | + | 27 | 4 |
| Small nuclear ribonucleoprotein Sm D2 | *SNRPD2* | + | - | + | 25 | 2 |
| 60S ribomal protein L35a | *RPL35A* | + | - | + | 23 | 3 |
| Serine/arginine-rich splicing factor 3 | *SRSF3* | + | - | + | 21 | 3 |
| Tubulin beta-6 chain | *TUBB6* | + | - | + | 20 | 6 |
| Core histone macro-H2A.1 | *MACROH2A1* | + | - | + | 16 | 4 |
| Stress-70 protein, mitochondrial | *HSPA9* | + | - | + | 15 | 6 |
| Isoform Long of Proteasome subunit alpha type-1 | *PSMA1* | + | - | + | 15 | 3 |
| Small nuclear ribonucleoprotein Sm D3 | *SNRPD3* | + | - | + | 15 | 2 |
| Isoform 2 of Nucleoside diphosphate kinase A | *NME1* | + | - | + | 12 | 2 |
| Isoform 2 of Eukaryotic translation initiation factor 5A-1 | *EIF5A* | + | - | + | 10 | 2 |
| Adenosylhomocysteinase | *AHCY* | + | - | + | 9 | 3 |
| Transformer-2 protein homolog beta | *TRA2B* | + | - | + | 9 | 2 |
| Probable ATP-dependent RNA helicase DDX17 | *DDX17* | + | - | + | 8 | 6 |
| Bystin | *BYSL* | + | - | + | 7 | 2 |
| Isoform SM-B1 of Small nuclear ribonucleoprotein-associated proteins B and B' | *SNRPB* | + | - | + | 5 | 2 |
| CCAAT/enhancer-binding protein beta | *CEBPB* | + | - | + | 4 | 2 |
| DNA replication licensing factor MCM7 | *MCM7* | + | - | + | 4 | 2 |
| Exportin-2 | *CSE1L* | + | - | + | 4 | 2 |
| Serine/arginine-rich splicing factor 6 | *SRSF6* | + | - | + | 4 | 2 |
| Histone deacetylase 2 | *HDAC2* | + | - | + | 4 | 2 |
| Glutamine--tRNA ligase | *QARS1* | + | - | + | 3 | 2 |
| ATP-dependent 6-phosphofructokinase, platelet type | *PFKP* | + | - | + | 2 | 2 |
| Helicase MOV-10 | *MOV10* | + | - | + | 2 | 2 |
| Histone H2AX | *H2AX* | + | - | - | 55 | 6 |
| Isoform Beta-2 of Protein phosphatase 1B | *PPM1B* | + | - | - | 53 | 17 |
| Isoform 2 of Plectin | *PLEC* | + | - | - | 43 | 185 |
| Y-box-binding protein 3 | *YBX3* | + | - | - | 19 | 10 |
| E3 ubiquitin-protein ligase TRIM47 | *TRIM47* | + | - | - | 16 | 6 |
| Ras-related protein Rab-10 | *RAB10* | + | - | - | 11 | 2 |
| Isoform 4 of 6-phosphofructo-2-kinase/fructose-2,6-bisphosphatase 3 | *PFKFB3* | + | - | - | 9 | 4 |
| Serine/threonine-protein phosphatase 2A 65 kDa regulatory subunit A alpha isoform | *PPP2R1A* | + | - | - | 8 | 4 |
| Serine/arginine-rich splicing factor 9 | *SRSF9* | + | - | - | 8 | 2 |
| EH domain-containing protein 2 | *EHD2* | + | - | - | 7 | 4 |
| 6-phosphofructo-2-kinase/fructose-2,6-bisphosphatase 2 | *PFKFB2* | + | - | - | 7 | 3 |
| FACT complex subunit SSRP1 | *SSRP1* | + | - | - | 6 | 4 |
| Erlin-1 | *ERLIN1* | + | - | - | 6 | 2 |
| Very-long-chain enoyl-CoA reductase | *TECR* | + | - | - | 6 | 2 |
| Sulfide:quinone oxidoreductase, mitochondrial | *SQOR* | + | - | - | 6 | 2 |
| Rho guanine nucleotide exchange factor 10 | *ARHGEF10* | + | - | - | 4 | 4 |
| Isoform 2 of ATP-dependent RNA helicase DHX30 | *DHX30* | + | - | - | 4 | 4 |
| Isoform 3 of Sodium/potassium-transporting ATPase subunit alpha-3 | *ATP1A3* | + | - | - | 4 | 2 |
| Peroxiredoxin-6 | *PRDX6* | + | - | - | 4 | 2 |
| Scaffold attachment factor B2 | *SAFB2* | + | - | - | 4 | 2 |
| D-3-phosphoglycerate dehydrogenase | *PHGDH* | + | - | - | 3 | 2 |
| Poly [ADP-ribose] polymerase 1 | *PARP1* | + | - | - | 3 | 2 |
| Girdin Acetyl [N-Term] | *CCDC88A* | + | - | - | 2 | 2 |
| DNA-dependent protein kinase catalytic subunit | *PRKDC* | + | - | - | 1 | 4 |
| Galectin-7 | *LGALS7* | - | - | + | 74 | 10 |
| Protein S100-A9 | *S100A9* | - | - | + | 61 | 7 |
| Serpin B3 Acetyl [N-Term] | *SERPINB3* | - | - | + | 41 | 11 |
| Tubulin beta-2A chain | *TUBB2A* | - | - | + | 40 | 14 |
| Protein S100-A8 | *S100A8* | - | - | + | 31 | 3 |
| Tubulin alpha-4A chain | *TUBA4A* | - | - | + | 28 | 11 |
| Junction plakoglobin | *JUP* | - | - | + | 25 | 17 |
| Fatty acid-binding protein 5 | *FABP5* | - | - | + | 24 | 5 |
| Desmoplakin | *DSP* | - | - | + | 21 | 54 |
| Isoform 2 of Serpin B12 | *SERPINB12* | - | - | + | 17 | 4 |
| 14-3-3 protein sigma | *SFN* | - | - | + | 12 | 2 |
| Desmoglein-1 | *DSG1* | - | - | + | 11 | 7 |
| Isoform 3 of L-lactate dehydrogenase A chain | *LDHA* | - | - | + | 10 | 3 |
| Heterogeneous nuclear ribonucleoprotein F | *HNRNPF* | - | - | + | 10 | 3 |
| Triosephosphate isomerase | *TPI1* | - | - | + | 9 | 2 |
| 14-3-3 protein zeta/delta | *YWHAZ* | - | - | + | 9 | 2 |
| Plakophilin-1 | *PKP1* | - | - | + | 9 | 5 |
| Serpin B13 | *SERPINB13* | - | - | + | 8 | 2 |
| Epiplakin | *EPPK1* | - | - | + | 7 | 7 |
| Desmocollin-1 | *DSC1* | - | - | + | 5 | 3 |
| Isoform 3 of Keratin, type II cytoskeletal 80 | *KRT80* | - | - | + | 5 | 3 |
| Isoform 1 of Protein POF1B | *POF1B* | - | - | + | 4 | 2 |
| Filaggrin | *FLG* | - | - | + | 1 | 2 |

**Table S8**. Enriched oncogenic signatures in non-transformed BEAS-2B cells with forced SATB2 over-expression.

| Signature ID | Set Size | Enrichment Score | NES | p_adj_ | q value |
| --- | --- | --- | --- | --- | --- |
| CAMP_UP.V1_UP | 199 | 0.50263699 | 2.10399852 | 2.5685E-08 | 1.3018E-08 |
| TBK1.DF_UP | 283 | 0.44755769 | 1.94364446 | 2.5685E-08 | 1.3018E-08 |
| CSR_EARLY_UP.V1_UP | 151 | 0.52377049 | 2.16623157 | 1.3815E-07 | 7.0018E-08 |
| MTOR_UP.N4.V1_UP | 195 | 0.47607561 | 1.99952005 | 3.5347E-07 | 1.7915E-07 |
| BMI1_DN_MEL18_DN.V1_UP | 145 | 0.48913375 | 2.01105242 | 1.4807E-05 | 7.5044E-06 |
| MEL18_DN.V1_UP | 140 | 0.47792898 | 1.96668321 | 2.429E-05 | 1.231E-05 |
| BMI1_DN.V1_UP | 147 | 0.45518038 | 1.89869029 | 4.0517E-05 | 2.0535E-05 |
| AKT_UP.V1_UP | 166 | 0.4369931 | 1.7845724 | 4.0517E-05 | 2.0535E-05 |
| STK33_NOMO_UP | 286 | 0.36930822 | 1.60129346 | 4.0517E-05 | 2.0535E-05 |
| KRAS.DF.V1_UP | 189 | 0.4136183 | 1.73122988 | 8.2599E-05 | 4.1863E-05 |
| MTOR_UP.V1_UP | 169 | 0.4189456 | 1.70995885 | 0.00018337 | 9.2938E-05 |
| STK33_UP | 281 | 0.36556656 | 1.57469259 | 0.00018337 | 9.2938E-05 |
| KRAS.600.LUNG.BREAST_UP.V1_DN | 277 | -0.4975731 | -1.5912219 | 0.00022578 | 0.00011443 |
| LTE2_UP.V1_DN | 195 | 0.40139451 | 1.68585905 | 0.0002917 | 0.00014784 |
| RB_P107_DN.V1_DN | 126 | 0.44834815 | 1.81394815 | 0.00031019 | 0.00015721 |
| E2F1_UP.V1_DN | 187 | 0.39297895 | 1.63424821 | 0.00040258 | 0.00020404 |
| KRAS.LUNG.BREAST_UP.V1_UP | 136 | 0.42374364 | 1.71340009 | 0.00051781 | 0.00026244 |
| P53_DN.V1_DN | 193 | 0.38572892 | 1.61889018 | 0.00075997 | 0.00038517 |
| SIRNA_EIF4GI_UP | 92 | 0.48369093 | 1.84829077 | 0.00078633 | 0.00039853 |
| RAF_UP.V1_UP | 192 | 0.38157478 | 1.59190072 | 0.00142189 | 0.00072064 |
| ERBB2_UP.V1_UP | 190 | 0.37681972 | 1.57813237 | 0.00149945 | 0.00075995 |
| EGFR_UP.V1_UP | 191 | 0.36249147 | 1.51974573 | 0.00222858 | 0.00112949 |
| LTE2_UP.V1_UP | 188 | 0.36352186 | 1.51365056 | 0.00222858 | 0.00112949 |
| KRAS.600_UP.V1_DN | 271 | -0.4724353 | -1.5061418 | 0.00222858 | 0.00112949 |
| STK33_DN | 277 | 0.33067498 | 1.42285335 | 0.00222858 | 0.00112949 |
| CYCLIN_D1_KE_.V1_DN | 187 | -0.5066232 | -1.5822447 | 0.00238088 | 0.00120669 |
| AKT_UP_MTOR_DN.V1_UP | 177 | 0.37001476 | 1.53513426 | 0.00250234 | 0.00126824 |
| EIF4E_DN | 99 | 0.43843667 | 1.6851388 | 0.00316258 | 0.00160287 |
| KRAS.PROSTATE_UP.V1_DN | 137 | -0.5262641 | -1.6012712 | 0.00316258 | 0.00160287 |
| KRAS.KIDNEY_UP.V1_DN | 130 | -0.5248992 | -1.5876315 | 0.00316258 | 0.00160287 |
| MEK_UP.V1_DN | 191 | 0.3542014 | 1.48498957 | 0.00357518 | 0.00181198 |
| STK33_NOMO_DN | 275 | 0.32024737 | 1.37184053 | 0.00467779 | 0.00237081 |
| RB_DN.V1_DN | 122 | 0.4030223 | 1.62376767 | 0.00518035 | 0.00262552 |
| PTEN_DN.V2_UP | 135 | 0.38691388 | 1.56430857 | 0.00544271 | 0.00275849 |
| MYC_UP.V1_UP | 177 | 0.35235432 | 1.46186383 | 0.00575731 | 0.00291793 |
| KRAS.600.LUNG.BREAST_UP.V1_UP | 270 | 0.31669428 | 1.38333226 | 0.00575731 | 0.00291793 |
| SNF5_DN.V1_DN | 154 | 0.37018432 | 1.53025186 | 0.00641372 | 0.00325062 |
| CSR_LATE_UP.V1_DN | 146 | 0.37774544 | 1.56271658 | 0.00690945 | 0.00350186 |
| MEK_UP.V1_UP | 195 | 0.34312969 | 1.44114651 | 0.00876804 | 0.00444384 |
| CTIP_DN.V1_DN | 131 | -0.5103057 | -1.5462426 | 0.00938572 | 0.00475689 |
| ATF2_S_UP.V1_UP | 190 | 0.33807488 | 1.4158678 | 0.01231474 | 0.00624139 |
| KRAS.LUNG.BREAST_UP.V1_DN | 136 | -0.4861551 | -1.4775541 | 0.01596239 | 0.0080901 |
| KRAS.LUNG_UP.V1_UP | 133 | 0.37035096 | 1.48577667 | 0.01632227 | 0.0082725 |
| RB_P130_DN.V1_DN | 134 | 0.36627441 | 1.48267779 | 0.01681188 | 0.00852064 |
| PTEN_DN.V1_UP | 179 | -0.4666712 | -1.4517145 | 0.01712931 | 0.00868152 |
| P53_DN.V1_UP | 191 | 0.32541135 | 1.36428727 | 0.01712931 | 0.00868152 |
| P53_DN.V2_DN | 144 | -0.4818378 | -1.470764 | 0.01721956 | 0.00872726 |
| KRAS.BREAST_UP.V1_DN | 140 | -0.4904674 | -1.4918082 | 0.01722365 | 0.00872934 |
| KRAS.LUNG_UP.V1_DN | 138 | -0.4862701 | -1.4779895 | 0.01722365 | 0.00872934 |
| PDGF_UP.V1_UP | 144 | 0.34290397 | 1.40717855 | 0.01722365 | 0.00872934 |
| PRC1_BMI_UP.V1_DN | 180 | -0.4667875 | -1.4527574 | 0.01903934 | 0.00964957 |
| EIF4E_UP | 100 | 0.38238329 | 1.47145303 | 0.01940231 | 0.00983353 |
| RPS14_DN.V1_DN | 185 | 0.32475093 | 1.35732531 | 0.01940231 | 0.00983353 |
| SIRNA_EIF4GI_DN | 96 | 0.38333509 | 1.48533479 | 0.0200196 | 0.01014639 |
| TBK1.DF_DN | 283 | 0.29358654 | 1.27498171 | 0.02398189 | 0.01215456 |
| E2F3_UP.V1_UP | 187 | -0.4572665 | -1.4280979 | 0.02625956 | 0.01330894 |
| LEF1_UP.V1_UP | 192 | 0.31741881 | 1.32424695 | 0.02625956 | 0.01330894 |
| KRAS.AMP.LUNG_UP.V1_UP | 134 | -0.4662687 | -1.4156422 | 0.02883211 | 0.01461276 |
| NOTCH_DN.V1_UP | 179 | -0.4528759 | -1.4088003 | 0.02883211 | 0.01461276 |
| ATF2_S_UP.V1_DN | 183 | -0.446338 | -1.3913087 | 0.02883211 | 0.01461276 |
| KRAS.300_UP.V1_UP | 138 | 0.33841884 | 1.39057511 | 0.02883211 | 0.01461276 |
| RPS14_DN.V1_UP | 189 | 0.31494052 | 1.31820676 | 0.02883211 | 0.01461276 |
| ERBB2_UP.V1_DN | 197 | 0.31026053 | 1.29320505 | 0.02920462 | 0.01480156 |
| CSR_LATE_UP.V1_UP | 164 | 0.32019403 | 1.32029449 | 0.03129088 | 0.01585893 |
| KRAS.300_UP.V1_DN | 134 | -0.4614742 | -1.4010855 | 0.0318821 | 0.01615857 |
| RB_P107_DN.V1_UP | 130 | -0.4635045 | -1.4019347 | 0.03283017 | 0.01663907 |
| BCAT_BILD_ET_AL_DN | 46 | 0.45403149 | 1.50813667 | 0.03530264 | 0.01789218 |
| DCA_UP.V1_UP | 180 | -0.4458282 | -1.387527 | 0.03965407 | 0.02009758 |
| ALK_DN.V1_UP | 135 | 0.33399205 | 1.35034344 | 0.04057011 | 0.02056185 |
| PKCA_DN.V1_UP | 156 | -0.4539461 | -1.3974191 | 0.04164544 | 0.02110685 |
| IL21_UP.V1_DN | 180 | -0.4422233 | -1.3763074 | 0.04259468 | 0.02158795 |
| HINATA_NFKB_MATRIX | 11 | 0.70263646 | 1.64767807 | 0.04282204 | 0.02170318 |
| RAF_UP.V1_DN | 192 | 0.30554325 | 1.27470301 | 0.04333274 | 0.02196201 |
| ATF2_UP.V1_DN | 181 | -0.4379311 | -1.3629073 | 0.04334126 | 0.02196633 |
| KRAS.50_UP.V1_DN | 44 | -0.5555258 | -1.4651564 | 0.04530801 | 0.02296312 |
| PRC2_SUZ12_UP.V1_DN | 179 | -0.4364153 | -1.3575947 | 0.04530801 | 0.02296312 |
| IL21_UP.V1_UP | 181 | -0.4356321 | -1.3557524 | 0.0455453 | 0.02308339 |
| JNK_DN.V1_UP | 179 | -0.4361389 | -1.3567349 | 0.04568175 | 0.02315254 |
| KRAS.50_UP.V1_UP | 47 | 0.44144535 | 1.47111674 | 0.04575401 | 0.02318917 |

**Table S9**. siRNAs for *SATB2* and *circ3915* knockdown experiments. ON-TARGETplus siRNAs are from Dharmacon Reagents.

| ON-TARGETplus siRNA | Catalog # /Sequence |
| --- | --- |
| Non-targeting siRNA (Control) | #D-001810-01 |
| SATB2-1 | #J-023161-09 |
| SATB2-2 | #J-023161-11 |
| SATB2-3 | #J-023161-12 |
| SATB2-Circular+Linear | #J-023161-10 |
| circ3915-1 | [UAAAGGUUUGAUGAUUCCUGUdTdT](http://blast.ncbi.nlm.nih.gov/Blast.cgi?PROGRAM=blastn&PAGE_TYPE=BlastSearch&LINK_LOC=blasthome&QUERY=%3Ehsa_circ_0003915-siRNA1%0ATAAAGGTTTGATGATTCCTGT&DATABASE=nr&EQ_MENU=Homo%C2%A0sapiens%C2%A0(taxid:9606)) |
| circ3915-2 | [GAUUAAAGGUUUGAUGAUUCC](http://blast.ncbi.nlm.nih.gov/Blast.cgi?PROGRAM=blastn&PAGE_TYPE=BlastSearch&LINK_LOC=blasthome&QUERY=%3Ehsa_circ_0003915-siRNA2%0AGATTAAAGGTTTGATGATTCC&DATABASE=nr&EQ_MENU=Homo%C2%A0sapiens%C2%A0(taxid:9606))dTdT |
| circ3915-3 | [UUAAAGGUUUGAUGAUUCCUG](http://blast.ncbi.nlm.nih.gov/Blast.cgi?PROGRAM=blastn&PAGE_TYPE=BlastSearch&LINK_LOC=blasthome&QUERY=%3Ehsa_circ_0003915-siRNA3%0ATTAAAGGTTTGATGATTCCTG&DATABASE=nr&EQ_MENU=Homo%C2%A0sapiens%C2%A0(taxid:9606))dTdT |

**Table S10**. shRNA sequences for stable *SATB2* mRNA and *circ3915* knockdown experiments.

| shRNA Clones/Target | Catalog # /Sequence |
| --- | --- |
| shScrambled Control | GCGCGAUAGCGCUAAUAAUUU |
| shSATB2-1 | GCCAUGCAGAAUUUCCUCAAU |
| shSATB2-2 (Circular+Linear) | GCCUUAAAGGAACUGCUCAAA |
| shcirc3915-1 | [UAAAGGUUUGAUGAUUCCUGU](http://blast.ncbi.nlm.nih.gov/Blast.cgi?PROGRAM=blastn&PAGE_TYPE=BlastSearch&LINK_LOC=blasthome&QUERY=%3Ehsa_circ_0003915-siRNA1%0ATAAAGGTTTGATGATTCCTGT&DATABASE=nr&EQ_MENU=Homo%C2%A0sapiens%C2%A0(taxid:9606)) |
| shcirc3915-2 | [GAUUAAAGGUUUGAUGAUUCC](http://blast.ncbi.nlm.nih.gov/Blast.cgi?PROGRAM=blastn&PAGE_TYPE=BlastSearch&LINK_LOC=blasthome&QUERY=%3Ehsa_circ_0003915-siRNA2%0AGATTAAAGGTTTGATGATTCC&DATABASE=nr&EQ_MENU=Homo%C2%A0sapiens%C2%A0(taxid:9606)) |
| shcirc3915-3 | [UUAAAGGUUUGAUGAUUCCUG](http://blast.ncbi.nlm.nih.gov/Blast.cgi?PROGRAM=blastn&PAGE_TYPE=BlastSearch&LINK_LOC=blasthome&QUERY=%3Ehsa_circ_0003915-siRNA3%0ATTAAAGGTTTGATGATTCCTG&DATABASE=nr&EQ_MENU=Homo%C2%A0sapiens%C2%A0(taxid:9606)) |

**Table S11**. RT-qPCR primers and probes. F= forward primer; R= reverse primer.

| Target | Targeted exon(s) |  | Sequence (5’ 🡪 3’) | T_m_  (°C) | Amplicon length (bp) |
| --- | --- | --- | --- | --- | --- |
| Circular + linear SATB2 | 3 | F | ttggacggctctcttgaatatg | 60 | 130 |
|  |  | R | ccgcagagctgtgagaatac |  |  |
| *SATB2* linear mRNA | 11 | F | tctccccaaacacaccatca | 60 | 125 |
|  |  | R | gcagctcctcgtccttatattca |  |  |
| circ3915 | 3-6 junction | F | gaagattaaaggtttgatgattcctgtc | 60 | 211 |
|  |  | R | ccaccttcccagcttgattatt |  |  |
| GAPDH |  | F | gtcagccgcatcttcttttg | 60 | 100 |
|  |  | R | gcgcccaatacgaccaaatc |  |  |
| RPII |  | F | gcaccacgtccaatgacat | 60 | 267 |
|  |  | R | gtgcggctgcttccataa |  |  |
| TBP |  | F | ttcggagagttctgggattgta | 60 | 227 |
|  |  | R | tggactgttcttcacttcttggc |  |  |

**Table S12**. Nanostring codesets for verifying differentially expressed circular and linear RNA transcripts.

| **Target ID** | **Target Sequence** | **Position** | **Capture Probe T_m_ (°C)** | **Reporter Probe T_m_ (°C)** | **Isoform Coverage and Design Notes** |
| --- | --- | --- | --- | --- | --- |
| ACTB | TGCAGAAGGAGATCACTGCCCTGGCACCCAGCACAATGAAGATCAAGATCATTGCTCCTCCTGAGCGCAAGTACTCCGTGTGGATCGGCGGCTCCATCCT | 1011-1110 | 87 | 87 | NM_001101.5  Also targets alpha and gamma-actin transcripts and a beta-actin pseudogene @ >91% |
| ADAM17 | AGAACGCAGCAATAAAGTTTGTGGGAACTCGAGGGTGGATGAAGGAGAAGAGTGTGATCCTGGCATCATGTATCTGAACAACGACACCTGCTGCAACAGC | 1596-1695 | 80 | 82 | NM_003183.6;XM_017004785.2;XM_024453056.1;NM_001382777.1;NM_001382778.1;XM_011510375.3 |
| ALDH1A1 | ATTGCTGAGCCAGTCACCTGTGTTCCAGGAGCCGAATCAGAAATGTCATCCTCAGGCACGCCAGACTTACCTGTCCTACTCACCGATTTGAAGATTCAAT | 12-111 | 84 | 82 | NM_000689.5 |
| ALDH1A3 | TTCGGCTTCAAACCAATACTGCCTTTGGAATATGACAGAATCAATAGCCCAGAGAGCTTAGTCAAAGACGATATCACGGTCTACCTTAACCAAGGCACTT | 2281-2380 | 80 | 83 | NM_001293815.2;NM_000693.4 |
| AS3MT | GAACTGCCAGAAGAAATCAGGACACACAAAGTTTTATGGGGTGAGTGTCTGGGTGGTGCTTTATACTGGAAGGAACTTGCTGTCCTTGCTCAAAAAATTG | 711-810 | 82 | 81 | NM_020682.4  Also targets BORCS7-ASMT readthrough (NR_037644) @ 100% |
| BAMBI | CCAAAAAGGGGCAGGTTGCAAAGTTAGACTTGGAATGCATGGTGCCGGTCAGTGGGCACGAGAACTGCTGTCTGACCTGTGATAAAATGAGACAAGCAGA | 1011-1110 | 79 | 81 | NM_012342.3 |
| CASP7 | ATCAATGACACAGATGCTAATCCTCGATACAAGATCCCAGTGGAAGCTGACTTCCTCTTCGCCTATTCCACGGTTCCAGGCTATTACTCGTGGAGGAGCC | 916-1015 | 80 | 82 | NM_033339.5;NM_001267057.1;NM_001320911.2;NM_033338.6;NM_001267056.2;XM_011540260.1;XM_006718017.3;XM_017016763.1;NM_001227.5;NM_001267058.2;XM_017016764.1;NM_033340.4 |
| CAV1 | AACCGCGACCCTAAACACCTCAACGATGACGTGGTCAAGATTGACTTTGAAGATGTGATTGCAGAACCAGAAGGGACACACAGTTTTGACGGCATTTGGA | 435-534 | 84 | 83 | NM_001172896.2;NM_001172895.1;NM_001753.5;NM_001172897.2 |
| CD44 | ACACCATGGACAAGTTTTGGTGGCACGCAGCCTGGGGACTCTGCCTCGTGCCGCTGAGCCTGGCGCAGATCGATTTGAATATAACCTGCCGCTTTGCAGG | 430-529 | 85 | 85 | NM_001001389.2;NM_001001390.2;XM_011520485.2;XM_011520484.2;NM_001001392.2;NM_001001391.2;XM_006718390.4;NM_001202555.2;XM_005253238.3;XM_005253239.3;NM_001202556.2;XM_005253240.3;XM_011520486.2;XM_005253232.3;XM_011520488.2;XM_005253235.3;XM_005253231.3;NM_001202557.2;XM_017018585.2;XM_011520482.2;XM_017018583.2;XM_006718388.2;XM_017018584.2;XM_011520489.3;XM_011520483.2;NM_000610.4;XM_011520487.3 |
| CDH1 | GAATTGCTCACATTTCCCAACTCCTCTCCTGGCCTCAGAAGACAGAAGAGAGACTGGGTTATTCCTCCCATCAGCTGCCCAGAAAATGAAAAAGGCCCAT | 536-635 | 80 | 78 | NM_001317185.2;NM_001317186.2;NM_004360.5;NM_001317184.2 |
| CDH2 | GGTCATCCCTCCAATCAACTTGCCAGAAAACTCCAGGGGACCTTTTCCTCAAGAGCTTGTCAGGATCAGGTCTGATAGAGATAAAAACCTTTCACTGCGG | 942-1041 | 84 | 82 | NM_001308176.2;XM_017025514.2;XM_011525788.1;NM_001792.5 |
| CDH3 | CCCTCGACCGTGAGGATGAGCAGTTTGTGAGGAACAACATCTATGAAGTCATGGTCTTGGCCATGGACAATGGAAGCCCTCCCACCACTGGCACGGGAAC | 2006-2105 | 81 | 83 | NM_001793.6;NM_001317195.3;NM_001317196.2;XM_011522800.3 |
| CDK8 | GGTATGGGCCATGAGAATGTACTGTACAACCACATCTTCAAAATGTCCAGTAGCCAAGTTCCACCACTTTTCACAGATTGGGGTAGTGGCTTCCAAGTTG | 1495-1594 | 83 | 80 | NM_001346501.2;XM_011534865.2;XM_017020318.1;NM_001318368.2;XM_017020317.2;NM_001260.3 |
| CDKN1A | CATGTGTCCTGGTTCCCGTTTCTCCACCTAGACTGTAAACCTCTCGAGGGCAGGGACCACACCCTGTACTGTTCTGTGTCTTTCACAGCTCCTCCCACAA | 1976-2075 | 80 | 82 | NM_000389.5;NM_001374513.1;NM_001220777.2;NM_001374510.1;NM_001374511.1;NM_001374509.1;NM_078467.3;NM_001374512.1;NM_001220778.2;NM_001291549.3  Sub-maximal isoform coverage due to non-overlapping transcripts |
| CER1 | CAGAACAACCTTTGCTTTGGGAAATGCGGGTCTGTTCATTTTCCTGGAGCCGCGCAGCACTCCCATACCTCCTGCTCTCACTGTTTGCCTGCCAAGTTCA | 589-688 | 79 | 84 | NM_005454.3;XR_001746419.1 |
| CLDN1 | GCAAAGTCTTTGACTCCTTGCTGAATCTGAGCAGCACATTGCAAGCAACCCGTGCCTTGATGGTGGTTGGCATCCTCCTGGGAGTGATAGCAATCTTTGT | 411-510 | 83 | 82 | NM_021101.5 |
| CLDN3 | TGCTCACCCTCGTGCCGGTGTCCTGGTCGGCCAACACCATTATCCGGGACTTCTACAACCCCGTGGTGCCCGAGGCGCAGAAGCGCGAGATGGGCGCGGG | 607-706 | 82 | 83 | NM_001306.4 |
| CLTC | GGGTATCAACCCAGCAAACATTGGCTTCAGTACCCTGACTATGGAGTCTGACAAATTCATCTGCATTAGAGAAAAAGTAGGAGAGCAGGCCCAGGTGGTA | 291-390 | 82 | 78 | NM_004859.4;NM_001288653.2 |
| COL1A1 | CAGAAACATCGGATTTGGGGAACGCGTGTCAATCCCTTGTGCCGCAGGGCTGGGCGGGAGAGACTGTTCTGTTCCTTGTGTAACTGTGTTGCTGAAAGAC | 5211-5310 | 82 | 81 | XM_005257059.4;XM_005257058.4;XM_011524341.1;NM_000088.4 |
| COL4A2 | GGCATTTCCTTGAAGGGAGAAGAAGGAATCATGGGCTTTCCTGGACTGAGGGGTTACCCTGGCTTGAGTGGTGAAAAAGGATCACCAGGACAGAAGGGAA | 1151-1250 | 81 | 82 | NM_001846.4 |
| CREBBP | GTGTCAGAGACGAGAGCAAGCAAACGGAGAGGTTCGGGCCTGCTCGCTCCCGCATTGTCGAACCATGAAAAACGTTTTGAATCACATGACGCATTGTCAG | 1302-1401 | 84 | 81 | NM_001079846.1;XM_005255125.4;XM_017022944.1;XM_006720848.3;XM_011522381.2;XM_011522382.3;XM_005255124.4;NM_004380.3 |
| CTCF | ATTGGTTCGGCATCGTCGTTACAAACACACCCACGAGAAGCCATTCAAGTGTTCCATGTGCGATTACGCCAGTGTAGAAGTCAGCAAATTAAAACGTCAC | 491-590 | 80 | 78 | XM_017022868.1;NM_001363916.1;XM_005255775.4;NM_006565.4;NM_001191022.2 |
| CTNNB1 | TCTTGCCCTTTGTCCCGCAAATCATGCACCTTTGCGTGAGCAGGGTGCCATTCCACGACTAGTTCAGTTGCTTGTTCGTGCACATCAGGATACCCAGCGC | 1816-1915 | 82 | 81 | XM_006712985.1;XM_024453358.1;XM_024453356.1;XM_024453357.1;XM_024453360.1;NM_001098209.2;XM_006712983.2;XM_017005738.1;NM_001098210.2;NM_001330729.2;XM_024453359.1;NM_001904.4 |
| CUL1 | CTGGTAATGTCTGCATTCAACAATGACGCTGGCTTTGTGGCTGCTCTTGATAAGGCTTGTGGTCGCTTCATAAACAACAACGCGGTTACCAAGATGGCCC | 1488-1587 | 81 | 84 | NM_001370660.1;NM_001370661.1;NM_001370664.1;NM_001370662.1;NM_003592.3;NM_001370663.1 |
| DKK1 | CGGCACGGTTTCGTGGGGACCCAGGCTTGCAAAGTGACGGTCATTTTCTCTTTCTTTCTCCCTCTTGAGTCCTTCTGAGATGATGGCTCTGGGCGCAGCG | 76-175 | 81 | 81 | NM_012242.4 |
| DNMT1 | CAAAACCAATCTATGATGATGACCCATCTCTTGAAGGTGGTGTTAATGGCAAAAATCTTGGCCCCATAAATGAATGGTGGATCACTGGCTTTGATGGAGG | 1496-1595 | 81 | 83 | NM_001130823.3;NM_001318730.2;NM_001318731.2;NM_001379.4 |
| DNMT3A | GGCGAGAGGACTGGCCCTCCCGGCTCCAGATGTTCTTCGCTAATAACCACGACCAGGAATTTGACCCTCCAAAGGTTTACCCACCTGTCCCAGCTGAGAA | 2057-2156 | 83 | 83 | XM_017003527.1;NM_001320893.1;XR_001738657.1;XM_017003526.1;NR_135490.2;XM_011532664.2;NM_001375819.1;NM_022552.5;XM_011532667.3;XM_011532662.2;XM_011532666.2;NM_175629.2;XM_011532663.2;NM_153759.3;XM_005264175.5  Sub-maximal isoform coverage due to non-overlapping transcript variants |
| DNMT3B | CGAAGGCGGCCCATTCGAGTCCTGTCATTGTTTGATGGCATCGCGACAGGCTACCTAGTCCTCAAAGAGTTGGGCATAAAGGTAGGAAAGTACGTCGCTT | 1951-2050 | 83 | 82 | NM_001207055.2;NM_175848.2;NM_001207056.2;XR_936510.2;NM_175850.3;NM_175849.2;XM_011528653.2;XM_011528654.2;XR_936511.2;NM_006892.4;XR_936512.2 |
| EFNA1 | TGCTGCCCCACGCCTCTTCCCACTTGCCTGGACTGTGCTGCTCCTTCCACTTCTGCTGCTGCAAACCCCGTGAAGGTGTATGCCACACCTGGCCTTAAAG | 651-750 | 82 | 82 | NM_004428.3;NM_182685.2 |
| EP300 | CCAGCCAGGCCCAACAGAGCAGTCCTGGATTAGGTTTGATAAATAGCATGGTCAAAAGCCCAATGACACAGGCAGGCTTGACTTCTCCCAACATGGGGAT | 716-815 | 82 | 82 | NM_001362843.2;NM_001429.4 |
| EZH1 | CAACTTAGGCAGTTCCCAACAAGCGCTAGCCTGTAATTGTAGCTTTCCACATCAAGAGTCCTTATGTTATTGGGATGCAGGCAAACCTCTGTGGTCCTAA | 2721-2820 | 81 | 81 | XM_017024352.2;NM_001991.5;NM_001321081.2;NM_001321079.2;XM_011524517.2;XM_017024350.1;XM_017024351.1;NM_001321082.2;XM_005257145.2 |
| EZH2 | ACACAGAAACAGCTCTAGACAACAAACCTTGTGGACCACAGTGTTACCAGCATTTGGAGGGAGCAAAGGAGTTTGCTGCTGCTCTCACCGCTGAGCGGAT | 1122-1221 | 85 | 85 | XM_024446680.1;XM_011515889.2;XM_005249963.4;NM_004456.5;XM_017011820.2;XM_017011818.1;XM_005249964.4;XM_017011821.1;XM_011515901.3;XM_011515887.3;XM_011515898.2;XM_011515890.2;XM_011515883.2;XM_011515884.2;NM_001203247.2;NM_001203249.2;NM_152998.3;XM_017011819.1;XR_002956414.1;XM_011515894.2;XM_011515888.2;XR_001744581.1;XM_011515891.3;XM_011515896.2;XM_011515895.2;XM_011515885.2;XM_011515893.2;XR_002956413.1;XM_005249962.4;XM_011515892.2;NM_001203248.2;XM_011515897.2;XM_011515886.2;XM_017011817.2;XM_011515899.3 |
| FGF2 | GTCCGGGAGAAGAGCGACCCTCACATCAAGCTACAACTTCAAGCAGAAGAGAGAGGAGTTGTGTCTATCAAAGGAGTGTGTGCTAACCGTTACCTGGCTA | 621-720 | 82 | 80 | NM_002006.5;NM_001361665.2 |
| FGFR2 | AAAGATGATGCCACAGAGAAAGACCTTTCTGATCTGGTGTCAGAGATGGAGATGATGAAGATGATTGGGAAACACAAGAATATCATAAATCTTCTTGGAG | 2205-2304 | 85 | 78 | NM_022970.3;NM_000141.5;XM_017015925.2;XM_024447887.1;XM_017015920.2;NM_001144915.2;NM_001144913.1;NM_001144917.2;NM_001144916.2;XM_017015921.2;NM_001144919.2;XM_006717710.4;XM_024447892.1;XM_017015924.2;XM_024447891.1;XM_006717708.3;XM_024447890.1;NM_001320658.2;NM_001144914.1;NM_023029.2;XM_024447888.1;NM_001144918.2;NR_073009.2;NM_001320654.2;XM_024447889.1 |
| FN1 | GGGAATGGACATGCATTGCCTACTCGCAGCTTCGAGATCAGTGCATTGTTGATGACATCACTTACAATGTGAACGACACATTCCACAAGCGTCATGAAGA | 1777-1876 | 83 | 81 | NM_001306129.2;XM_005246416.1;XM_005246404.1;NM_001365524.2;NM_001365519.2;NM_212482.4;NM_001365517.2;XM_017003692.1;NM_001306130.2;NM_001306132.2;NM_212474.3;NM_212476.3;NM_001365522.2;NM_001365521.2;XM_005246407.1;XM_005246402.1;NM_054034.3;XM_005246398.1;XM_005246411.1;XM_024452769.1;XM_005246397.1;XM_005246401.1;NM_001306131.2;XM_017003695.1;NM_212478.3;XM_005246410.1;NM_001365518.2;XM_005246403.1;XM_005246399.1;XM_005246408.1;NM_001365523.2;NM_002026.4;NM_001365520.2;XM_024452770.1 |
| FOXA1 | TGATACATTCTCAAGAGTTGCTTGACCGAAAGTTACAAGGACCCCAACCCCTTTGTCCTCTCTACCCACAGATGGCCCTGGGAATCAATTCCTCAGGAAT | 2466-2565 | 83 | 84 | NM_004496.5;XM_017021246.1 |
| FOXC2 | CCCCGGCCACACGTTCGCGGCCCAGCAGCAAACTTTCCCCAACGTGCGGGAGATGTTCAACTCCCACCGGCTGGGGATTGAGAACTCGACCCTCGGGGAG | 1390-1489 | 85 | 86 | NM_005251.3 |
| FOXD3 | CCGCGCGCCCGCCGGGGATAGCTTTCCATACAGGTAAAACCGAAAACCGAATTTTCCAAAAATGCACCCCGACGGCGCCTGCTCTTAGTACCGTGGGGAT | 1650-1749 | 81 | 80 | NM_012183.3 |
| FZD7 | CCCTCTACTGAGAAGTGACCTGGAAGTGAGAAGTTCTTTGCAGATTTGGGGCGAGGGGTGATTTGGAAAAGAAGACCTGGGTGGAAAGCGGTTTGGATGA | 1891-1990 | 79 | 81 | NM_003507.2 |
| GAPDH | GAACGGGAAGCTTGTCATCAATGGAAATCCCATCACCATCTTCCAGGAGCGAGATCCCTCCAAAATCAAGTGGGGCGATGCTGGCGCTGAGTACGTCGTG | 387-486 | 85 | 86 | NM_001256799.3;NM_001357943.2;NM_002046.7;NM_001289746.2;NM_001289745.3;NR_152150.2 |
| HIST1H2BJ | GCACGCCGTGTCCGAGGGTACTAAGGCCGTCACCAAGTACACCAGCGCTAAGTAAACAGTGAGTTGGTTGCAAACTCTCAACCCTAACGGCTCTTTTAAG | 373-472 | 86 | 79 | NM_021058.4 |
| HIST1H2BK | TACTTCCCGTTTTCTCGATCTGCTGCTCGTCTCAGGCTCGTAGTTCGCCTTCAACATGCCGGAACCAGCGAAGTCCGCTCCCGCGCCCAAGAAGGGCTCG | 6-105 | 78 | 93 | NM_080593.2;NM_001312653.2 |
| HIST1H2BN | GGTGTCGGAGGGCACCAAGGCCGTCACCAAGTACACCAGTTCCAAGTGAGCCCGCCCACCGCGGAACGTTCGGTCAGTCTCGGCCCACACCCCAAAGGCT | 333-432 | 85 | 85 | NM_003520.4 |
| HIST2H2BE | TGCGAGGCACTTACCATGTAGATACGGGCTCAAAAGTCACCTCTCAGAGACCTACGTCATCCACTCAGGAATTCGCGCCTCTCATACTTGCCTGTCTCAT | 1181-1280 | 84 | 80 | NM_003528.3 |
| HIST1H2BC | GCTGGCCAAGCACGCCGTGTCGGAGGGCACCAAGGCCGTCACCAAGTACACCAGCTCCAAGTAAACATTCCAAGTAAGCGTCTTAACACCTAACCCCAAA | 318-417 | 86 | 81 | NM_003526.3  Also targets HIST1H2BD (NM_003526) @ 92% |
| HIST1H2BD | CTCAGGTGTTTGCAACAGTGTTCTAACTATTAACGCTACGATGCCTGAACCTACCAAGTCTGCTCCTGCCCCAAAGAAGGGCTCCAAGAAGGCGGTGACT | 10-109 | 70 | 87 | XM_005249039.4;NM_021063.4;NM_138720.2 |
| HIST1H2BF | GAGAGCTATTCCGTGTACGTGTACAAGGTGCTAAAGCAGGTCCACCCCGACACCGGCATCTCATCCAAGGCCATGGGCATCATGAACTCCTTCGTCAACG | 106-205 | 87 | 85 | NM_003522.4 |
| HIST1H2BH | CCTGGCTCATTACAACAAGCGTTCGACCATCACCTCCAGGGAGATCCAGACAGCCGTGCGCCTGCTGCTGCCTGGGGAACTGGCCAAGCACGCCGTGTCC | 240-339 | 87 | 95 | NM_003524.3  Also targets multiple other HIST1 memebers HIST1H2BO;HIST1H2BD;HIST1H2BK;HIST1H2BE @93%+ |
| HAND1 | TTTGGAGCGAATTTAGAACCTCAGCCCTATCTCCATTTCCCTATCTGGCTCTTTCTCTCTTGTCCCTCCATATGATCCGCCCCGACGCCGTCTTCTCTAA | 1231-1330 | 79 | 78 | NM_004821.3;XM_005268531.1 |
| HAND2 | TCCATATTTTAATACGAAGAGGACACTCCCGTGTGGTAAGGGATCCCGTCGTCTCATAGATTCTGTGTGCGTGAATGTTCCCTCTTGGCTGTGTAGACAC | 1822-1921 | 81 | 80 | NM_021973.3 |
| HDAC1 | CAAGCCGGTCATGTCCAAAGTAATGGAGATGTTCCAGCCTAGTGCGGTGGTCTTACAGTGTGGCTCAGACTCCCTATCTGGGGATCGGTTAGGTTGCTTC | 786-885 | 81 | 82 | NM_004964.3;XM_011541309.2 |
| HES-1 | GCTGGAGAGGCGGCTAAGGTGTTTGGAGGCTTCCAGGTGGTACCGGCTCCCGATGGCCAGTTTGCTTTCCTCATTCCCAACGGGGCCTTCGCGCACAGCG | 861-960 | 82 | 82 | NM_005524.4 |
| HIST2H2BF | AGTTTTGCATGGGGGCAAGAGACTATGGGGAGTTTCTAGATGTGGAAGATCCTGAGTGTAAGGGTGGGGCGAGTCCCTTAAAAATTATAAACCCGTTTCC | 129-228 | 82 | 81 |  |
| ICAM1 | AAATACTGAAACTTGCTGCCTATTGGGTATGCTGAGGCCCCACAGACTTACAGAAGAAGTGGCCCTCCATAGACATGTGTAGCATCAAAACACAAAGGCC | 2254-2353 | 86 | 84 | NM_000201.3 |
| IGFBP3 | TATCAAAATATTCAGAGACTCGAGCACAGCACCCAGACTTCATGCGCCCGTGGAATGCTCACCACATGTTGGTCGAAGCGGCCGACCACTGACTTTGTGA | 1256-1355 | 81 | 82 | NM_000598.5;NM_001013398.2 |
| ITGA5 | GCCAGCTGCACTGATGCTGCCCCTCATCTCTCTGCCCAACCCTTCCCTCACCTTGGCACCAGACACCCAGGACTTATTTAAACTCTGTTGCAAGTGCAAT | 4114-4213 | 85 | 82 | XM_024448970.1;NM_002205.5 |
| ITGA6 | CTCATGCGAGCCTTCATTGATGTGACTGCTGCTGCCGAAAATATCAGGCTGCCAAATGCAGGCACTCAGGTTCGAGTGACTGTGTTTCCCTCAAAGACTG | 3066-3165 | 81 | 80 | XM_017004007.1;XM_017004008.1;NM_000210.4;NM_001079818.3;NM_001365530.2;NM_001365529.2;NM_001316306.2;XM_017004005.1;XM_017004006.1 |
| ITGB1 | TTTTAACATTACCAAGGTAGAAAGTCGGGACAAATTACCCCAGCCGGTCCAACCTGATCCTGTGTCCCATTGTAAGGAGAAGGATGTTGACGACTGTTGG | 2001-2100 | 81 | 83 | NM_133376.3;NM_033668.2;NM_002211.4  Target accesion has been removed from RefSeq but probe hits current RefSeq transcripts (NM_002211.3;NM_133376.2&NM_033668.2) @100% |
| ITGB3 | GAATAAGCCTTGGAATTAGATATGGGGCAATGACTGAGCCCTGTCTCACCCATGGATTACTCCTTACTGTAGGGAATGGCAGTATGGTAGAGGGATAAAT | 4486-4585 | 82 | 79 | NM_000212.3 |
| KRAS | GCATGGACTGTGTCCCCACGGTCATCCAGTGTTGTCATGCATTGGTTAGTCAAAATGGGGAGGGACTAGGGCAGTTTGGATAGCTCAACAAGATACAATC | 1791-1890 | 81 | 80 | NM_004985.5;NM_033360.4;NM_001369786.1;NM_001369787.1 |
| LAMC1 | TCTTGATAGGAAAGTGTCTGACCTGGAGAATGAAGCCAAGAAGCAGGAGGCTGCCATCATGGACTATAACCGAGATATCGAGGAGATCATGAAGGACATT | 4916-5015 | 82 | 81 | NM_002293.4 |
| MECP2 | CGCAGAAAAGTACAAACACCGAGGGGAGGGAGAGCGCAAAGACATTGTTTCATCCTCCATGCCAAGGCCAAACAGAGAGGAGCCTGTGGACAGCCGGACG | 1440-1539 | 88 | 88 | NM_001110792.2;NM_001369394.2;XM_011531166.2;XM_024452383.1;NM_001369392.2;NM_001369391.2;NM_001369393.2;NM_001386137.1;NM_001386138.1;NM_004992.4;NM_001386139.1;NM_001316337.2 |
| MMP1 | AAATGGGCTTGAAGCTGCTTACGAATTTGCCGACAGAGATGAAGTCCGGTTTTTCAAAGGGAATAAGTACTGGGCTGTTCAGGGACAGAATGTGCTACAC | 1118-1217 | 84 | 84 | NM_002421.4;NM_001145938.2 |
| MMP14 | GACAAGATTGATGCTGCTCTCTTCTGGATGCCCAATGGAAAGACCTACTTCTTCCGTGGAAACAAGTACTACCGTTTCAACGAAGAGCTCAGGGCAGTGG | 1471-1570 | 80 | 79 | NM_004995.4 |
| MMP2 | CCCGGAGGGGCCTGGCAGCCGTGCCTTCAGCTCTACAGCTAATCAGCATTCTCACTCCTACCTGGTAATTTAAGATTCCAGAGAGTGGCTCCTCCCGGTG | 2361-2460 | 82 | 81 | NM_001127891.3;NM_001302510.2;NM_001302509.2;NM_004530.6;NM_001302508.1 |
| CBP | TAAAGCATCACTTAGGAGCTGCTACTCCAGAAAATCCAGAAATAGAGCTGCTTCGCCTAGAACTGGCCGAAATGAAAGAGAAGTATGAAGCTATTGTAGA | 216-315 | 83 | 82 | NR_037632.1;NM_012333.5 |
| NAP1L1 | CAAAGACATACAGGATGAGGTCAGAACCAGATGATTCTGATCCCTTTTCTTTTGATGGACCAGAAATTATGGGTTGTACAGGGTGCCAGATAGATTGGAA | 1101-1200 | 80 | 81 | NM_004537.7;NM_001330232.2;XR_001748714.2;NM_001307924.3;XR_001748717.2;XM_017019339.1;XR_001748715.2;XM_017019340.2;XM_017019338.1;XM_011538393.2;XR_002957328.1;NM_001330231.2;NM_139207.5;XM_024448983.1;XR_001748716.1 |
| NCAM2 | ATATTAAAGATGTGAAGTTGTCAGATTCAGGGAGATATGACTGTGAAGCTGCAAGCAGAATTGGAGGGCATCAAAAGAGCATGTACCTTGATATTGAATA | 1365-1464 | 78 | 82 | XM_011529582.3;NM_001352592.2;NM_004540.5;XM_017028356.2;NM_001352596.2;NM_001352597.2;XM_011529576.3;XM_011529580.3;XM_011529585.3;NM_001352595.2;XM_024452081.1;XM_011529581.3;XM_017028357.2;XM_011529575.3;NM_001352594.2;NM_001352593.2;NM_001352591.2 |
| NGFR | TGAAGAAAAGTGGGCCAGTGTGGGAATGCGGCAAGAAGGAATTGACTTCGACTGTGACCTGTGGGGATTTCTCCCAGCTCTAGACAACCCTGCAAAGGAC | 2731-2830 | 81 | 83 | NM_002507.4 |
| NOTCH1 | AGGCAAAGCTGGCTCACCTTCCGCACGCGGATTAATTTGCATCTGAAATAGGAAACAAGTGAAAGCATATGGGTTAGATGTTGCCATGTGTTTTAGATGG | 8212-8311 | 83 | 78 | NM_017617.5;XM_011518717.2 |
| NTRK2 | TCACATGAACAATGGGGACTACACTCTAATAGCCAAGAATGAGTATGGGAAGGATGAGAAACAGATTTCTGCTCACTTCATGGGCTGGCCTGGAATTGAC | 1606-1705 | 81 | 83 | NM_001369535.1;NM_001369539.1;NM_001369534.1;NM_001018065.2;NM_001369532.1;NM_001369552.1;NM_001369544.1;XM_005252001.3;XM_017014753.2;NM_001369540.1;NM_001369551.1;NM_001369542.1;XM_005252004.2;NM_001291937.2;XM_005252003.3;NM_001369548.1;NM_001369536.1;NM_001369547.1;XM_011518720.3;NM_001369543.1;NM_001369546.1;NM_001369541.1;NM_001007097.3;NM_001369545.1;NM_001369538.1;XM_017014752.1;XM_011518718.3;XM_017014751.2;NM_006180.6;NM_001369550.1;XM_017014760.2;XM_017014755.1;XM_005252006.4;NM_001369533.1;NM_001369537.1;NM_001018064.3;NM_001369549.1;NM_001018066.3 |
| PARG | GTGCTGGATCACAATGAATGTCTAATTATCACAGGTACTGAGCAGTACAGTGAATACACAGGCTATGCTGAGACATATCGTTGGTCCCGGAGCCACGAAG | 2581-2680 | 78 | 82 | NR_130168.3;NM_001303489.3;NR_136753.3;NM_001324381.3;NR_136756.3;NM_001303486.3;NR_136755.3;NR_136754.3;NM_003631.5;NR_136752.3;NR_130169.3;XM_011540305.1;NM_001303487.3;XR_001747247.1 |
| PARP1 | AAGGTTTGGGCAAAACTACCCCTGATCCTTCAGCTAACATTAGTCTGGATGGTGTAGACGTTCCTCTTGGGACCGGGATTTCATCTGGTGTGAATGACAC | 3017-3116 | 82 | 83 | NM_001618.4 |
| PARP10 | GATGGAGCCAGGGGCGATGCGCTTCCTGCAGCTCTACCATGAGGACCTTCTTGCGGGCCTGGGAGACGTCGCTCTCTTGCCACTTGAAGGACCGGATATG | 1455-1554 | 83 | 84 | NM_032789.5;NR_134234.2;XM_011517336.3;NM_001317895.2 |
| PARP11 | AGATGCTGCTTATTCCAGTCGTTTCTGCAAAGATGACATAAAGCATGGGAACACATTCCAAATTCATGGTGTCAGCTTGCAACAGCGGCATCTGTTTAGA | 815-914 | 80 | 79 | NM_001286522.2;XM_011520973.2;XM_011520970.2;XM_017019671.1;NM_001286521.2;NM_020367.6;NR_104461.2 |
| PARP12 | ACGGCTCTTGTGCCTTTCAAAAGCAGTGCATCAAGCTCCATATCTGCCAGTATTTTTTACAGGGGGAATGCAAGTTTGGCACTAGCTGTAAGAGATCCCA | 1371-1470 | 83 | 79 | NM_022750.4;XR_001744855.1;XM_011516491.2;NR_130117.2;XM_005250038.3;XM_005250040.4;XR_927514.2;XM_005250039.4 |
| PARP14 | AGAGCCCGAAGAGGTCGGGAGGCGGCGAGTGTGAGGTCCGCCAGGATCCCAGGAGCCCATCCCGCTTCCTGGTGTTCTTCTACCCGGAGGACGTTCGGCA | 154-253 | 91 | 87 | XR_002959544.1;NM_017554.3;XM_011512928.3;XM_011512929.2 |
| PARP15 | GAAGATTGAGAGGATACAGAATGCATTTCTCTGGCAGAGCTACCAGGTAAAGAAAAGGCAAATGGATATCAAGAATGACCATAAGAATAATGAGAGACTC | 899-998 | 85 | 83 | NM_001308321.2;XM_005247160.4;XM_011512475.3;NM_152615.3;XM_011512477.3;XM_005247159.4;XM_011512478.3;XM_017005792.2;XM_017005791.2;NM_001308320.2;XM_011512476.3;NM_001113523.3;XM_011512479.3 |
| PARP16 | ACGCCTGTTCCTGCACCGGACTTCCTGTTTGAAATTGAGTACTTTGACCCAGCCAACGCCAAATTTTATGAGACCAAAGGAGAACGAGACCTAATCTATG | 766-865 | 81 | 83 | XR_429460.3;XM_017022387.2;XM_017022386.2;XM_024449969.1;XM_006720590.3;NM_001316943.2;XR_001751341.2;XR_001751343.2;NM_017851.6;XR_429459.3;XM_006720592.3;XM_006720589.3;XM_011521742.3;XR_001751342.2 |
| PARP2 | GTTATGAGTTCAAAGTGATTTCCCAGTACCTACAATCTACCCATGCTCCCACACACAGCGACTATACCATGACCTTGCTGGATTTGTTTGAAGTGGAGAA | 1155-1254 | 78 | 82 | XM_005267247.3;NM_005484.4;XM_017020912.1;NM_001042618.2 |
| PARP3 | TACAACTGCACCCTGAACCAGACCAACATCGAGAACAACAACAACAAGTTCTACATCATCCAGCTGCTCCAAGACAGCAACCGCTTCTTCACCTGCTGGA | 567-666 | 83 | 83 | NM_001370240.1;NM_005485.6;XM_017005490.1;NM_001370239.1;NM_001003931.4 |
| PARP4 | CATGGTTAATGTCTGTGAAACTAATTTGTCCAAACCCAACCCACCATCCCTGGCCAAATACCGAGCTTTGAGGTGCAAAATTGAGCATGTTGAACAGAAT | 1171-1270 | 79 | 82 | XM_011534932.2;NM_006437.4;XM_011534931.1;XR_941496.1 |
| PARP6 | TGGTGTGTGATGAGCAGCATGTCTTCCAAAATGGATCTATGCTGAAGCCAGCTGTCTGTACTCGTGAACTATGCGTTTTCTCCTTCTACACACTGGGCGT | 1319-1418 | 83 | 80 | NM_001323515.2;NM_001323526.2;NR_136606.2;NM_001323523.2;NR_136610.2;XR_002957660.1;NR_136599.2;XM_024449996.1;NR_136594.2;XR_002957663.1;NM_001323516.2;NR_136604.2;XR_001751361.2;XR_001751363.1;NR_136605.2;XM_011521805.2;NM_001323530.2;NM_020214.4;NR_136609.2;XR_002957662.1;NM_001323531.2;XR_001751362.2;NM_001323524.2;XR_002957661.1;XM_024449995.1;XR_429467.4;XR_002957664.1;NR_136603.2;NR_136611.2;NR_136596.2;NR_136607.2;XM_024449997.1;NR_136608.2;XM_024449998.1;XM_017022418.2;NM_001323522.2;NM_001323532.2;XR_002957665.1;XR_002957659.1;NM_001323521.2;NM_001323519.2;NM_001323528.2;NM_001323525.2 |
| PARP8 | CTATGACAGCAATTAAATCGCACAAACTTTTGAACCGTCCTTGCCCTGCAGCTGTTAAGTCAGAGGAATGCCTAACTCTAAAGTCGCATAGACTATTGAC | 1216-1315 | 78 | 82 | XM_011543640.3;XM_011543636.2;XM_011543631.3;NM_001178056.2;XM_011543633.3;XM_011543634.2;XM_005248596.5;XM_011543641.2;NM_024615.4;NM_001331028.2;XM_011543642.2;XM_017009851.1;XR_002956178.1;XM_011543632.3;XM_011543643.2;XM_024446206.1;XM_011543635.2;NM_001178055.2 |
| PARP9 | CTATTGTAGAGACTATCCGGGTTAGTTTGCAAGGGAAGCCAATGATGAGTAATTTGAAAGAAATTCACCTGGTGAGCAATGAGGACCCTACTGTTGCTGC | 906-1005 | 81 | 84 | XR_001740305.1;NM_001387873.1;NM_001146103.2;NM_001387884.1;NR_170865.1;NM_001146105.2;NM_001146102.2;XR_001740307.1;NM_001387876.1;NR_170863.1;NM_001146106.3;NM_001387871.1;NM_001387885.1;NM_001387880.1;NM_001387874.1;NM_001387886.1;NR_170857.1;NM_001387878.1;NM_001387875.1;NR_170858.1;NM_031458.3;XR_002959597.1;NR_170862.1;NM_001387877.1;NM_001387883.1;NR_170859.1;NM_001146104.2;NM_001387879.1;NR_170864.1;NM_001387887.1;NM_001387881.1;NR_170861.1;NM_001387872.1;XM_005247820.2;NR_170860.1;NM_001387882.1 |
| PDGFRB | CCCCTTCCTCCATCCCTCTGTTCTCCTGAGCCTTCAGGAGCCTGCACCAGTCCTGCCTGTCCTTCTACTCAGCTGTTACCCACTCTGGGACCAGCAGTCT | 266-365 | 85 | 85 | NM_001355016.2;NR_149150.2;NM_002609.4;NM_001355017.2 |
| PGK1 | GCAAGAAGTATGCTGAGGCTGTCACTCGGGCTAAGCAGATTGTGTGGAATGGTCCTGTGGGGGTATTTGAATGGGAAGCTTTTGCCCGGGGAACCAAAGC | 1031-1130 | 83 | 82 | NM_000291.4 |
| PIAS1 | AGTTAAGTGCAGGAGGCAGTACTTCTCTGCCAACCACCAATGGAAGCAGTAGTGGCAGTAACAGCAGCCTGGTTTCTTCCAACAGCCTAAGGGAAAGCCA | 1871-1970 | 81 | 83 | NM_016166.3;XM_011522127.2;XM_017022688.1;NM_001320687.1;XM_017022690.1;XM_017022691.2;XM_024450094.1;XM_011522126.2;XM_017022689.1 |
| POLR1B | GGAGAACTCGGCCTTAGAATACTTTGGTGAGATGTTAAAGGCTGCTGGCTACAATTTCTATGGCACCGAGAGGTTATATAGTGGCATCAGTGGGCTAGAA | 3321-3420 | 81 | 80 | NM_001282779.2;NM_019014.6;NM_001282772.2;NM_001371971.1;NM_001282774.2;NM_001282777.2;NM_001371970.1;NM_001137604.3;NM_001282776.2;NM_001371969.1 |
| POLR2a | TTCCAAGAAGCCAAAGACTCCTTCGCTTACTGTCTTCCTGTTGGGCCAGTCCGCTCGAGATGCTGAGAGAGCCAAGGATATTCTGTGCCGTCTGGAGCAT | 3776-3875 | 78 | 84 | NM_000937.5 |
| PPIA | TCTATGGGGAGAAATTTGAAGATGAGAACTTCATCCTAAAGCATACGGGTCCTGGCATCTTGTCCATGGCAAATGCTGGACCCAACACAAATGGTTCCCA | 316-415 | 82 | 85 | XM_024446809.1;XM_024446808.1;NM_021130.5;NM_001300981.2;XR_002956460.1  Also targets multiple PPIA-like genes (PPIAL4A, C, D, E, G) @ 95% |
| PPP2CA | GACATTTAATCATGCCAATGGCCTCACGTTGGTGTCTAGAGCTCACCAGCTAGTGATGGAGGGATATAACTGGTGCCATGACCGGAATGTAGTAACGATT | 1076-1175 | 81 | 81 | NM_001355019.2;NR_149151.2;NM_002715.4 |
| PPP2R1A | AACTTAACTCCTTGTGCATGGCCTGGCTTGTGGATCATGTATATGCCATCCGCGAGGCAGCCACCAGCAACCTGAAGAAGCTAGTGGAAAAGTTTGGGAA | 1441-1540 | 79 | 82 | NM_001363656.2;NM_014225.6;NR_033500.2 |
| PPP2R1B | GCTGAATTCTTTATGTATGGCTTGGCTCGTGGACCATGTATACGCCATCCGAGAAGCTGCCACCAACAACCTCATGAAACTAGTTCAGAAGTTTGGTACA | 1446-1545 | 80 | 84 | XM_024448598.1;NM_001177562.2;NM_181700.2;XM_017017960.2;NM_001177563.2;NM_181699.3;XM_017017961.1;XM_024448599.1;NM_002716.5;XM_024448600.1 |
| CD133 | AGCCTGCGGTCATCTCTCAATGACCCTCTGTGCTTGGTGCATCCATCAAGTGAAACCTGCAACAGCATCAGATTGTCTCTAAGCCAGCTGAATAGCAACC | 926-1025 | 83 | 79 | NM_001145850.2;NM_001145849.2;XM_017008800.1;NM_001145847.2;XM_011513897.3;XM_011513893.2;XM_006713974.3;XM_005248196.5;XM_005248195.5;NM_001371408.1;XM_011513896.2;XM_024454276.1;XM_011513895.2;NM_001145852.2;NM_001145848.2;XM_011513903.2;NM_001371406.1;XM_011513892.2;XM_011513900.2;NM_001371407.1;NM_001145851.2;NM_006017.3;XM_011513902.2;XM_011513894.3 |
| PTEN | TGTGGTCTGCCAGCTAAAGGTGAAGATATATTCCTCCAATTCAGGACCCACACGACGGGAAGACAAGTTCATGTACTTTGAGTTCCCTCAGCCGTTACCT | 1676-1775 | 79 | 78 | NM_001304717.5;NM_000314.8;NM_001304718.2  Also targets PTENP1 (NR_023917) @ 96% |
| PTK2 | GGTTCAAGCTGGATTATTTCAGTGGAACTGGCAATCGGCCCAGAAGAAGGAATCAGTTACCTAACGGACAAGGGCTGCAATCCCACACATCTTGCTGACT | 1006-1105 | 79 | 82 | XM_024447203.1;NM_001352708.2;NM_001352747.2;NM_001387585.1;NM_001387662.1;NM_001387607.1;NM_001352740.2;NM_001352723.2;NM_001387613.1;NM_001387642.1;NM_001387616.1;NM_001387641.1;NM_001387622.1;NM_001387645.1;NM_001352717.2;NM_001387660.1;NM_001387604.1;NM_001387646.1;NM_001387611.1;NM_001387630.1;NM_001199649.2;XM_024447202.1;NM_001352707.2;NM_001387584.1;NM_001352702.2;NM_001352712.2;NR_148038.2;NM_001352720.2;XM_024447201.1;NM_001387654.1;NM_001387627.1;NM_001387644.1;NM_001387636.1;NM_001387621.1;NM_001387638.1;NM_001352734.2;NM_001352703.2;NM_001352738.2;NM_001352704.2;NM_001387659.1;NM_001352695.2;NM_001387639.1;XM_024447206.1;NR_170671.1;NM_001352735.2;NM_001387605.1;NM_001387603.1;NM_001352741.2;NM_001352727.2;NM_001387657.1;NM_001387608.1;NM_001387649.1;NM_001352696.2;NM_001387612.1;XM_024447207.1;NM_001352721.2;NM_001352705.2;XM_017013656.2;NM_001387637.1;NM_001387634.1;NM_001387628.1;NM_001387620.1;NM_001387609.1;NM_001352730.2;NM_001387661.1;NM_001316342.2;NM_001352694.2;NM_001352706.2;XM_0170136 |
| RAC1 | AAAGACCTTCGTCTTTGAGAAGACGGTAGCTTCTGCAGTTAGGAGGTGCAGACACTTGCTCTCCTATGTAGTTCTCAGATGCGTAAAGCAGAACAGCCTC | 1251-1350 | 82 | 84 | NM_018890.4;NM_006908.5  Target accession removed from RefSeq - probe targets region common to all current variants of gene (2/2) |
| RELA | GATGGCTTCTATGAGGCTGAGCTCTGCCCGGACCGCTGCATCCACAGTTTCCAGAACCTGGGAATCCAGTGTGTGAAGAAGCGGGACCTGGAGCAGGCTA | 361-460 | 85 | 82 | NM_001243985.2;XM_011545207.2;NM_021975.4;NM_001145138.2;XM_011545206.2;NM_001243984.2 |
| RHOA | GGTACTCTGGTGAGTCACCACTTCAGGGCTTTACTCCGTAACAGATTTTGTTGGCATAGCTCTGGGGTGGGCAGTTTTTTGAAAATGGGCTCAACCAGAA | 1231-1330 | 79 | 84 | NM_001313947.2;NM_001313941.2;NM_001313946.2;NM_001664.4;NM_001313943.2;NM_001313945.2;NM_001313944.2 |
| RPL13A | AGTCCAGGTGCCACAGGCAGCCCTGGGACATAGGAAGCTGGGAGCAAGGAAAGGGTCTTAGTCACTGCCTCCCGAAGTTGCTTGAAAGCACTCGGAGAAT | 721-820 | 92 | 79 | NM_001270491.2;NR_073024.2;NM_012423.4 |
| RPLP0 | CGAAATGTTTCATTGTGGGAGCAGACAATGTGGGCTCCAAGCAGATGCAGCAGATCCGCATGTCCCTTCGCGGGAAGGCTGTGGTGCTGATGGGCAAGAA | 251-350 | 80 | 82 | NM_053275.4;NM_001002.4  Also targets pseudogene RPLP0P2 (NR_002775) @ 92% |
| RUNX2 | GAAGCCACAGCAGTTCCCCAACTGTTTTGAATTCTAGTGGCAGAATGGATGAATCTGTTTGGCGACCATATTGAAATTCCTCAGCAGTGGCCCAGTGGTA | 1851-1950 | 82 | 81 | NM_001369405.1;NM_001015051.4;NM_001278478.2;NM_001024630.4  Sub-maximal isoform coverage due to non-overlapping variants |
| SATB1 | TTCCGAAATCTACCAGTGGGTACGCGATGAACTGAAACGAGCAGGAATCTCCCAGGCGGTATTTGCACGTGTGGCTTTTAACAGAACTCAGGGCTTGCTT | 1336-1435 | 79 | 80 | XM_011533988.3;XM_011533989.2;XM_011533990.3;NM_002971.6;NM_001322874.2;NM_001322875.2;NM_001195470.3;NM_001322876.2;NM_001322872.2;NM_001322871.2;NM_001322873.2;NM_001131010.4 |
| SATB2 | TCGACACCGACAACAGACCTCCCTATTAAGGTGGACGGCGCCAACATCAACATCACAGCTGCCATTTATGACGAGATCCAACAGGAGATGAAAAGGGCCA | 1536-1635 | 82 | 82 | NM_001172517.1;XM_024452768.1;NM_015265.4;NM_001172509.2;XM_017003656.1;XM_024452767.1;XM_005246396.3;XM_011510840.3 |
| SHH | TCTGCACTACGAGGGCCGCGCAGTGGACATCACCACGTCTGACCGCGACCGCAGCAAGTACGGCATGCTGGCCCGCCTGGCGGTGGAGGCCGGCTTCGAC | 565-664 | 90 | 92 | NM_000193.4;NM_001310462.2;XM_011516480.2;NR_132318.2;NR_132319.2;XM_011516479.2 |
| SKP1 | CCCAGGTACGCAAAGAGAACCAGTGGTGTGAAGAGAAGTGAAATGTTGTGCCTGACACTGTAACACTGTAAGGATTGTTCCAAATACTAGTTGCACTGCT | 631-730 | 82 | 81 | NM_170679.3;NM_006930.4  Also targets SKP1P2 pseudogene (NR_036619) @ 95% |
| SLBP | ACTTGAAGCTCTCTTCCAGCGAGCTTAATTGCGTAATCCCTGTTGTCCTCCAGGGTAAGCTGACACGTCTACATAACTGGTTTTCCACAGGCATCTTCAG | 1155-1254 | 79 | 81 | XR_002959759.1;NM_006527.4;NM_001306074.2;NM_001306075.2 |
| SMAD5 | CCTGATGATCAGATGGGTCAAGATAATTCCCAGCCTATGGATACAAGCAATAATATGATTCCTCAGATTATGCCCAGTATATCCAGCAGGGATGTTCAGC | 1045-1144 | 84 | 79 | NM_005903.7;NM_001001419.3;XM_024446047.1;NM_001001420.3;XM_024446046.1;XM_017009470.2 |
| SMARCA4 | GCCCTGTCCTGGCATCAGTAGCATCTGTAACAGCATTAACTGTCTTAAAGAGAGAGAGAGAGAATTCCGAATTGGGGAACACACGATACCTGTTTTTCTT | 5401-5500 | 82 | 79 | NM_001128846.2;XM_024451667.1;XM_024451661.1;XM_024451658.1;XM_024451662.1;NR_164683.1;XM_024451659.1;XM_024451660.1;XM_006722846.2;NM_001374457.1;XM_024451663.1;XM_024451664.1;NM_001387283.1;NM_001128847.4;NM_001128844.3;NM_003072.5;XM_024451665.1;NM_001128845.2;NM_001128849.3;XM_011528198.1;NM_001128848.2 |
| SMARCA5 | AACGAGAAAGAAAAGCCAACTATGCCGTTGATGCATATTTCAGGGAAGCTCTTCGTGTTAGTGAACCTAAAGCACCCAAGGCTCCTCGACCTCCAAAACA | 2666-2765 | 83 | 83 | XR_001741338.1;NM_003601.4 |
| SMARCAL1 | TCCATCAGCTCCATCCCTTTCATTTGTCAAAGGGCGATGCATGCTCATCTCCAGGGCCTACTTCGAGGCAGACATCAGTTATTCACAGGACCTTATTGCG | 1296-1395 | 79 | 83 | NM_001127207.2;NM_014140.4 |
| SMARCB1 | AGAAGGAGAACTCACCAGAGAAGTTTGCCCTGAAGCTGTGCTCGGAGCTGGGGTTGGGCGGGGAGTTTGTCACCACCATCGCATACAGCATCCGGGGACA | 1061-1160 | 82 | 82 | NM_001362877.2;NM_003073.5;NM_001007468.3;NM_001317946.2 |
| SMARCC1 | CCAAACTACCTCTTGTTACAGTAGCCAGGGAGTGGAATTTCGTCAACCGGTACTTTTAAGGTTAGGATGGGACGGGAAAAGTGAAGCAGGATATTAGCTC | 4476-4575 | 82 | 82 | NM_003074.4 |
| SMO | AACGAGACCATGCTGCGCCTGGGCATTTTTGGCTTCCTGGCCTTTGGCTTTGTGCTCATTACCTTCAGCTGCCACTTCTACGACTTCTTCAACCAGGCTG | 1616-1715 | 82 | 81 | XM_024446891.1;NM_005631.5 |
| SNAI1 | GACCACTATGCCGCGCTCTTTCCTCGTCAGGAAGCCCTCCGACCCCAATCGGAAGCCTAACTACAGCGAGCTGCAGGACTCTAATCCAGAGTTTACCTTC | 64-163 | 85 | 86 | NM_005985.4 |
| SNAI2 | GCGTTTTCCAGACCCTGGTTGCTTCAAGGACACATTAGAACTCACACGGGGGAGAAGCCTTTTTCTTGCCCTCACTGCAACAGAGCATTTGCAGACAGGT | 741-840 | 80 | 81 | NM_003068.5 |
| SOX10 | GGCCGTGTCTCCCACTCAGGGGCTGAGAGTAGCTTTGAGGAGCCTCATTGGGGAGTGGGGGGTTCGAGGGACTTAGTGGAGTTCTCATCCCTTCAATGCC | 2160-2259 | 86 | 85 | NM_006941.4 |
| SSRP1 | CTTTGACTTTGAAATTGAGACCAAGCAGGGCACTCAGTATACCTTCAGCAGCATTGAGAGGGAGGAGTACGGGAAACTGTTTGATTTTGTCAACGCGAAA | 1431-1530 | 80 | 81 | XM_024448664.1;XM_024448665.1;NM_003146.3;XM_024448666.1;XM_017018180.1;XM_024448667.1 |
| SUPT16H | TGTGAAGATATGTGACGTGTATAACGCTGTCATGGACGTGGTTAAAAAGCAGAAGCCAGAACTGCTGAACAAAATTACCAAAAACCTAGGGTTTGGGATG | 1296-1395 | 74 | 74 | NM_007192.4;XM_011536381.2 |
| SUZ12 | TTGCAGTTCACTCTTCGTTGGACAGGAGAGACCAATGATAAATCTACGGCTCCTATTGCCAAACCTCTTGCCACTAGAAATTCAGAGAGTCTCCATCAGG | 1285-1384 | 78 | 77 | NM_015355.4;XM_017024409.1;XM_006721794.3;XR_001752462.1;NM_001321207.2 |
| TBP | ACAGTGAATCTTGGTTGTAAACTTGACCTAAAGACCATTGCACTTCGTGCCCGAAACGCCGAATATAATCCCAAGCGGTTTGCTGCGGTAATCATGAGGA | 588-687 | 79 | 82 | NM_001172085.2;NM_003194.5 |
| TET1 | AATGCCAATCAGAAAGCCCATCCTTTGACCCAGCCCTCCTCTCCACCTAACCAGTGTGCTAACGTGATGGCAGGCGATGACCAAATACGGTTTCAGCAGG | 4376-4475 | 81 | 82 | NM_030625.3;XM_011540206.2;XR_001747212.1;XR_001747210.1;XM_017016688.1;XR_001747211.1;XM_017016689.1;XM_011540204.2;XM_017016686.2;XM_011540207.2;XM_017016687.1;XM_011540205.2 |
| TET2 | CTCATAATGTCCAAATGGGACTGGAGGAAGTACAGAATATAAATCGTAGAAATTCCCCTTATAGTCAGACCATGAAATCAAGTGCATGCAAAATACAGGT | 2883-2982 | 81 | 79 | XR_427546.4;NM_001127208.3;XR_001741246.1;XM_024454102.1;XM_006714242.3;XR_938747.3;NM_017628.4;XR_244633.3;XR_938746.2;XM_005263082.3;XM_017008319.1;XM_024454103.1 |
| TET3 | AAGCCATCCGGATCGAGAAGGTCATCTACACGGGGAAGGAGGGAAAGAGCTCCCGCGGTTGCCCCATTGCAAAGTGGGTGATCCGCAGGCACACGCTGGA | 2198-2297 | 92 | 91 | NM_001287491.2;XM_011532682.2;XM_011532686.2;XM_017003566.1;XM_011532684.2;NM_001366022.1;XM_011532690.1;XM_011532688.2;XM_011532683.2;XM_024452747.1;XM_024452745.1;XM_024452746.1;XM_011532687.2;XM_011532685.2 |
| TFRC | CAGTTTCCACCATCTCGGTCATCAGGATTGCCTAATATACCTGTCCAGACAATCTCCAGAGCTGCTGCAGAAAAGCTGTTTGGGAATATGGAAGGAGACT | 1221-1320 | 79 | 81 | NM_003234.4;XR_002959577.1;XR_002959578.1;XM_024453732.1;XM_024453731.1;NM_001313965.2;XR_002959575.1;NM_001128148.3;NM_001313966.2;XR_002959576.1 |
| TIPARP | CCAATACATTCTGGACACCAGTGATAAGCTGAGTACTGAGCTCTTTCAGGACAAAAGTGAAGAGGCTTCCCTTGACCTCGTGTTTGAGCTGGTGAACCAG | 836-935 | 80 | 79 | NM_015508.5;NM_001184717.1;NM_001184718.2 |
| TJP1 | CACATTTTTCTTAGGGAAGGATACAAAAGCATGTGAGACTGGTTCCATGGCCTCTTCAGATCTCTAACTTCACCATATTACCACAGACATACTAACCAGC | 6278-6377 | 78 | 80 | NM_175610.4;XM_017022524.1;NM_001355015.2;XM_005254620.3;XM_005254619.3;XM_017022531.1;NM_001301025.3;NM_001301026.2;NM_001355014.2;XM_017022522.1;XM_017022527.1;XM_017022526.1;XM_017022525.1;XM_017022521.1;XM_017022523.1;NM_001355012.2;NM_001330239.4;XM_011521972.2;NM_001355013.1;NM_003257.5 |
| TNKS | AAAGCATGGAGCTTGTGTTAATGCCATGGATCTCTGGCAGTTTACTCCACTGCACGAGGCTGCTTCCAAGAACCGTGTAGAAGTCTGCTCTTTGTTACTT | 1271-1370 | 82 | 81 | XM_011543845.3;XM_006716263.4;XM_011543846.3;NM_003747.3 |
| TNKS2 | CAAACCTGCCGCCTACTCCTGAGCTATGGGTGTGATCCTAACATTATATCCCTTCAGGGCTTTACTGCTTTACAGATGGGAAATGAAAATGTACAGCAAC | 1636-1735 | 84 | 81 | NM_025235.4;XM_005270185.4;XM_017016700.2;XM_017016699.1;XM_011540213.1;XM_011540215.2;XM_017016698.2;XM_017016697.1;XM_017016701.1;XM_017016696.1 |
| TNRC6A | GTTGAGTGGAACAAACTGCCTAGCAATCAGCATTCCAATGATAGTGCAAATGGCAATGGTAAGACGTTTACAAATGGATGGAAATCTACTGAGGAAGAGG | 1957-2056 | 82 | 83 | XM_024450233.1;XM_017023152.2;NM_001330520.3;XM_017023145.2;XM_017023154.1;XM_017023144.2;XM_017023153.1;XM_024450232.1;XM_017023150.2;XM_005255257.4;XM_017023146.1;NM_001351850.2;NM_014494.4;XM_017023148.2;XM_024450231.1 |
| TUBB | TTCTAAGTATGTCCATTTCCCATCTCAGCTTCAAGGGAGGTGTCAGCAGTATTATCTCCACTTTCAATCTCCCTCCAAGCTCTACTCTGGAGGAGTCTGT | 1956-2055 | 79 | 81 | NM_001293213.2;NM_001293214.2;NM_001293215.2;NR_120608.2;NM_001293212.2;NM_001293216.2;NM_178014.4 |
| TWIST1 | CAACTCCCAGACACCTCGCGGGCTCTGCAGCACCGGCACCGTTTCCAGGAGGCCTGGCGGGGTGTGCGTCCAGCCGTTGGGCGCTTTCTTTTTGGACCTC | 36-135 | 87 | 82 | NM_000474.4;NR_149001.2 |
| VEGFA | GAGTCCAACATCACCATGCAGATTATGCGGATCAAACCTCACCAAGGCCAGCACATAGGAGAGATGAGCTTCCTACAGCACAACAAATGTGAATGCAGAC | 1326-1425 | 82 | 82 | NM_001171630.2;NM_001204384.2;NM_001287044.2;NM_001171627.2;NM_001025370.3;NM_001171628.2;NM_003376.6;NM_001171622.2;NM_001317010.1;NM_001171624.2;NM_001025368.3;NM_001171623.2;NM_001025366.3;NM_001025369.3;NM_001025367.3;NM_001204385.2;NM_001171626.2;NM_001033756.3;NM_001171625.2;NM_001171629.2 |
| VEGFC | GGCGAGGCCACGGCTTATGCAAGCAAAGATCTGGAGGAGCAGTTACGGTCTGTGTCCAGTGTAGATGAACTCATGACTGTACTCTACCCAGAATATTGGA | 566-665 | 81 | 79 | NM_005429.5 |
| VIM | GAGGAGATGCTTCAGAGAGAGGAAGCCGAAAACACCCTGCAATCTTTCAGACAGGATGTTGACAATGCGTCTCTGGCACGTCTTGACCTTGAACGCAAAG | 695-794 | 81 | 79 | NM_003380.5;XM_006717500.2 |
| XPO1 | AATCTTTCTTCAGGAATATGTGGCTAATCTCCTTAAGTCGGCCTTCCCTCACCTACAAGATGCTCAAGTAAAGCTCTTTGTGACAGGGCTTTTCAGCTTA | 3641-3740 | 72 | 75 | XM_024453126.1;XM_005264546.2;XM_011533099.3;NM_003400.4;XM_011533098.2;XM_011533097.1;XM_005264544.2;XM_024453127.1;XM_006712094.3;XM_024453125.1 |
| ZEB1 | TTACAAAATGGGGTTTTCACTGGTGGTGGCCCATTACAGGCAACCAGTTCTCCTCAGGGCATGGTGCAAGCTGTTGTTCTGCCAACAGTTGGTTTGGTGT | 1451-1550 | 82 | 83 | NM_001323665.2;NM_001323664.2;NM_001174094.2;XM_006717498.2;NM_001323673.2;NM_001323662.2;NM_001323674.2;NM_001323657.2;NM_001323649.2;NM_001323642.2;NM_001323675.2;NM_001323654.2;NM_001323672.2;XM_017016597.1;NM_001323648.2;NM_001323645.2;NM_001323638.2;NM_001323651.2;XM_011519643.2;NM_001323666.2;NM_001323656.2;NM_001323678.2;NM_001323671.2;NM_001174093.2;NM_001323661.2;NM_001323643.2;NM_001323653.2;NM_001323659.2;NM_001323658.2;NM_001323652.2;NM_001323677.2;NM_001323644.2;NM_001323647.2;NM_030751.6;NM_001174095.2;NM_001128128.3;NM_001174096.2;NM_001323676.2;NM_001323655.2;NM_001323663.2;NM_001323646.2;NM_001323641.2;NM_001323660.2;NM_001323650.2 |
| ZEB2 | CAAGACTTCGCAGATCGAGCCTGCGTGCTGCCGAAGCAGGGCGCCGAGTCCATGCGAACTGCCATCTGATCCGCTCTTATCAATGAAGCAGCCGATCATG | 441-540 | 84 | 85 | NM_014795.4;NM_001171653.2;NR_033258.2 |
